# A DNA-based synthetic apoptosome

**DOI:** 10.1101/660183

**Authors:** Bas J.H.M. Rosier, Albert J. Markvoort, Berta Gumí-Audenis, Job A.L. Roodhuizen, Anniek den Hamer, Luc Brunsveld, Tom F.A. de Greef

## Abstract

Living cells are able to regulate key cellular processes by physically assembling signaling components on dedicated molecular platforms. The spatial organization of proteins in these higher-order signaling complexes facilitates proximity-driven activation and inhibition events, allowing tight regulation of the flow of information. Here, we employ the programmability and modularity of DNA origami as a controllable molecular platform for studying protein-protein interactions involved in intracellular signaling. Specifically, we engineer a synthetic, DNA origami-based version of the apoptosome, a large multi-protein signaling complex that regulates apoptosis by co-localization of multiple caspase-9 monomers. Our *in vitro* characterization using both wildtype caspase-9 monomers and inactive mutants tethered to a DNA origami platform directly demonstrates that enzymatic activity is induced by proximity-driven dimerization with asymmetric, half-of-sites reactivity. Additionally, experimental results supported by a detailed thermodynamic model reveal a multivalent activity enhancement in tethered caspase-9 oligomers of three and four enzymes, partly originating from a statistical increase in the number of active catalytic units in higher-order enzyme clusters. Our results offer fundamental insights in caspase-9 activity regulation and demonstrate that DNA origami provides a modular platform to construct and characterize higher-order signaling complexes. The engineered DNA-based protein assembly platform has the potential to be broadly applied to inform the function of other important multi-enzyme assemblies involved in inflammation, innate immunity, and necrosis.

## Introduction

Nanoscale organization of interacting proteins is a key regulatory principle in signaling pathways involved in all major cell events, including apoptosis, metabolism, inflammation, and immunity^1,2^. Inactive enzymes with a low intrinsic affinity and present at low intracellular concentrations, can be physically assembled into well-defined higher-order signaling complexes^3^ or open-ended assemblies^4^. In these processes, dedicated scaffold proteins serve as supramolecular organizing centers (SMOCs), facilitating protein-protein interactions with precise control over the position and orientation of the individual components (Fig. 1a)^5^. Efforts to address and rewire SMOC-based signaling complexes have provided important structural and functional understanding into the underlying design principles^6–8^. In general, experimental and theoretical work illustrate that co-localization of signaling components promotes proximity-induced enzyme activation through weak non-covalent interactions, thereby overcoming signal thresholds, increasing pathway robustness, and shaping response dynamics^9–11^.

**Figure 1.**
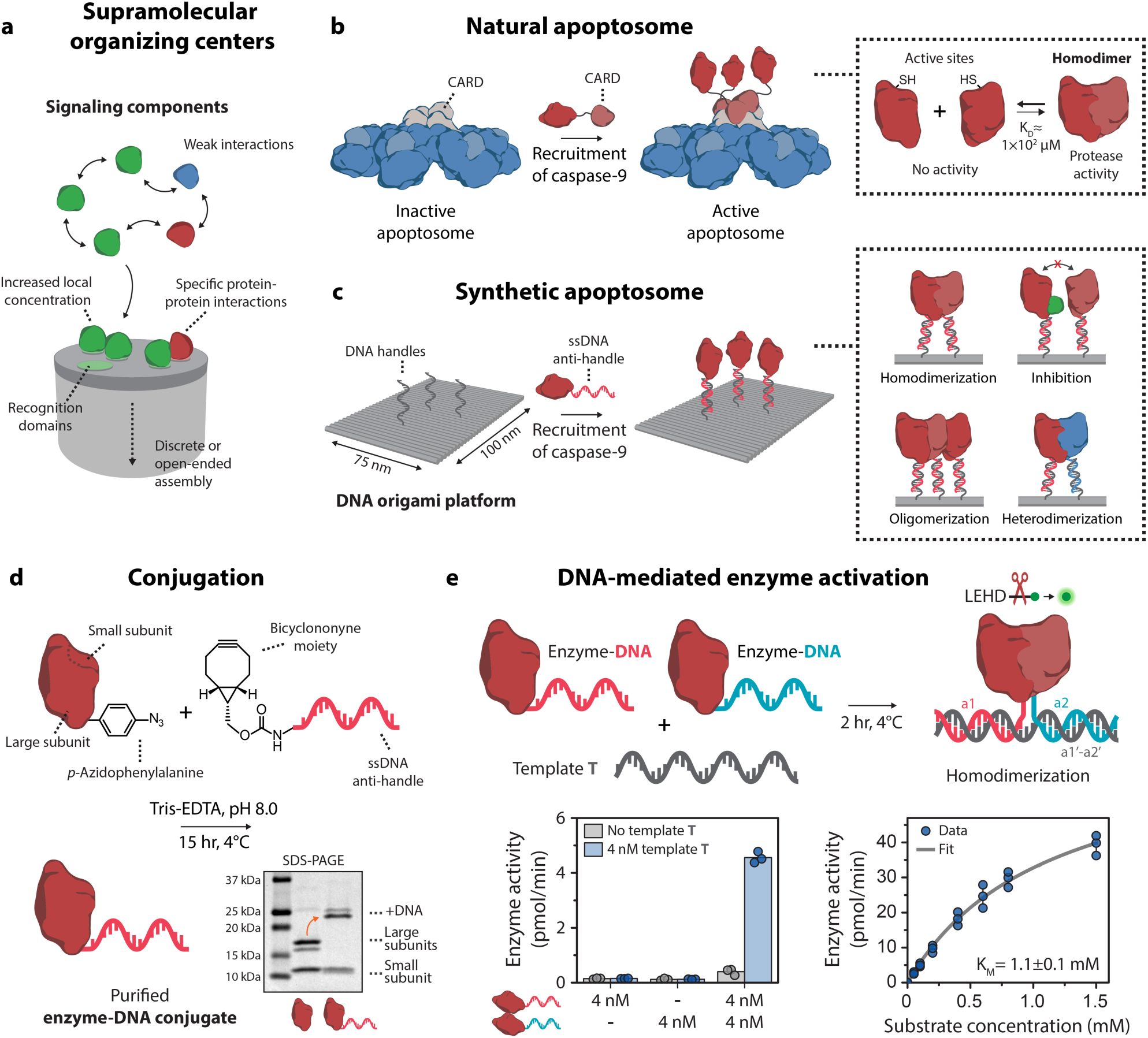
General concept and design elements for the construction of a DNA-based synthetic apoptosome. **a,** Schematic concept of supramolecular organizing centers (SMOCs). Signaling components are recruited to discrete or open-ended assembly platforms resulting in proximity-induced protein-protein interactions. **b**, Schematic drawing of the natural apoptosome that functions by assembling inactive caspase-9 monomers through caspase recruitment domains (CARDs). The increase in local concentration induces caspase-9 dimerization, leading to proteolytic cleavage of downstream caspases and eventually apoptosis. **c**, Schematic drawing of the DNA origami-based synthetic apoptosome, which uses single-stranded DNA (ssDNA) handles to control the number, position, and relative geometry of caspase-9 monomers. The programmability and modularity of DNA origami allows for the characterization of specific protein-protein interactions, such as homodimerization, inhibition, oligomerization, and heterodimerization. **d**, Reaction scheme for the conjugation of caspase-9 to an oligonucleotide anti-handle. Caspase-9 was expressed in *E. coli* with unnatural amino acid *p*-azidophenylalanine at the N-terminus and, due to autocatalytic processing, consists of N-terminal large (18-19 kDa) and C-terminal small subunits (13 kDa). Overnight reaction of caspase-9 to a bicyclononyne-functionalized oligonucleotide (8 kDa) using strain-promoted azide-alkyne cycloaddition and subsequent purification with ion-exchange chromatography and affinity chromatography afforded pure enzyme-DNA conjugates, as shown by 4-20% SDS-PAGE analysis. **e**, To verify the activity of caspase-9 enzyme-DNA conjugates, bivalent template **T** was used to induce dimerization. Enzyme kinetics were determined by measuring protease activity at varying concentrations of synthetic tetrapeptide substrate LEHD-AFC and the data was fitted with the standard Michaelis-Menten expression. Bars represent mean enzyme activity. All experiments were performed in independent triplicates.

Bottom-up approaches employing synthetic platforms enable systematic analysis and full control over the number, position, and orientation of interacting components, providing an excellent strategy to further unravel the molecular mechanisms behind spatial organization in signaling pathways^12,13^. The programmability of DNA and its inherent biocompatibility enables rational design of defined synthetic architectures for the construction of protein-DNA hybrid systems^14–17^. DNA origami-based nanostructures, in particular, are well-suited as synthetic platforms as their unique addressability allows for precise assembly of multiple non-identical proteins with nanometer accuracy^18–20^. The DNA origami technique has found broad applicability as an experimental tool for spatial organization of native multi-protein systems, such as amyloid fibrils^21^, membrane fusion proteins^22^, nucleosomes^23,24^, and intrinsically disordered proteins^25^. Additionally, these platforms have been used to engineer localized genetic circuits^26^, to study confinement-induced enzyme activity^27^, and to investigate scaffolded metabolic cascades^28–31^. Although these studies have elegantly applied the programmability of DNA nanotechnology to facilitate organization of e.g. structural protein assemblies and metabolic enzymes with small-molecule substrates, a DNA nanostructure-based platform for directly probing-protein interactions between catalytically active intracellular signaling components is currently lacking.

Here, we present the first DNA-based synthetic SMOC for studying proximity-induced protein-protein interactions involved in intracellular signal transduction. Specifically, we construct DNA origami-based synthetic variants of the apoptosome, a 27-nm diameter, sevenfold-symmetric multi-protein complex involved in the intrinsic apoptotic pathway. In the cell, mitochondrial outer membrane permeabilization and subsequent release of cytochrome c induces the assembly of the apoptosome, which recruits the <u>c</u>ysteine-dependent <u>asp</u>artic prote<u>ase</u> caspase-9 to initiate a cascade of proteolytic activity that eventually leads to programmed cell death^32^. Previous works suggest that the apoptosome recruits up to four caspase-9 monomers through caspase recruitment domains (CARDs), after which dimerization contributes to a dramatic increase in enzyme activity (Fig. 1b)^33–35^. By mimicking the scaffolding function of the apoptosome with a rectangular DNA origami platform^18^, we can assemble individual caspase-9 monomers with absolute control over their position, using the hybridization of DNA-functionalized enzymes to protruding single-stranded handles on the DNA origami surface (Fig. 1c). Using a bottom-up approach, our *in vitro* studies reveal that caspase-9 activity is induced by proximity-driven dimerization, driven by an increase in effective concentration as a result of tethering of the components to the platform. Exploiting the modularity of DNA origami, we then construct three- and four-enzyme systems and study the effect of higher-order clustering on caspase-9 activity. By combining experimental and theoretical results, we analyze kinetic data considering possible effects such as enzyme incorporation efficiency and statistical factors, and suggest a multivalent catalytic effect leading to enhanced activity in caspase-9 oligomers. Finally, we provide direct evidence that a caspase-9 heterodimer, consisting of a wildtype monomer and an active-site mutant has equal activity compared to the wildtype homodimer, confirming the hypothesis that conformational changes in the active sites of a caspase homodimer proceed via an asymmetric mechanism^36^. Our experimental platform demonstrates that systematic *in vitro* analysis of native protein-protein interactions using DNA-based synthetic SMOCs can facilitate the discovery of new molecular mechanisms in proximity-driven enzyme regulation.

## Results and discussion

### Activity of DNA-functionalized caspase-9

To construct a DNA origami-based platform that allows for programmable caspase-9 organization, well-defined enzyme-DNA conjugates were synthesized^20,37^. Full-length caspase-9 monomers consist of an N-terminal caspase recruitment domain (CARD) that interacts with the apoptosome, and a catalytic domain responsible for its protease activity^36^. Replacing the CARD with an oligonucleotide allows incorporation of the catalytic domain onto DNA nanostructures, and mimics the recruiting function of the CARD using hybridization to complementary single-stranded handles on the DNA origami surface (Fig. 1b,c). The specific protein-protein interactions involved in caspase-9 activation warrant stoichiometric, site-specific oligonucleotide functionalization without the use of additional, large protein helper domains. Therefore, we incorporated the non-canonical amino acid *p*-azidophenylalanine at the N-terminus of the catalytic domain of caspase-9 by using amber codon suppression in *E. coli* with an engineered orthogonal amino acyl tRNAse/tRNA pair from *M. janaschii* (see Methods and Supplementary Fig. 1 and 2)^38^. The small bioorthogonal azide moiety was used for conjugation to a bicyclononyne-functionalized oligonucleotide (BCN-DNA) using strain-promoted azide-alkyne cycloaddition (Fig. 1d)^39–41^. The oligonucleotide was designed with a 10-nucleotide (nt) single-stranded linker separating the enzyme and a 15-nt anti-handle used for hybridization to the handles on the DNA origami surface (Supplementary Table 2 and Supplementary Fig. 3). Analysis using polyacrylamide gel electrophoresis confirmed successful conjugation of a single oligonucleotide to the N-terminus of caspase-9 and subsequent purification resulting in complete removal of all unreacted protein and excess BCN-DNA (Fig. 1d, Supplementary Fig. 4 and 6).

Since conjugation of oligonucleotides to enzymes can significantly influence catalytic behavior^42,43^, we measured the activity of the caspase-9 enzyme-DNA conjugates. To this end, we employed a 30-nt single-stranded template **T** to bring two caspase-9 monomers into close proximity through DNA hybridization (Fig. 1e and Methods for design)^44^. Following proteolytic cleavage of synthetic caspase substrate LEHD-AFC^45^ over time, we observed a sharp increase in activity only when both enzyme-DNA conjugates and the template are present (Fig. 1e; left graph). This suggests that template **T** functions as a bivalent scaffold inducing dimerization of caspase-9 monomers by increasing the effective concentration, in a similar manner as protein-based dimerizing scaffolds reported in literature^46–48^. We performed a quantitative kinetic analysis and determined the Michaelis constant *K*_M_ of the ternary complex consisting of the caspase-9 conjugates and template **T**. The value of 1.1 ± 0.1 mM is in the same range as found for caspase-9 activation by the native apoptosome and by other, synthetic scaffold platforms (Fig. 1e; right graph)^45^. Collectively, these results confirm the successful synthesis of functionally active caspase-9 enzyme-DNA conjugates, and demonstrate that DNA can be used to facilitate proximity-induced enzyme activation.

### DNA origami-mediated caspase-9 enzyme activity

The native apoptosome is an organizing platform with seven enzyme recruitment domains, typically binding up to four caspase-9 monomers simultaneously, but so far, most cellular and synthetic biology approaches have only investigated caspase-9 activation by employing bivalent scaffolds^33–35^. To assess the effects of multivalent enzyme clustering, we constructed a synthetic version of the apoptosome using a rectangular DNA origami nanostructure^18^, on which the location of an arbitrary number of enzymes can be tightly controlled by hybridization to single-stranded handles on the DNA origami surface (see Supplementary Information for design). The unique addressability of DNA origami allows for programmable positioning of the handles with ∼6 nm resolution across the entire platform (Fig. 2a). Atomic force microscopy (AFM) confirmed the correct self-assembly of 75×100-nm^2^ DNA origami nanostructures and revealed that individual 4.5-nm diameter, 32-kDa caspase-9 monomers can be clearly observed on the DNA origami surface (Fig. 2b and Supplementary Fig. 10). Analysis of 260 well-formed one-enzyme DNA nanostructures revealed an average enzyme incorporation efficiency of 76% (Supplementary Fig. 13), which is similar to values reported in literature^49,50^.

**Figure 2.**
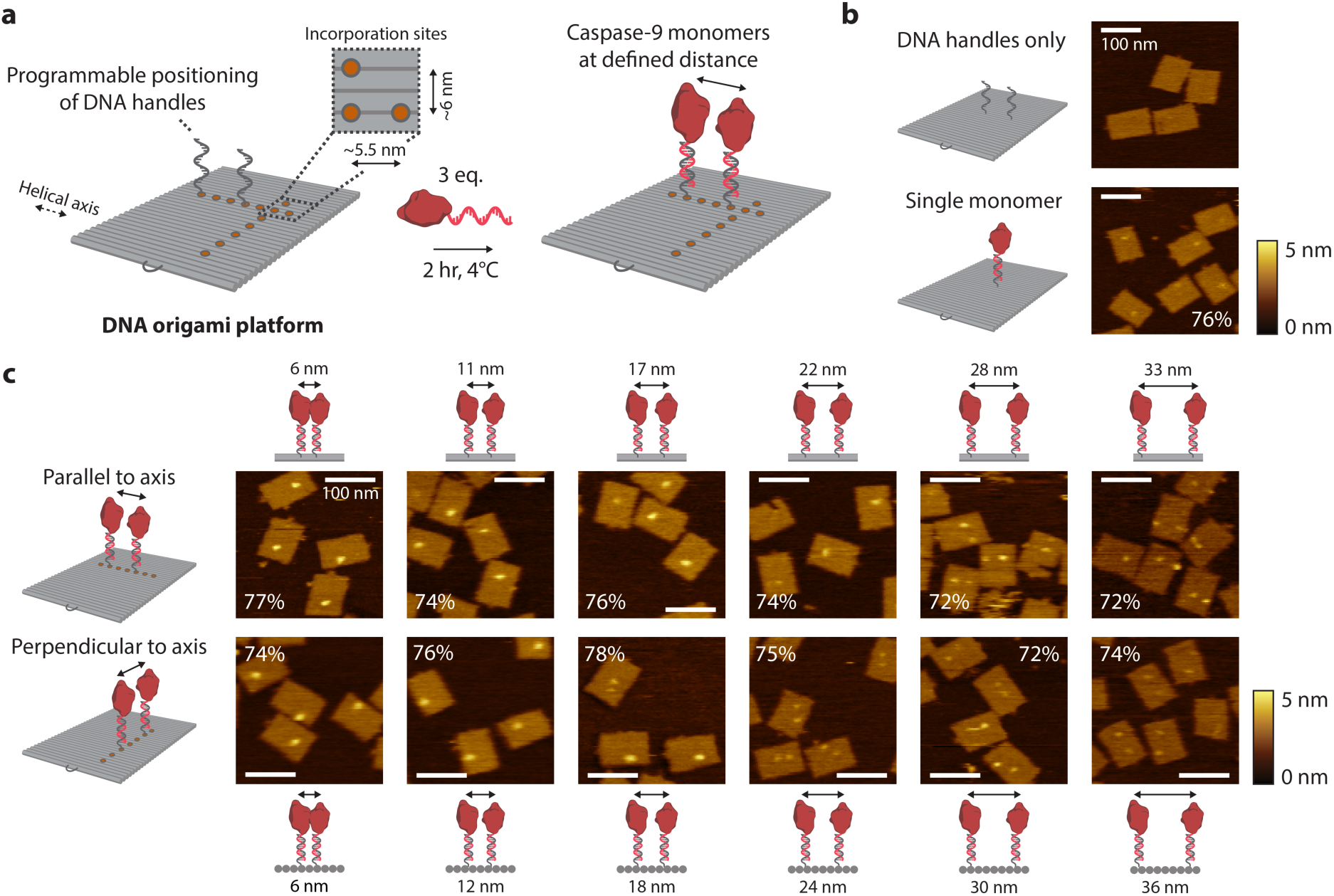
Characterization of caspase-9 assembly onto DNA origami nanostructures. **a,** Schematic of the general strategy for caspase-9 incorporation on DNA origami platforms. By including appropriate handle-extended staple strands during the self-assembly process the position of, and distance between, two ssDNA handles can be controlled. The minimal distance between incorporation sites (orange circles) is 5.5 nm parallel, and 6 nm perpendicular to the DNA helical axis. Incubation of complementary enzyme-DNA conjugates leads to hybridization and incorporation of two caspase-9 monomers at defined distances. Typically, 4 nM DNA origami was incubated with 3 equivalents of enzyme-DNA conjugate per handle for 2 hr at 4°C. For AFM imaging, functionalized nanostructures were purified using 1.5% agarose gel extraction. **b,c,** Topographic AFM (tapping mode in solution) images of control samples (**b**) and DNA origami nanostructures functionalized with two 32-kDa caspase-9 monomers (**c**) at various distances parallel (top row) and perpendicular (bottom row) to the helical axis. The caspase-9 incorporation efficiency per handle is indicated in percentages, and was calculated based on at least 250 well-formed nanostructures using 4 different images per sample. Color bars indicate height scale in AFM images. Scale bars, 100 nm.

Next, we designed two-enzyme DNA nanostructures with varying distance between two caspase-9 monomers arranged either parallel (Fig. 2c; top row) or perpendicular (bottom row) to the DNA helical axis, and used both agarose gel electrophoresis (Supplementary Fig. 9) and AFM imaging (Fig. 2c, Supplementary Fig. 11 and 12) to analyze the integrity of the structures. When monomer separation is large (i.e. >20 nm), two individual spots can be distinguished in the AFM images, indicating faithful incorporation of caspase-9 at the pre-programmed positions, as well as functional separation of the monomers by the DNA nanostructure. In contrast, only a single spot is discerned at smaller separation distances (<20 nm), with the shape and intensity of these features suggesting the presence of two adjacent caspase-9 monomers (Supplementary Fig. 13). We attribute this observation to a combination of the limited resolution of the imaging technique and intermolecular interactions (both specific and non-specific, see below) between the enzymes. Visual inspection of the AFM images allows straightforward identification of DNA nanostructures with either one or two enzymes (Fig. 2c; compare e.g. bottom row, 12 and 24 nm). The enzyme incorporation efficiency per handle was found to be approximately 75% per handle for all samples, irrespective of monomer separation (Fig. 2c, Supplementary Fig. 13). Although this is in contrast with previous work that reported a diminished incorporation when two enzymes were brought into close proximity on a similar DNA origami scaffold^29^, we hypothesize that the attractive interaction between two caspase-9 monomers balances possible steric effects at small separation distances, leading to an overall constant incorporation efficiency. Taken together, the results of the AFM analysis confirm that DNA nanostructures containing two caspase-9 monomers can be constructed with tight control over the position of, and distance between, tethered monomers.

We then assessed if caspase-9 displays functional enzymatic activity when assembled on DNA origami nanostructures. Since proximity-induced activation of caspase-9 involves dimerization, we envisioned that the activity can be tuned by varying the distance between the monomers. To this end, we assembled two-enzyme DNA nanostructures with monomer separation varying between 6 and 36 nm, and followed proteolytic activity over time (Fig. 3a,b and Methods). Highest enzyme activity was observed when the monomers were in closest proximity, after which the activity dropped sharply and approached background activity levels for monomer separations >20 nm (Fig. 3b). The arrangement of caspase-9 monomers either parallel or perpendicular to the DNA helical axis did not influence the distance-dependent enzymatic activity (Fig. 3b; compare top and bottom graphs). Increasing the single-stranded DNA linker between enzyme and handle from 10 to 15 nt slightly decreased the maximum activity while retaining distance-dependent behavior, illustrating that enzyme activation requires tight co-localization of the monomers and that the DNA-based assembly method does not introduce adverse steric effects (Supplementary Fig. 15). After correcting for background activity and incorporation efficiency as determined by AFM, DNA origami-mediated caspase-9 activation at 6-nm monomer separation resulted in a 23-fold increase in enzyme activation, equivalent to caspase-9 dimerization enforced by the bivalent template **T** (Fig. 3c).

**Figure 3.**
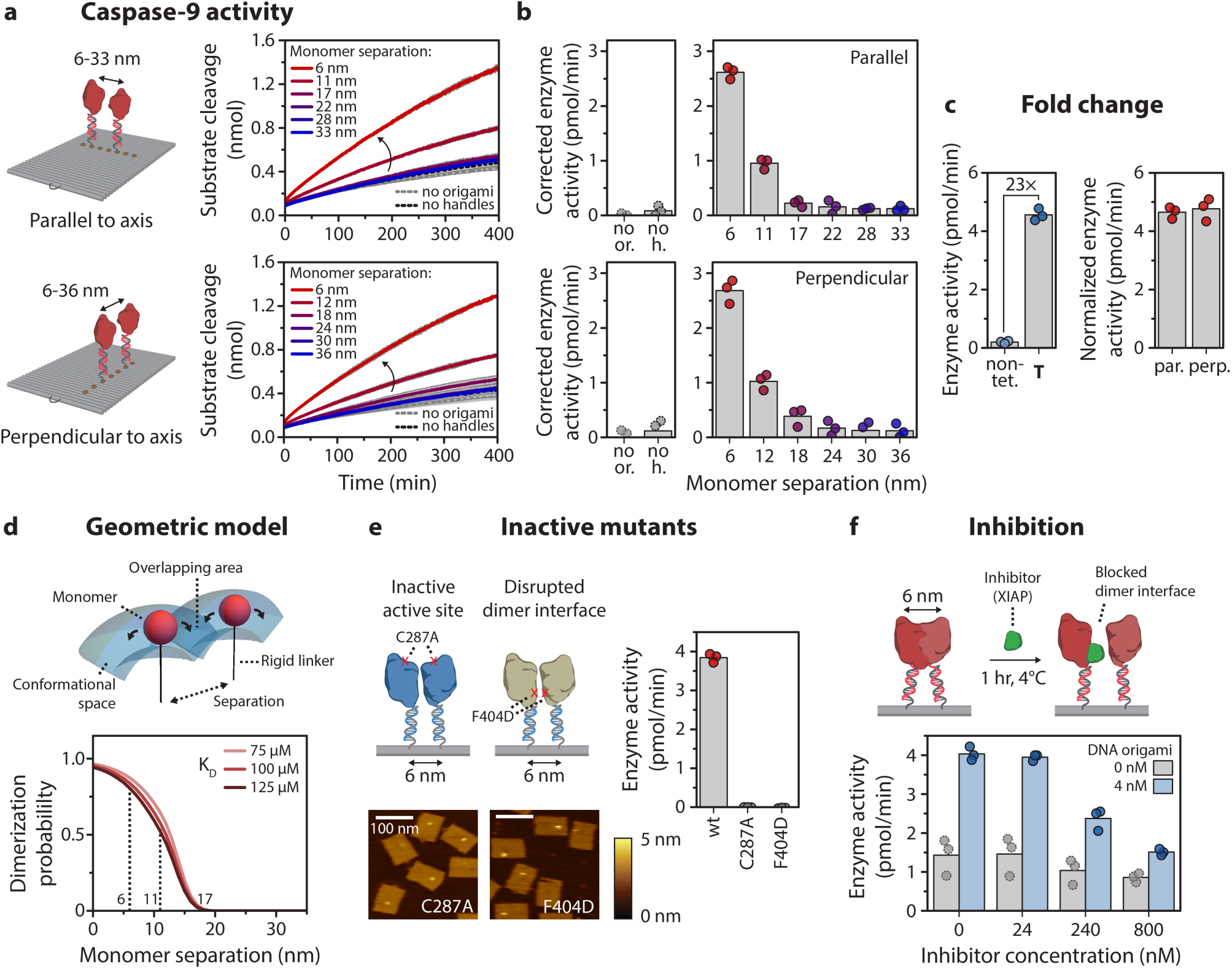
Activation of caspase-9 occurs by distance-dependent dimerization of tethered monomers. Reactions were carried out with 4 nM DNA origami (unless indicated otherwise) and 3 equivalents of enzyme-DNA conjugate per handle, and were incubated for 2 hr at 4°C. Activity was determined by monitoring cleavage of 167 µM LEHD-AFC caspase substrate at 18°C. Bars represent mean activity. All experiments were performed in independent triplicates. **a,b,** Distance-dependent enzyme activity of two caspase-9 monomers on DNA origami nanostructures parallel (top) and perpendicular (bottom) to the DNA helical axis. Data in **a** is represented as mean ± s.d. of three independent experiments. Enzyme activity (**b**) was determined by taking the initial slope of the kinetic time traces in **a** and corrected by subtracting the mean background activity (no or.). Labels: no or., no DNA origami present; no h., DNA origami without handles for enzyme incorporation. **c**, Fold change in enzyme activity for template **T**, and 6-nm samples parallel (par.) and perpendicular (perp.) to DNA origami helical axis was calculated by comparing with the activity of 4 nM non-tethered caspase-9 in buffer (non-tet.). The activity of DNA origami samples was normalized based on an incorporation efficiency per handle of 75%. **d**, Three-dimensional geometric model assuming free movement for each tethered monomer in the conformational space, determined by molecular dynamics simulations. Dimerization can only occur when both monomers are in the overlapping area between both conformational spaces. The tethered dimerization probability was plotted as a function of the K_D_ of caspase-9 dimerization in solution and the separation between the tethered monomers. **e**, Expression, conjugation, and assembly of caspase-9 point mutants C287A and F404D on DNA origami nanostructures at 6 nm monomer separation was performed similarly to wildtype (wt) caspase-9. AFM images show correct incorporation of enzymes, but both mutants exhibit no enzymatic activity. Color bars indicate height scale in AFM images. Scale bars, 100 nm. **f**, Inhibition of caspase-9 by the BIR3 domain of X-linked inhibitor of apoptosis protein (XIAP), which binds to the N-terminus of the small subunit of a caspase-9 monomer (*K*_i_ < 20 nM), blocking the dimer interface and preventing dimerization.

To validate DNA origami nanostructures as inert assembly platforms we performed several essential control experiments. First, a control in which DNA origami without handles was used exhibited only background activity, confirming that the DNA nanostructures do not influence caspase-9 function (Fig. 3b; compare dotted gray and black). Next, we assessed the influence of an altered pH near the surface of the negatively-charged DNA origami structure^51^, by measuring enzyme activity both in solution and mediated by the 6-nm two-enzyme DNA nanostructure at varying pH levels. Both experiments revealed a bell-shaped pH dependence with an optimum at pH 7.0^52^, suggesting that the behavior of tethered caspase-9 near the surface of the DNA origami platform is not affected by local changes in pH (Supplementary Fig. 16). Finally, we determined the kinetic parameters of the 6-nm two-enzyme DNA nanostructure and found a *K*_M_ of 1.8 ± 0.1 mM (Supplementary Fig. 17). Although this is slightly higher than the value reported for the bivalent template **T** (Fig. 1e), it is known that immobilization of enzymes on a surface can result in an increase in *K*_M_, which can be attributed to a lower affinity of the substrate for the enzyme due to diffusional limitations or conformational changes of the enzyme near the surface^53^. Collectively, these results establish DNA origami nanostructures as an ideal platform for the assembly of physically interacting enzymes, such as caspase-9, and the systematic analysis of the effects of relative geometry on enzyme activity.

To further rationalize the experimental results, we constructed a geometric model for two tethered interacting enzymes based on the concept of effective concentration^54^ and calculated the fraction of tethered dimer as a function of the distance between the anchor points (Fig. 3d). The model was constructed by estimating the dimensions of the system, including the size of a caspase-9 monomer, the length of the 15-nt double-stranded handle-anti-handle tether, and the length of linkers between the individual components (Supplementary Fig. 18). Coarse-grained molecular dynamics simulations were employed to define a conformational volume in which the tethered enzymes can move freely (Fig. 3d and Supplementary Fig. 18). The model allowed us to calculate the dimerization probability as a function of monomer separation and the *K*_D_ of non-tethered caspase-9 (see Methods)^55^. The results clearly show that at monomer separation >20 nm, the calculated fraction of dimer approaches zero as the tethered monomers cannot physically interact (Fig. 3d). In accordance with the experimental data, a sharp drop in the dimerization probability is observed in the regime between 5 and 15 nm, decreasing from 90% at 5 nm to less than 10% at 15 nm. Although the model does not consider steric effects or the specific mutual orientation of the monomers, it describes the experimental data particularly well, suggesting that caspase-9 dimerization on the DNA origami platform originates from an increase in effective concentration. Taken together, these results confirm that DNA origami-mediated activation of caspase-9 is consistent with proximity-induced homodimerization, and that the extent of activity can be tuned by varying the separation between interacting monomers.

To illustrate the functionality of the DNA origami platforms for studying protein-protein interactions, we investigated the behavior of caspase-9 mutants and the effect of inhibition on enzyme activity using a biologically relevant inhibitor. First, we expressed and conjugated two caspase-9 point mutants to DNA (Supplementary Fig. 1 and 5), resulting in enzyme-DNA conjugate with either a disabled active site (C287A mutant) or a disrupted dimer interface (F404D mutant). While AFM imaging revealed correct assembly of 6-nm-spaced two-enzyme DNA nanostructures, both mutants did not exhibit enzymatic activity, reaffirming that DNA origami-mediated caspase-9 activation proceeds via a homodimerization mechanism (Fig. 3e and Supplementary Fig. 14). Second, we investigated the response of DNA origami-mediated caspase-9 activity to the X-linked inhibitor of apoptosis protein (XIAP), an important human regulatory protein that strongly binds to the C-terminal small subunit of caspase-9 (*K*_i_ < 20 nM), forming a heterodimeric complex preventing caspase-9 dimerization^56,57^. After assembly of 6-nm two-enzyme wildtype caspase-9 DNA nanostructures, increasing concentrations of inhibitor were added and protease activity was measured (Fig. 3f). Although the binding affinity of the inhibitor to caspase-9 is in the low nanomolar range, the experiments reveal that very high concentrations (60 equivalents and higher) are needed to effectively inhibit enzyme activity, illustrating the high effective concentration of caspase-9 on the DNA origami platform. Combined, these results demonstrate that our DNA origami platform can serve as a versatile tool for biochemical analysis of intracellular signaling components and their regulation by other proteins, such as inhibitors.

### Co-localization of more than two caspase-9 monomers leads to enhanced enzymatic activity

We have demonstrated that caspase-9 dimerization is sufficient to induce activity, however, the apoptosome is hypothesized to bind up to four enzymes simultaneously^33–35^. We therefore wondered how clustering of enzymes influences their catalytic activity. While multivalent effects on enzyme catalysis have not been described in biochemical literature, previous work by Prins *et al.* on zinc-based catalysts has revealed that clustering of dimerizing subunits can lead to activity enhancement^58^. Using a theoretical model, the authors showed that the activity increase is correlated to a statistical increase in the number of active catalytic units that are formed upon clustering of subunits in a multivalent catalytic system. To investigate the effect of caspase-9 oligomerization, we constructed several multivalent caspase-9 DNA nanostructures, including linear ([**125**]) and triangular ([**123**]) three-enzyme configurations, and a four-enzyme variant ([**1234**]), and confirmed successful assembly using agarose gel electrophoresis (Supplementary Fig. 19) and AFM imaging (Fig. 4a and Supplementary Fig. 20). For both three-enzyme systems, enzymatic activity increased approximately two-fold compared to control configurations [**126**] and [**256**], in which one monomer is positioned such that it cannot interact with the other two monomers (Fig. 4b; top). As expected, the enzyme activity of these ‘2+1’ control configurations was similar to the 6-nm two-enzyme controls. Activity of the four-enzyme [**1234**] proximal configuration increased by 59% compared to the [**1278**] distal control (Fig. 4b; bottom). Since the latter can be viewed as a non-interacting pair of two-enzyme systems on the same DNA origami, it exhibited similar activity compared to the two-enzyme controls, as expected (Fig. 4b; bottom). Collectively, these experimental results demonstrate a clear increase in proteolytic activity when more than two caspase-9 monomers are brought into close proximity.

**Figure 4.**
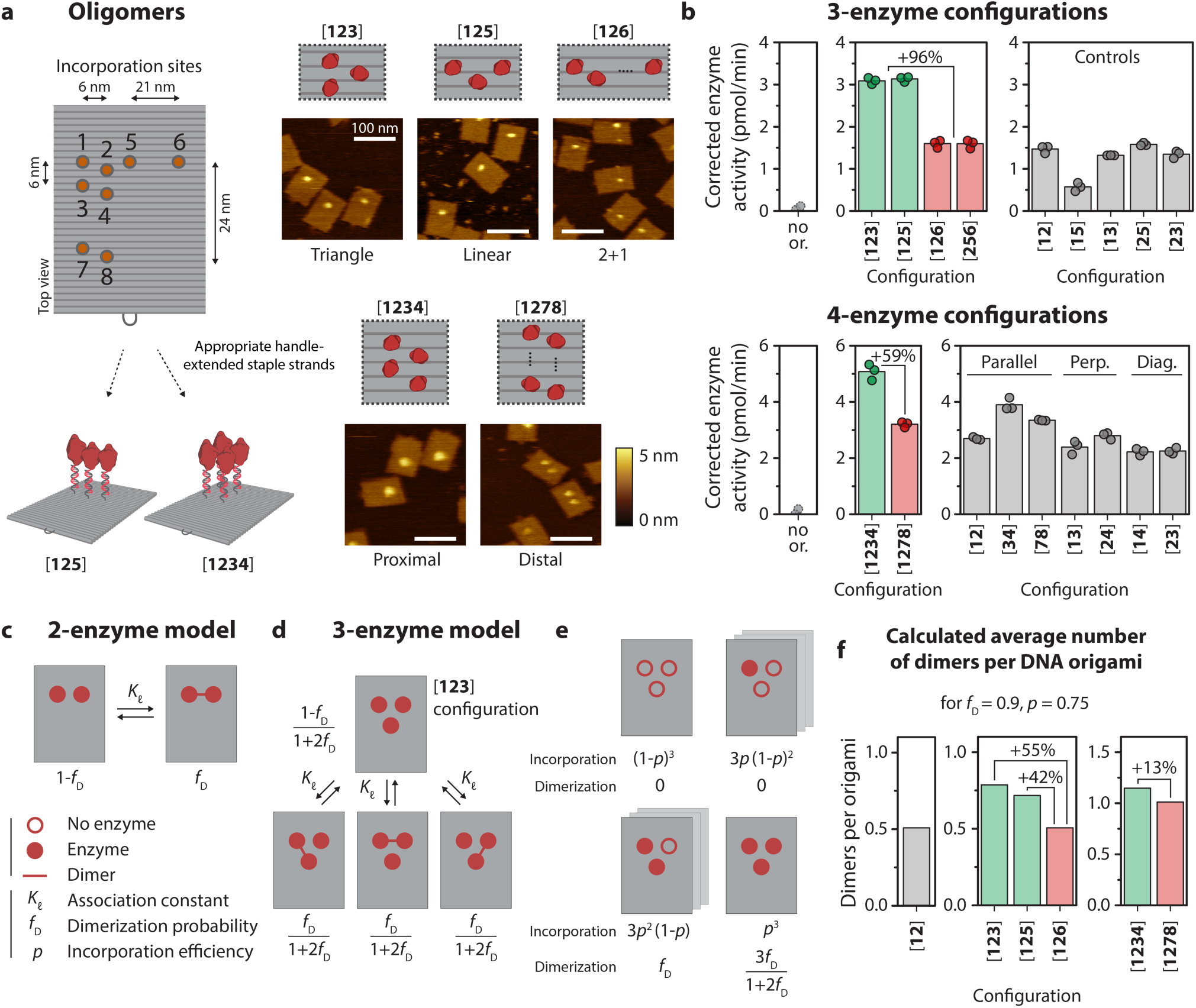
Co-localization of more than two caspase-9 monomers leads to enhanced enzymatic activity. Reactions were performed as described in Fig. 3. The DNA origami concentration was adjusted to keep the total concentration of enzyme at 24 nM (see Methods). All experiments were performed in triplicate. **a,** Schematic overview of possible incorporation sites for handle-extended staple strands (orange circles) for constructing three- and four-enzyme DNA nanostructures. Topographic AFM images show successful incorporation of caspase-9 according to the pre-programmed positions. The bracket notation indicates enzyme configuration as determined by the number and location of the indicated incorporation sites. Color bars indicate height scale. Scale bars, 100 nm. **b** Enzymatic activity measurements for three-enzyme (top) and four-enzyme (bottom) DNA nanostructures (green and red) and two-enzyme controls (gray). Activity was corrected by subtracting the mean background activity (no or.) in all samples. Labels: no or., no DNA origami present; perp., perpendicular arrangement; diag., diagonal arrangement. **c**, Schematic depiction of the mass-balance model for a tethered two-enzyme system, with an equilibrium between a monomeric (left) and dimeric (right) state defined by effective association constant *K*_*ℓ*_, leading to the dimerization probability *f*_D_. **d**, Model for the tethered three-enzyme system in triangular configuration ([**123**]) (see Supplementary Information for models of other configurations). One monomeric and three symmetric dimeric states can be defined, with corresponding state fractions expressed as a function of *f*_D_. **e**, Schematic depiction of the possible configurations in the triangular three-enzyme system with incomplete enzyme incorporation. **f**, When taking into account all possible states and incomplete incorporation, the models predict the average number of enzyme dimers per DNA origami for all two-, three-, and four-enzyme systems.

To dissect the potential factors contributing to the observed activity increase, we developed a thermodynamic model describing dimerization of tethered enzymes in two-, three-, and four-enzyme configurations. The underlying principle of the model is that two tethered enzymes exist in an equilibrium between two concentration-independent states, either as inactive monomers or as a fully active dimer, while other higher-order interactions are not possible (Fig. 4c and Supplementary Information). This allows us to calculate, for each enzyme configuration, the average number of dimers per DNA origami, which we assume is proportional to the experimentally measured caspase-9 activity. The average number of dimers for the 6-nm two-enzyme DNA nanostructure is simply given by the dimerization probability *f*_D_ (with a value of approximately 0.9 at 6 nm monomer separation, see Fig. 3c,d), allowing us to express the number of dimers in higher-order enzyme configurations as a function of system parameter *f*_D_. Applying this approach to, for example, the triangular three-enzyme system (reflecting the experimental [**123**] configuration) we can define four distinct states, one fully monomeric and three symmetric dimeric states, for each of which we derive an expression for the dimer fraction as a function of *f*_D_ (Fig. 4d). In practice, however, enzyme incorporation onto DNA nanostructures is not 100%, resulting in a distribution of species with varying enzyme occupancy, i.e. eight different species with 0, 1, 2, or 3 enzymes (Fig. 4e). We systematically derived expressions for the contribution of each species as a function of *f*_D_ and incorporation efficiency *p* (as determined in Fig. 2), with the sum of all contributions representing the average number of dimers per DNA nanostructure (see Supplementary Information for derivation and for the models of other configurations). Although the model only considers dimerization and does not include any higher-order allosteric effects, the three- and four-enzyme configurations exhibit a moderate increase in the average number of dimers per DNA origami (Fig. 4f; compare green bars to red control bars). This effect has a statistical origin and is correlated to an increase in the number of dimerization possibilities compared to a two-enzyme situation. Interestingly, the models predict that the number of dimers in the [**123**] three-enzyme system is only 55% higher compared to the two-enzyme system, while experimentally an approximately two-fold increase in activity was observed (Fig. 4f; the increase of the linear three-enzyme system [**125**] is 42%). Similarly, the computed number of dimers of the proximal four-enzyme system is only 13% higher than the distal configuration, while experiments indicated a 59% activity increase (Fig. 4f; right graph). In addition to statistical effects related to an increased number of dimerization possibilities, we speculate that the discrepancy between theoretical and experimental results points to an allosteric effect in oligomers of three or more caspase-9 enzymes, leading to an additional enhancement in activity.

Taken together, this combined experimental and theoretical approach allowed us to determine the factors contributing to caspase-9 activation in multi-enzyme assemblies, demonstrating an increase in enzyme activity in assemblies consisting of more than two monomers. Importantly, we found that the presence of multiple co-localized binding sites leads to a statistical enhancement in activity, by increasing the probability of interaction between tethered enzymes when enzyme occupancy is incomplete. We envision that this principle, facilitated by spatial organization of enzymes on SMOCs, could represent a general regulatory mechanism for inducing proximity-driven protein-protein interactions^4,5^.

### Enzymatic activity of the caspase-9 dimer originates from a single catalytic site

Finally, we used the modularity of the DNA origami method to investigate DNA-mediated assembly and activity of caspase-9 heterodimers. Specifically, because the crystal structure of the caspase-9 homodimer reveals that only one of the two active sites is in an accessible, open conformation (Fig. 5a)^36^, we hypothesized that a heterodimer consisting of a wildtype monomer and a mutant with a disabled active site would still display enzymatic activity. Recent work suggests that autoproteolytic processing of the intersubunit linker in wildtype caspase-9 is essential for the correct formation of an active dimeric state^59^. Catalytically inactive point mutant C287A does not undergo autoproteolytic processing, and therefore we induced processing using caspase-3, which is able to cleave caspase-9 in the intersubunit linker to generate a large and small enzyme subunit (Fig. 5b and Supplementary Fig. 5)^60^. We then used wildtype caspase-9 monomer **C** and inactive monomer **I** to assemble homo- and heterodimeric DNA nanostructures, and confirmed that the heterodimeric variants assembled correctly using agarose gel electrophoresis (Supplementary Fig. 21) and AFM imaging (Fig. 5c,d and Supplementary Fig. 22). Remarkably, the level of protease activity of both heterodimeric systems [**C I**] and [**I C**] is equivalent to the activity of the homodimeric wildtype system [**C C**] (Fig. 5e). Control experiments, in which similar heterodimeric configurations were tested with unprocessed **I** and point mutant F404D, exhibited background activity levels (Supplementary Fig. 23). These results suggest that the formation of an active caspase-9 dimer proceeds through an asymmetric mechanism in which only a single active site is brought into an active conformation and that cleavage of the intersubunit linker of both monomers is strictly necessary for enzymatic activity. These experiments illustrate that the modularity of DNA origami-based platforms can be harnessed to investigate relevant biological questions concerning the molecular mechanisms behind interacting signaling proteins.

**Figure 5.**
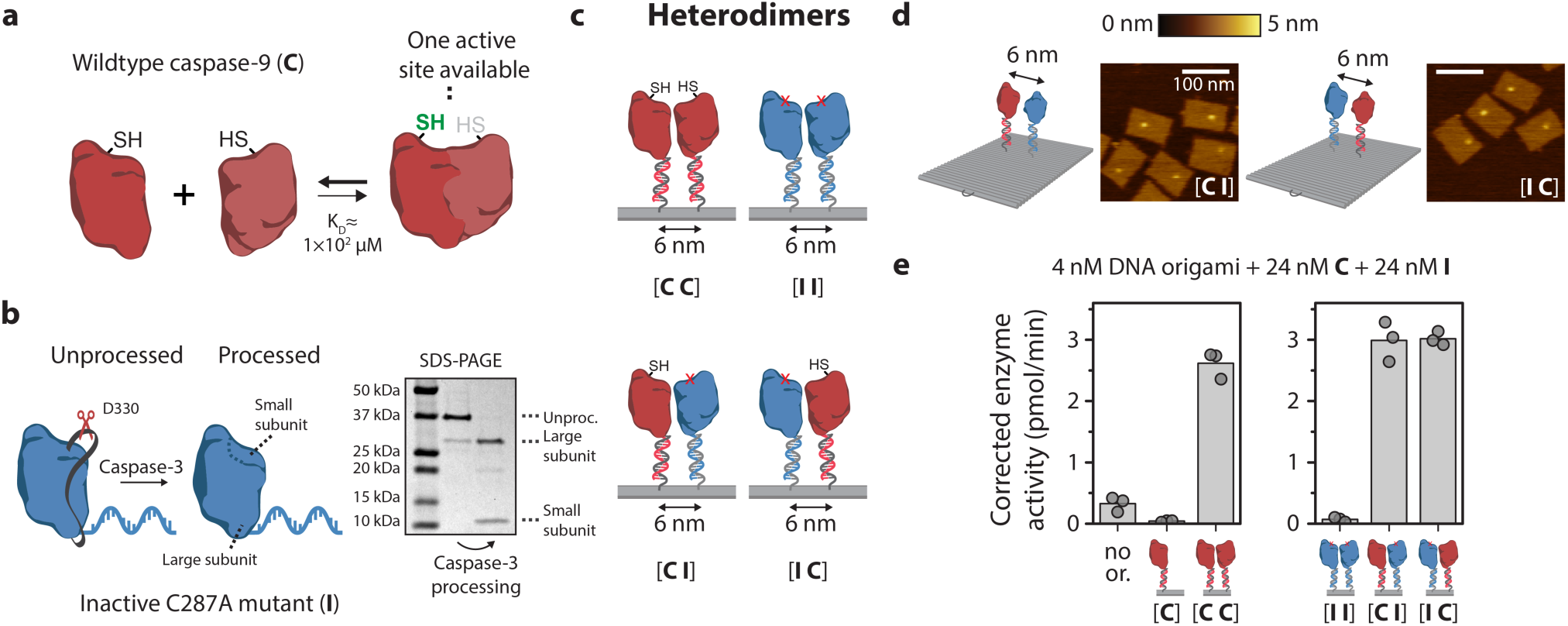
Enzymatic activity of the caspase-9 dimer originates from a single catalytic site. **a,** Schematic overview of wildtype caspase-9 (**C**) homodimerization. Crystal structures of dimeric caspase-9 indicate that only one of the two active sites is in an active, accessible conformation. **b**, Wildtype caspase-9 undergoes autoproteolytic processing at three cleavage sites in the intersubunit linker region (black ribbon), but mutant C287A (**I**) is inactive and therefore does not undergo autoprocessing. To mimic processing, enzyme-DNA conjugates were incubated with caspase-3, a constitutively active protease that also cleaves caspase-9 in the linker region. Purification was performed as described in Fig. 1, and confirmed by SDS-PAGE under reducing conditions. Label: unproc., unprocessed enzyme-DNA conjugate. **c**, The modularity of DNA origami was exploited to construct both homo- and heterodimeric 6-nm two-enzyme DNA nanostructures consisting of combinations of wildtype and inactive monomers. The specific configuration of wildtype (**C**) and inactive (**I**) mutants is denoted using bracket notation. **d**, Topographic AFM images of [**I C**] and [**C I**] heterodimers. Color bar indicates height scale. Scale bars, 100 nm. **e**, Enzymatic activity measurements were performed as described in Fig. 3. In all samples, both 24 nM **C** and 24 nM processed **I** were added (for data with unprocessed **I**, see Supplementary Fig. 23). Activity was corrected by subtracting the mean background activity in all samples. All experiments were performed in triplicate. Label: no or., no DNA origami present.

## Conclusion

Many intracellular signalling proteins assemble into multi-molecular complexes composed of unique combinations of pathway components. Co-localization of proteases, kinases, and phosphatases via their association to dedicated scaffold proteins results in the assembly of higher-order signaling machines such as the myddosome, the apoptosome, and the necrosome, that are able to efficiently control signal transmission via proximity-driven enzyme activation^4,5,61^. Here, we show that DNA origami can be used as a unique *in vitro* platform for constructing synthetic higher-order signaling machines. As a proof-of-principle, we engineered synthetic DNA origami-based variants of the apoptosome and revealed how the distance and number of caspase-9 monomers influence enzymatic activity. Our results reveal a multivalent catalytic effect as evidenced by an increase in catalytic activity in three- and four-enzyme systems compared to a two-enzyme configuration. A thermodynamic model based on tethered dimerization revealed that the observed activity enhancement partially originates from a statistical increase in the number of active catalytic units in higher-order enzyme configurations. We envision that clustering of catalytic enzymatic subunits into higher-order complexes, either through SMOC-based assembly^4,5^, functional homotypic interactions^62,63^, or via liquid phase separation^64,65^, could represent a general mechanism for enzymatic activity enhancement or regulation in various intracellular processes.

In contrast to other available platforms for engineering higher-order signaling machines, such as synthetic protein scaffolds^7^ or leucine zipper-induced assemblies^45^, DNA origami allows oligomerization of non-identical signaling proteins and user-defined control over their number, position, and relative geometry. The construction of higher-order signaling complexes using DNA-origami-based SMOCs allows a detailed analysis of their function and can be used to probe unresolved molecular mechanisms in intracellular signaling, such as for example the multivalent enhancement of catalytic activity as reported in this work. Recent work has shown the possibility of genetically encoded DNA and RNA nanostructures and revealed successful intracellular assembly of a simple DNA cross-over nanostructure on which proteins could be organized using orthogonal zinc fingers^66–68^. As such, *in vivo* production and assembly of DNA-based SMOCs, analogous to those developed by us, is a realistic possibility and could find application as modular synthetic control elements for diversifying signaling dynamics of existing pathways. We anticipate that DNA origami platforms will find broad use to inform the function of many other important SMOCs for which oligomerization-driven allosteric regulation of non-identical enzymes, such as for example kinases, is a common regulatory principle.

## Acknowledgements

We thank Joost van Dongen for help with the mass spectrometry analyses, Nick van der Zon for initial protein expression experiments, and Glenn Cremers for helpful discussions. The ICMS Animation Studio contributed the cartoons of DNA strands and the DNA origami structure. This work was supported by the European Research Council, ERC (project n. 677313 BioCircuit), an NWO-VIDI grant from the Netherlands Organization for Scientific Research (NWO, 723.016.003) and funding from the Ministry of Education, Culture and Science (Gravity programs, 024.001.035 & 024.003.013).

## Author contributions

B.R. designed the study, performed experiments, developed the geometric model, analyzed the data, and wrote the manuscript. A.M. developed and derived the thermodynamic model and analyzed the data. B.G.A. performed and analyzed all AFM measurements. J.R. performed molecular dynamics simulations. A.d.H. performed initial protein expression and provided critical input for the experiments. L.B. supervised the study and provided critical feedback on the manuscript. T.d.G. conceived, designed, and supervised the study, analyzed the data, and wrote the manuscript. All authors discussed the results and commented on the manuscript.

## Data availability

The data that support the findings of this study are available from the corresponding authors, upon reasonable request.

