## Supporting Information for "A DNA-based synthetic apoptosome"

### Contents

Methods ... 3
Chemicals and reagents ... 3
Expression and purification of caspase-9 ... 3
DNA and protein sequence caspase-9 ... 4
Functionalization of oligonucleotides ... 4
Enzyme-DNA conjugation ... 5
Assembly of caspase-9 on template T ... 5
Self-assembly of DNA origami nanostructures ... 5
Assembly of caspase-9 on DNA origami ... 6
Activity assays ... 6
Activity assay data processing and correction ... 7
Gel electrophoresis ... 7
AFM imaging and analysis ... 8
Geometric model ... 8
Supplementary results ... 10
Expression and purification of caspase-9 variants ... 10
Enzyme-DNA conjugation and purification ... 12
Calibration of substrate LEHD-AFC ... 17
Agarose gel analysis of two-enzyme DNA nanostructures ... 18
AFM analysis of DNA origami and one-enzyme DNA nanostructures ... 19
AFM analysis of two-enzyme DNA nanostructures ... 20
Control experiments for two-enzyme DNA nanostructures ... 24
Geometric model and molecular dynamics simulations ... 26
Agarose gel and AFM analysis of three- and four-enzyme DNA nanostructures ... 27
Agarose gel and AFM analysis of heterodimer DNA nanostructures ... 28
Activity of unprocessed heterodimer DNA nanostructures ... 30
Derivation of the thermodynamic models ... 31
Section 1: dimerization of tethered enzymes ... 31
Section 2: dimerization of tethered enzymes with incomplete enzyme incorporation ... 38
DNA origami design and DNA sequences ... 47
Supplementary references ... 54

### Methods

#### Chemicals and reagents

All reagents and solvents were obtained from commercial sources and used without further purification. Unnatural amino acid *p*-azidophenylalanine was purchased from Bachem. Mutagenesis primers and M13mp18 scaffold were purchased from Eurofins. 7-amino-4-(trifluoromethyl)coumarin (AFC) was obtained from Fluorochem, while synthetic tetrapeptide caspase-9 substrate Ac-LEHD-AFC was purchased from Enzo Life Sciences. Amino-functionalized oligonucleotides (ODNs) were obtained HPLC-purified from Integrated DNA Technologies and dissolved in DNase/RNase-free water at 250 µM. Unmodified ODNs were obtained in desalted form from Integrated DNA Technologies and dissolved at a stock concentration of 500 µM in DNase/RNase-free water.

#### Expression and purification of caspase-9

The catalytic domain of human caspase-9 (140-416) with an N-terminal amber stop codon was encoded on a pET28a plasmid and synthesized by GenScript. The construct contains an N-terminal His-SUMO tag (the SUMO tag was included to improve stability and solubility during expression) and a C-terminal Strep-tag (see Supplementary Section 'DNA and protein sequence caspase-9'). C287A and F404D mutants were generated using the QuikChange Lightning Multi Site-Directed Mutagenesis kit (Agilent), according to the manufacturer's instructions and using the primers in **Supplementary Table 1**. The pEVOL-pAzF vector, encoding for the orthogonal aminoacyl tRNA synthetase/tRNA pair, was a kind gift from Peter Schultz (Addgene plasmid #31186). Both plasmids were transformed into E. coli BL21(DE3) competent bacteria (Novagen) and cultured at 37°C in 500 mL 1xTB (terrific broth, VWR) supplemented with 0.4% (v/v) glycerol, 25 µg/mL kanamycin and 25 µg/mL chloramphenicol. Protein expression was induced at $OD_{600}$=0.6 by addition of 1 mM β-D-1-thiogalactopyranoside and 0.02% (w/v) arabinose. Simultaneously, the non-canonical amino acid *p*-azidophenylalanine was added directly to the culture medium at a final concentration of 1 mM. Expression was carried out for ~16 h at 18°C. Cells were harvested by centrifugation at 10,000 g for 10 min at 4°C, the pellet resuspended in lysis buffer (10 mL per gram pellet, 1x PBS, 370 mM NaCl, 10% (v/v) glycerol, 20 mM imidazole, Benzonase nuclease (25 U per 10 mL buffer, Novagen), pH 7.4) and the cells were lysed using an EmulsiFlexC3 High Pressure homogenizer (Avestin) at 15,000 psi for two rounds. The soluble fraction (cleared lysate) containing caspase-9 was collected by centrifugation at 40,000 g for 30 min at 4°C.

Typically, non-canonical amino acids are not fully incorporated leading to the presence of truncated protein fragments. Therefore, purification was performed by both $Ni^{2+}$-affinity chromatography and *Strep*-tactin affinity chromatography, using the N- and C-terminal affinity tags on caspase-9, respectively. The cleared lysate was loaded on a Ni-charged column (His-Bind® Resin, Novagen), washed with wash buffer (1x PBS, 370 mM NaCl, 10% (v/v) glycerol, 20 mM imidazole, pH 7.4), and eluted with elution buffer (1x PBS, 370 mM NaCl, 10% (v/v) glycerol, 250 mM imidazole, pH 7.4). Cleavage of the N-terminal His-SUMO tag was performed by adding SUMO protease dtUD1 (1:500, purified according to standard procedure[1]) and 2 mM EDTA to the elution fractions, while dialyzing (MWCO 3.5 kDa, Thermo Fisher) against 4 L of dialysis buffer (50 mM Tris, 150 mM NaCl, pH 8.0) for ~16 hr at 4°C. Finally, the concentrate was applied to a *Strep*-tactin column (Superflow® resin, IBA Life Sciences). The column was washed with wash buffer (100 mM Tris-HCl, 150 mM NaCl, 1 mM EDTA, pH 8.0) and the protein eluted with wash buffer supplemented with 2.5 mM desthiobiotin. Elution fractions were combined and concentrated using Amicon 10 kDa MWCO centrifugal filters (Merck Millipore) to a final concentration of ~1.5 mg/mL (~47 µM), and then snap frozen in liquid nitrogen and stored in 100 µL aliquots at -80°C. The concentration of caspase-9 was determined by measuring the absorption at 280 nm (NanoDrop 1000, Thermo Scientific) assuming an extinction coefficient of $3.1 \times 10^4$ $M^{-1}$ $cm^{-1}$ [2]. Total yield after purification typically was ~4 mg/L culture medium. Purity of caspase-9 was assessed using SDS-

PAGE under reducing conditions (**Supplementary Figure 1**). Molecular weight was confirmed using liquid chromatography quadrupole time-of-flight mass spectrometry (Waters ACQUITY UPLC I-Class System coupled to a Xevo G2 Q-ToF) by injecting a 0.1 µL sample into an Agilent Polaris C18A RP column with a flow of 0.3 mL/min and a 15-60% acetonitrile gradient containing 0.1% formic acid (**Supplementary Figure 2**).

#### DNA and protein sequence caspase-9

The single-letter amino acid code is shown in uppercase, with above in lowercase the corresponding DNA sequence. The N-terminal hexahistidine tag is shown in green, the SUMO domain in orange, the catalytic domain of caspase-9 (140-416) in red, and the C-terminal *Strep*-tag in blue. The position of the non-canonical amino acid *p*-azidophenylalanine is indicated with an X, located at the N-terminus of the enzyme after removal of the SUMO domain. Relevant amino acids in caspase-9 are indicated, following the numbering system of PDB entry 1NW9: C287 is the cysteine in the active site, F404 is the phenylalanine present in the dimer interface, and E306, D315, and D330 are the proteolytic cleavage sites in the linker region between the N-terminal large subunit and the C-terminal small subunit. Autocleavage by caspase-9 occurs pre-dominantly at E306 and D315 and to a lesser extent at D330, while caspase-3 cleaves exclusively at D330[3].

```
 atgggcagcagccatcatcatcatcatcacagcagcggcctggtgccgcgcggcagccatatgggtggtagcgacagcgaagtgaatcaa
  M  G  S  S  H  H  H  H  H  H  S  S  G  L  V  P  R  G  S  H  M  G  G  S  D  S  E  V  N  Q

 gaagccaaaccggaagtgaaaccggaagtcaaaccggaaacgcacatcaacctgaaagtgagtgatggtagctctgaaattttctttaaa
  E  A  K  P  E  V  K  P  E  V  K  P  E  T  H  I  N  L  K  V  S  D  G  S  S  E  I  F  F  K

 attaagaaaaccacgccgctgcgtcgcctgatggaagcgtttgccaaacgccagggtaaagaaatggatagcctgcgcttcctgtatgac
  I  K  K  T  T  P  L  R  R  L  M  E  A  F  A  K  R  Q  G  K  E  M  D  S  L  R  F  L  Y  D

 ggcattcgtatccaggcggatcaagccccggaagatctggacatggaagataacgacattatcgaagcgcatcgcgaacagattggcggt
  G  I  R  I  Q  A  D  Q  A  P  E  D  L  D  M  E  D  N  D  I  I  E  A  H  R  E  Q  I  G  G
        140
 tagggcggtggcgccctggaatcactgcgtggtaatgcagatctggcttacatcctgtcgatggaaccgtgcggccactgtctgatcatc
  X  G  G  G  A  L  E  S  L  R  G  N  A  D  L  A  Y  I  L  S  M  E  P  C  G  H  C  L  I  I

 aacaacgttaacttctgccgtgaaagtggtctgcgcacccgtacgggctccaatattgactgtgaaaaactgcgtcgccgttttagttcc
  N  N  V  N  F  C  R  E  S  G  L  R  T  R  T  G  S  N  I  D  C  E  K  L  R  R  R  F  S  S

 ctgcatttcatggtggaagttaaaggtgatctgaccgcgaagaaaatggtgctggcgctgctggaactggcccagcaagatcacggcgcc
  L  H  F  M  V  E  V  K  G  D  L  T  A  K  K  M  V  L  A  L  L  E  L  A  Q  Q  D  H  G  A

 ctggactgctgtgtggttgtcatcctgagccatggttgccaggcatctcacctgcaatttccgggcgctgtctatggtaccgatggctgt
  L  D  C  C  V  V  V  I  L  S  H  G  C  Q  A  S  H  L  Q  F  P  G  A  V  Y  G  T  D  G  C

 ccggtctccgtggaaaaaattgtgaacatcttcaatggtacgagctgcccgtctctgggtggcaaaccgaaactgtttttcattcaggca
  P  V  S  V  E  K  I  V  N  I  F  N  G  T  S  C  P  S  L  G  G  K  P  K  L  F  F  I  Q  A
287                                                      306                        315
 tgtggtggcgaacaaaaagatcatggctttgaagttgctagcacctctccggaagacgaaagtccgggttccaacccggaaccggatgcc
  C  G  G  E  Q  K  D  H  G  F  E  V  A  S  T  S  P  E  D  E  S  P  G  S  N  P  E  P  D  A
                                       330
 accccgtttcaggaaggcctgcgcacgttcgatcaactggacgcaatttcatcgctgccgaccccgagcgacatctttgtgagttattcc
  T  P  F  Q  E  G  L  R  T  F  D  Q  L  D  A  I  S  S  L  P  T  P  S  D  I  F  V  S  Y  S

 acgtttccgggtttcgtttcttggcgtgatccgaaatcaggctcgtggtatgtggaaaccctggatgacattttcgaacagtgggcgcac
  T  F  P  G  F  V  S  W  R  D  P  K  S  G  S  W  Y  V  E  T  L  D  D  I  F  E  Q  W  A  H
                                                                                 404
 tcagaagatctgcaatcgctgctgctgcgcgtcgcaaacgctgtttctgtcaaaggtatctacaaacagatgccgggctgttttaatttc
  S  E  D  L  Q  S  L  L  L  R  V  A  N  A  V  S  V  K  G  I  Y  K  Q  M  P  G  C  F  N  F
                           416
 ctgcgtaaaaaactgttctttaaaacgtcaggtggttcgtggagccatccgcaatttgaaaaataa
  L  R  K  K  L  F  F  K  T  S  G  G  S  W  S  H  P  Q  F  E  K  *
```

#### Functionalization of oligonucleotides

Functionalization of amino-modified ODNs with bicyclononyne (BCN) was performed on a 100 µL scale by adding 2 mM (1*R*,8*S*,9*s*)-bicyclo[6.1.0]non-4-yn-9-ylmethyl *N*-succinimidyl carbonate (BCN-NHS, 4 mM stock solution in dry DMSO) to 100 µM of ODN in 1xPBS (pH 7.4), and incubating for 2 h at 25°C under continuous shaking. Excess BCN-NHS was removed by two rounds of ethanol precipitation. Briefly, 900 µL ice-cold 100% ethanol and 20 µL 3 M potassium acetate (pH 5.5) was added directly to the reaction mixture and incubated for 30 min at -30°C. After centrifugation (14,000 g for 30 min at 4°C) the supernatant was removed and the pellet was reconstituted in 100 µL water. This procedure was repeated once and after centrifugation the pellet was washed with 95% ice-cold ethanol (v/v, in water). The mixture was centrifuged again (14,000 g for 10 min at 4°C) and the pellet was reconstituted in TE buffer (10 mM Tris-HCl, 1 mM EDTA, pH 8.0) and stored at -30°C.

Molecular weight was confirmed using mass spectrometry by flow injection analysis on an LCQ fleet (Thermo Finnigan) ion-trap mass spectrometer in negative mode, using direct injection of 5 µL DNA (10 µM in 30% isopropanol (v/v in water), 0.1% triethylamine (v/v), pH 10) (**Supplementary Figure 3**). Concentrations were determined using absorption at 260 nm (NanoDrop 1000, Thermo Scientific), assuming extinction coefficients reported by the manufacturer (**Supplementary Table 2**).

#### Enzyme-DNA conjugation

Before conjugation, 100 µL caspase-9 aliquots were buffer exchanged to wash buffer (100 mM Tris-HCl, 150 mM NaCl, 1 mM EDTA, pH 8.0) using Zeba desalting columns (7 kDa MWCO, 0.5 mL, Thermo Scientific) according to the manufacturer's instructions, to remove all remaining desthiobiotin. Typically, conjugation reactions were carried out on a 500 µL scale using 10 µM protein and 30 µM BCN-DNA in reaction buffer (100 mM Tris-HCl, 150 mM NaCl, 1 mM EDTA, 0.1% CHAPS (w/v), pH 8.0) for ~16 hr at 4°C. Competing thiol-yne reactions of BCN-DNA with the 11 cysteines in caspase-9 were suppressed by pre-incubating the protein with 1 mM β-mercaptoethanol for 30 min at 4°C[4]. For proteolytic processing of C287A and F404D mutants, 0.5 µM active caspase-3 (expressed and purified as described[5]) was added directly after the conjugation reaction and incubated for 2 hr at 18°C.

To remove excess BCN-DNA, *Strep*-tactin affinity chromatography was performed as described above. Then, ion-exchange chromatography was performed to remove unreacted protein. After equilibration of the ion-exchange column (0.5 mL strong anion-exchange spin columns, Thermo Scientific) with purification buffer (100 mM Tris-HCl, 150 mM NaCl, 1 mM EDTA, 1 mM DTT, 0.1% CHAPS (w/v), pH 8.0), the protein mixture was directly loaded onto the column in 400 µL fractions, according to the manufacturer's instructions. Elution was performed by stepwise increase of the NaCl concentration in the buffer (200, 300, 400, 500, 600 mM, respectively). Typically, the protein eluted at <300 mM NaCl, while enzyme-DNA conjugates eluted at 500-600 mM NaCl (**Supplementary Figure 4** and **Supplementary Figure 5**). Elution fractions containing pure enzyme-DNA conjugates were pooled, supplemented with glycerol (5% (v/v) final concentration), snap frozen in liquid nitrogen, and stored at -80°C in 5 µL aliquots.

#### Assembly of caspase-9 on template T

The linear 30-nt single-stranded template **T** was designed to act as a bivalent scaffold for caspase-9 (5' to 3': TCATACGACTCACTCCTGACTGACTGACTG), simultaneously hybridizing to enzyme-DNA conjugates with anti-handle sequences a1 and a2 (**Supplementary Table 2**), similar to designs used in literature[6,7]. Activation of caspase-9 with template **T** was performed by adding the two caspase-9 enzyme-DNA conjugates and template **T** in equimolar amounts (4 nM) for 2 hr at 4°C in activity buffer (10 mM Tris, 1 mM EDTA, 10 mM $MgCl_2$, 100 mM NaCl, 1 mM DTT, 0.1% (w/v) CHAPS, pH 7.5). Enzyme activity was measured as described below.

#### Self-assembly of DNA origami nanostructures

The DNA origami rectangle used in this study was based on the *tall rectangle* design by Rothemund[8]. The 7249-nt single-stranded M13mp18 scaffold strand folds into a single-layer structure of 32 helices using 192 staple strands. DNA nanostructure variants for enzyme incorporation were constructed by replacing structural staple strands with staple strands extended with handles complementary to the anti-handles of the caspase-9 enzyme-DNA conjugates. For detailed notes on the design and the construction of all DNA nanostructure variants used in this study, see Supplementary Section 'DNA origami design and DNA sequences'.

Folding reactions were performed at a volume of 50 µL in folding buffer (10 mM Tris, 1 mM EDTA, 10 mM $MgCl_2$, 50 mM NaCl, pH 8.0), with 25 nM scaffold strand and 250 nM of each staple strand. The reaction mixture was heated to 95°C for 15 min and then slowly cooled to 20°C at a rate of 1°C/min. Excess staple strands were removed using 100 kDa MWCO 0.5 mL Amicon centrifugal filters (Merck Millipore). Briefly, a filter was pre-wetted with 500 µL purification buffer (10 mM Tris, 1 mM EDTA, 10 mM $MgCl_2$, 100 mM NaCl, pH 8.0). The folding mixture was diluted to 500 µL with purification buffer, added to the filter and centrifuged at 4°C for 5 min at 5,000 g. This step was repeated for a total of three washing steps. The concentrate was recovered by inverting the filter and spinning for 2 min at 1,000 g. The concentration of purified enzyme-DNA conjugates was determined with gel densitometry on SDS-PAGE. To this end, conjugate gel band intensity was determined using the ImageJ (v1.52n) gel analysis plugin and then compared to a calibration curve of known concentrations of protein (**Supplementary Figure 7**). Samples were stored in DNA LoBind tubes (Eppendorf) at 4°C and used on the same day. The DNA origami concentration was determined by measuring the absorption at 260 nm, assuming an extinction coefficient of $1.24\times10^{8}$ $M^{-1}$ $cm^{-1}$ [9,10].

#### Assembly of caspase-9 on DNA origami

Incorporation of enzyme-DNA conjugates onto purified DNA origami nanostructures was performed by incubating DNA origami with 3 molar equivalents of caspase-9 DNA conjugate per handle for 2 hr at 4°C in activity buffer (10 mM Tris, 1 mM EDTA, 10 mM $MgCl_2$, 100 mM NaCl, 1 mM DTT, 0.1% (w/v) CHAPS, pH 7.6). The total concentration of caspase-9 DNA conjugate was kept constant at 24 nM for all experiments, reflecting typical concentrations in the cytosol[11]. As a result, 4 nM, 2.67 nM, and 2 nM DNA origami was used for two-, three-, and four-enzyme DNA nanostructures, respectively. To allow comparison of the activity of three-enzyme DNA nanostructures with two-enzyme controls, all experiments in the top row of **Figure 4b** including the two-enzyme configurations, were performed with 2.67 nM DNA origami. For heterodimer experiments, the concentration of each enzyme-DNA conjugate was kept at 24 nM, which results in the same background activity originating from wildtype caspase-9. For inhibition experiments, varying concentrations of XIAP (human recombinant BIR3-XIAP, R&D Systems) were added after caspase-9 incorporation, and the reaction was incubated additionally for 1 hr at 4°C before measuring enzyme activity at 18°C, as described below.

#### Activity assays

Enzyme activity was measured using the synthetic tetrapeptide caspase-9 substrate LEHD (dissolved in dry DMSO at 10 mM)[12], which is cleaved by caspase-9 after the aspartic acid residue releasing and unquenching the fluorescent dye 7-amino-4-(trifluoromethyl)coumarin (AFC). After assembly of caspase-9 on DNA origami, substrate was added to a final concentration of 167 µM and proteolytic cleavage was monitored over time in 384-well plates (60 µL reaction volume) at 18°C by measuring fluorescence (ex.: 400 nm, em.: 505 nm) in a Tecan Spark 10M platereader. Fluorescence units were converted to concentration using a calibration curve (**Supplementary Figure 8**). To this end, varying concentrations of free AFC (dissolved in dry DMSO at 10 mM) were added to activity buffer, and fluorescence was measured as described. Raw data of all activity assays were extracted, converted, and formatted using in-house MATLAB scripts (R2015a, available upon request). Enzyme activity (in pmol $min^{-1}$) was determined by fitting the initial slope (20-60 min) of the kinetic trace to a linear curve. Kinetic parameters were determined by measuring enzyme activity as a function of varying LEHD concentration between 0-1.5 mM, and fitting the results with the standard Michaelis-Menten equation:

$$\text{activity} = \frac{V_{\max}\,[\text{LEHD}]}{K_{\text{M}}+[\text{LEHD}]} \quad (1)$$

with $V_{\max}$ the maximum rate (in pmol $min^{-1}$) and $K_{\text{M}}$ the Michaelis constant (in mM). Because high concentrations of the substrate (>0.5 mM) strongly influence the pH, kinetic analyses were performed at a higher buffer concentration (50 mM HEPES, 5 mM Tris, 1 mM EDTA, 10 mM $MgCl_2$, 100 mM NaCl, 1 mM DTT, 0.1% (w/v) CHAPS, pH 7.6).

#### Activity assay data processing and correction

Typically, an excess of caspase-9 enzyme-DNA conjugate was used to incorporate caspase-9 onto DNA nanostructures, leading to background activity originating from untethered enzymes remaining in solution. In some cases background correction was performed by measuring the activity of untethered caspase-9 without DNA origami, at the same concentration as in measurements with DNA origami, in triplicate and in parallel for each experiment (indicated with 'no origami'). Subsequently, the mean activity was subtracted from the enzyme activity of other samples, leading to the *corrected enzyme activity,* as reported in **Figures 3b**, **4b**, and **5e**. The corrected enzyme activity for one-enzyme DNA nanostructures (see **Figure 5e**) and two-enzyme DNA nanostructures at large monomer separation (>20 nm, see **Figure 3b**) is similar to both 'no origami' and 'no handles' samples (see **Figure 3b**), suggesting that untethered caspase-9 in solution does not interact with or influence the behavior of caspase-9 on DNA nanostructures.

To compare activation of caspase-9 on DNA nanostructures with bivalent template **T** as reported in **Figure 3c**, activity normalization was performed. Due to incomplete enzyme incorporation (75% per handle, as determined in **Figure 2**), only ~56% (or ~2.3 nM) of the two-enzyme DNA nanostructures contain two enzymes and are therefore in an active state. Because the enzyme activity of caspase-9 on DNA nanostructures is concentration independent, the corrected enzyme activity was normalized to 100% enzyme incorporation (or 4 nM DNA origami), leading to the *normalized enzyme activity* as reported in **Figure 3c**. Assuming quantitative assembly of caspase-9 on template **T** (melting temperature >40°C for 15-nt handle-anti-handle duplexes at 4 nM[13]), this allows comparison of the activity of caspase-9 assembled on either DNA origami or 4 nM template **T**. The fold change in **Figure 3c** was calculated based on the untethered enzyme activity of 4 nM caspase-9 in solution (indicated with 'non-tet.').

#### Gel electrophoresis

Conjugation reactions and purification of conjugates were monitored by SDS-PAGE under reducing conditions. Samples were heated at 95°C for 5 min in sample buffer (62.5 mM Tris, 10% glycerol, 50 mM DTT, 2.5% SDS (w/v), 0.01% bromophenol blue, pH 6.8) and run on pre-cast 4-20% Mini-PROTEAN® TGX gels (Bio-rad) in running buffer (25 mM Tris, 192 mM glycine, 0.1% SDS, pH 8.3) for 45-50 min at 150 V. Precision Plus Protein All Blue ladder (Bio-Rad) was used as a reference. Gels were stained with Coomassie Brilliant Blue G-250 (Bio-Rad), imaged using an ImageQuant 400 Digital Imager (GE Healthcare), and analyzed with ImageJ.

Urea-PAGE analysis was used to confirm removal of excess DNA after enzyme-DNA conjugate purification (**Supplementary Figure 6**). Gels were prepared at 15% monomer concentration using the manufacturer's protocol (UreaGel, National Diagnostics). Samples were two-fold diluted in sample buffer (10 mM Tris, 1 mM EDTA, 0.01% xylene cyanol (w/v), 8 M urea, pH 8.0). DNA reference samples were prepared by four-fold serial dilution from a 2 µM stock solution of ODN a1 (see **Supplementary Table 2**) in water. The gel was run in 1x TBE (89 mM Tris-HCl, 89 mM boric acid, 2 mM EDTA, pH 8.0) for 60 min at 150 V and post-stained with SYBR Gold (Thermo Scientific). GeneRuler Ultra Low Range DNA ladder (Thermo Scientific) was included as a reference. Gels were imaged using an ImageQuant 400 Digital Imager (GE Healthcare) and analyzed with ImageJ.

Agarose gel electrophoresis was used for DNA origami folding analysis. In short, 1.5% agarose gels were cast in gel buffer (1x TAE, 10 mM $MgCl_2$, pH 8.0) supplemented with SYBR Safe. Gels were run in gel buffer for 90 min at 65 V in an ice bath. DNA origami samples were diluted just before loading to a final concentration of 2 nM and Ficoll-400 (final concentration 1.5% (w/v) was added. Gels were imaged using an ImageQuant 400 Digital Imager (GE Healthcare) and analyzed with ImageJ.

#### AFM imaging and analysis

Enzyme-functionalized DNA nanostructures were prepared as described, and purified using gel extraction. In short, agarose gel electrophoresis was performed as described and upon completion the correct DNA nanostructure band was excised from the gel. The band was cut into small pieces and loaded onto a Freeze 'N' Squeeze column (Bio-Rad). After centrifugation for 4 min at 1,000 g, the supernatant was collected and stored at 4°C.

Topographic AFM images were acquired in AC mode at room temperature under liquid conditions using an MFP-3D AFM (Asylum Research) and V-shaped $Si_3N_4$ cantilevers with sharpened pyramidal tip and a nominal spring constant of 0.04 N/m (OTR4, Bruker AFM Probes). Circular mica substrates (Ted Pella) were glued to Teflon (VWR) using epoxy-based mounting glue before use. Gel-purified DNA nanostructure solutions were first diluted to 2 nM in imaging buffer (10 mM Tris, 1 mM EDTA, 10 mM $MgCl_2$, pH 8.0) and then 10 µL was incubated for 30 s on a freshly-cleaved mica surface. Subsequently, 50 µL of imaging buffer was used to rinse the sample twice and finally 100 µL of imaging buffer was added to perform the AFM imaging. Topographic images of either 1.0×1.0 µm² or 1.5×1.5 µm² (512×512 px) were acquired in various regions of the mica substrate, using drive amplitudes within the range of 0.6-1.0 V, optimizing the scanning and feedback parameters for each image.

Data processing was performed using a combination of custom-written MATLAB code and Gwyddion (v2.51) software. Trace and retrace images were combined into a single image by determining the offset between the two using two-dimensional cross-correlation and averaging of the pixel values. Enzyme incorporation efficiency per handle was calculated by counting all well-formed, intact DNA origami rectangles in at least 4 separate AFM images, and determining the presence of 0, 1, or 2 caspase-9 monomers (**Supplementary Figure 13**). At least 250 DNA nanostructures were analyzed for each sample.

#### Geometric model

Tethered handle-anti-handle movement was determined using coarse-grained molecular dynamics simulations performed with oxDNA (v2.2.2)[14–16]. Coarse-graining is done by representing each nucleotide by a rigid body interacting with an effective potential that contains terms for backbone connectivity, excluded volume, hydrogen bonding, stacking and electrostatic interactions. The initial configuration of the system with a 20 bp handle was generated by converting the caDNAno file using a Python script included in the oxDNA package. The handle, positioned next to the DNA origami rectangle in the caDNAno design, was then translated towards the vicinity of the attaching staple strand and rotated 90 degrees with respect to the DNA rectangle using an in-house Python script. The initial configuration was relaxed to the oxDNA2 model by running a ~150 ps simulation at 1 K with a strong temperature coupling. Subsequently, three ~1.5 µs MD production runs with different random seeds were performed with ~15 fs time steps using an Andersen-like thermostat at 25°C and a NaCl concentration of 500 mM. Simulation parameters were used as described[17], and snapshots were saved every $10^4$ time step. Finally, using an in-house Python script, the position of the end of the handle relative to the DNA origami was analyzed for every snapshot. To this end, we defined the origami plane by the vectors $\boldsymbol{v}$ and $\boldsymbol{w}$ through the handle attachment point, where $\boldsymbol{v}$ is defined as the vector between the centers of mass of the base pairs 32 bases on either end of the handle attachment point parallel to the DNA origami helices, and $\boldsymbol{w}$ is defined as the vector between the centers of mass of the base pairs 4 helices away on either end of the handle attachment point perpendicular to the DNA origami helices (**Supplementary Figure 18**). Then, the center of mass of the base pair at the end of the handle is projected onto the normal vector of the plane to determine the height ($h$), and onto the plane itself to determine the distance to the handle attachment point ($r$). From this, we determine the two-dimensional histogram as a function of $r$ and $h$, and normalize by calculating the binned probability density $\rho_{\mathrm{j,k}}$ (in $\mathrm{nm}^{-2}$) according to

$$\rho_{\mathrm{j,k}}(r,h) = \frac{N_{\mathrm{j,k}}}{A_{\mathrm{j,k}} \sum_{\mathrm{j,k}} N_{\mathrm{j,k}}} \quad (2)$$

with $N_{\mathrm{j,k}}$ the number of counts per bin $(j,k)$ and $A_{\mathrm{j,k}}$ the area of that bin (**Supplementary Figure 18**).

Based on the results of the simulations, a three-dimensional geometric model was constructed using Mathematica (v10, Wolfram) based on a system with two tethered particles separated by a distance $s$. The diameter of a caspase-9 monomer was estimated to be 4.5 nm based on the crystal structure (PDB: 1JXQ). The tether connecting the enzyme to the DNA origami scaffold consists of the 15-bp handle-anti-handle DNA duplex, the BCN-azide moiety, single-stranded DNA linkers, and a short peptide linker, and was estimated to have a total length of approximately 8 nm (**Supplementary Figure 18**). By following the approach of Van Valen *et al.*[18], the model allows us to calculate the probability that the tethered particles form a dimer (expressed as the dimerization probability $f_{\mathrm{D}}$) as a function of monomer separation $s$ and the $K_{\mathrm{D}}$ of dimerization in solution. The approach relies on treating the particles as point objects and assuming that the particles are free to dimerize when they are within each other's vicinity. The probability that the particles are close depends on the exact conformational movement of both particles. A particle's conformational space with volume $v_{\mathrm{s}}$ is defined based on a hemi-shell bounded by an angle of 35° with the scaffold, as determined by the overall movement of the handle-anti-handle duplex in the coarse-grained simulations (**Supplementary Figure 18**). The intersection with volume $v_{\mathrm{i}}(s)$ between two conformational spaces can then be calculated as a function of the separation between the two particles. Assuming the particles can move freely and homogeneously in their conformational spaces and independently of each other, we can calculate the probability $p_1(s)$ that one particle is at the intersection as

$$p_1 = \frac{v_{\mathrm{i}}}{v_{\mathrm{s}}} \quad (3)$$

and probability $p_2 = p_1^2$ that two particles are at the intersection. This allows us to define a probability density function

$$J(s) = \frac{p_2}{v_{\mathrm{s}}} \quad (4)$$

as the concentration of tethered particles in each other's vicinity. Note that $J$ is expressed in units of concentration, and can therefore be viewed as an effective concentration. With this, we can write an expression for the dimerization probability $f_D$ based on statistical mechanical treatment of the system[18], as

$$f_{\mathrm{D}}(s, K_{\mathrm{D}}) = \frac{J}{J + K_{\mathrm{D}}} \quad (5)$$

This expression was used for the graph shown in **Figure 3d**. The $K_{\mathrm{D}}$ of non-tethered caspase-9 dimerization in solution is not known precisely but reported in the high micromolar range (> 50 µM)[11], and therefore results are shown for several values of $K_{\mathrm{D}}$ in this range.

### Supplementary results

#### Expression and purification of caspase-9 variants

**Supplementary Table 1 | Mutagenesis primers used for generating point mutants C287A and F404D.**

| Mutation | Sequence (5' to 3') |
|---|---|
| C287A | ACCGAAACTGTTTTTCATTCAGGCAGCTGGTGGCGAACAA |
| F404D | CTACAAACAGATGCCGGGCTGTGATAATTTCCTGCGTAAAAAACTG |

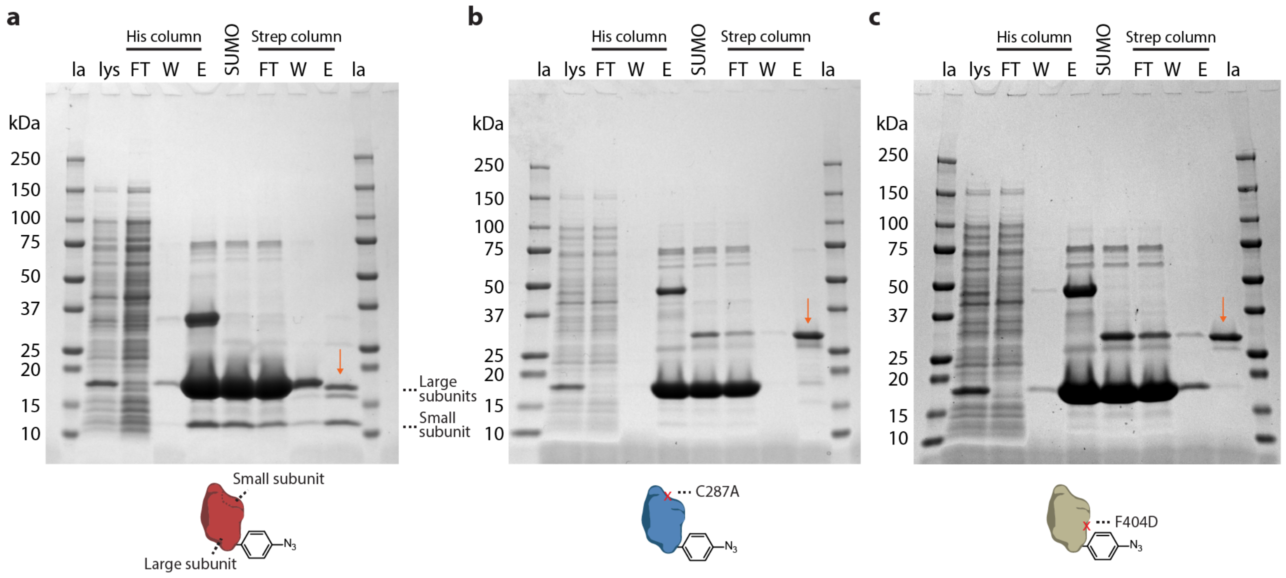


**Supplementary Figure 1 | SDS-PAGE analysis of the expression and purification of all caspase-9 variants.** Images of SDS-PAGE gels (4-20% polyacrylamide, stained for proteins with Coomassie Brilliant Blue), confirming successful purification of wildtype caspase-9 (**a**, expected mass, 18.3 and 19.2 kDa for large subunits, 12.8 kDa for small subunit), and point mutants C287A (**b**, expected mass, 31.9 kDa) and F404D (**c**, expected mass, 31.9 kDa). $Ni^{2+}$-affinity chromatography (His column) and *Strep*-tactin affinity chromatography (Strep column) was performed as described in the Methods section. Enzymatic cleavage of the SUMO-domain from the large subunit of caspase-9 was performed concurrent with dialysis after His column purification. Due to autocatalytic processing during expression, wildtype caspase-9 consists of a non-covalently bound large and small subunit in solution, which are separated during SDS-PAGE under denaturing conditions. Pure protein samples (orange arrows) were aliquoted and stored as described in the Methods section. Labels: la, reference protein ladder (10-250 kDa); lys, cleared lysate after bacterial expression; FT, flow-through fraction; W, wash fraction; E, elution fraction.

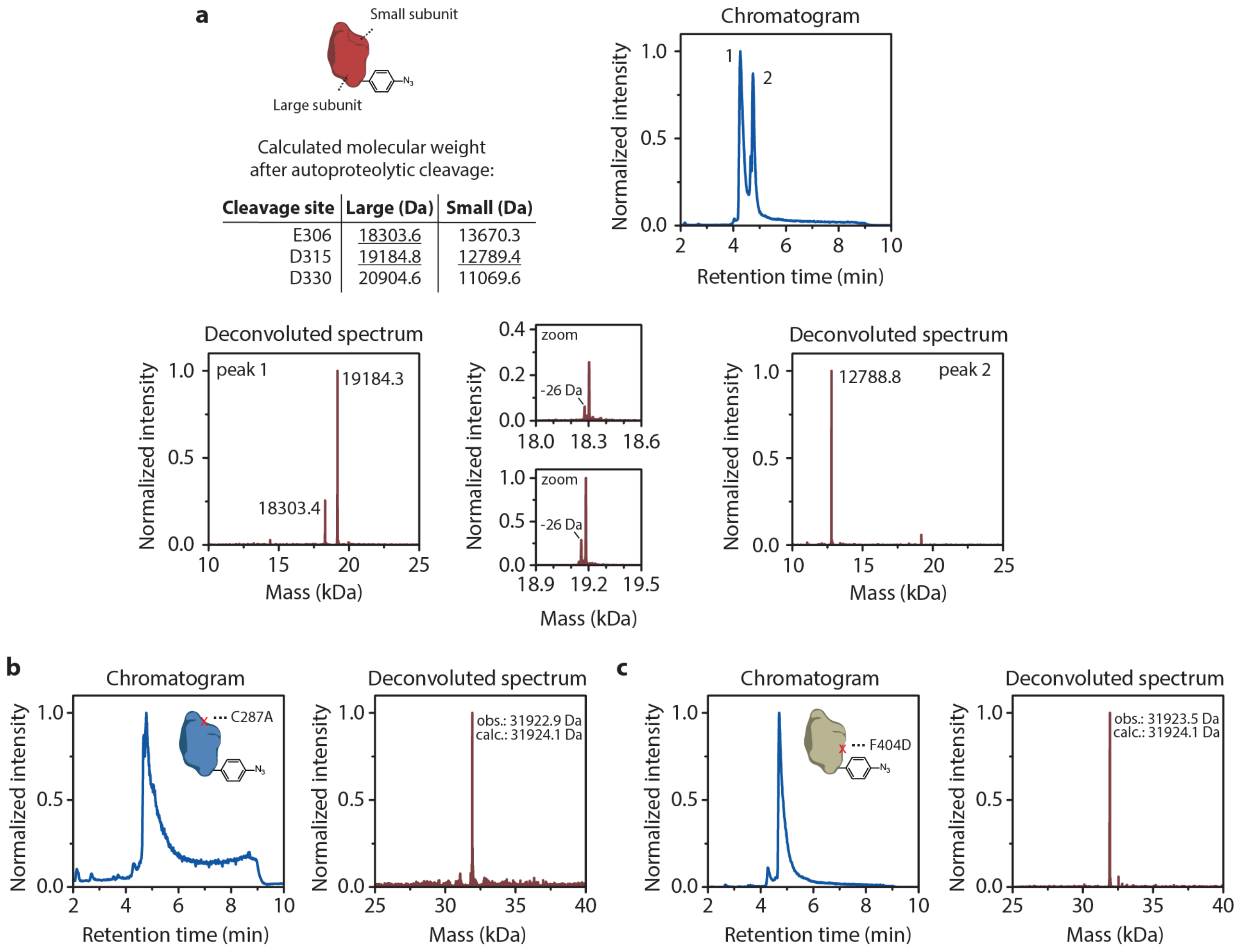


**Supplementary Figure 2 | Liquid chromatography-mass spectrometry (LCMS) analysis of purified caspase-9 variants.** Normalized chromatograms (blue) and deconvoluted mass spectra (red) of wildtype caspase-9 (**a**) and point mutants C287A (**b**) and F404D (**c**). Observed major peaks in all samples confirm the identity of the proteins. Due to autocatalytic processing at three cleavage sites during expression, wildtype caspase-9 consists of non-covalently bound large and small subunits in solution, which are separated during LCMS analysis, leading to the detecting of several protein fragments (**a**). Processing at D315 occurred predominantly and generated large and small subunit fragments of 19.2 and 12.8 kDa, respectively. Minor processing at E306 generated large subunit fragments of 18.3 kDa. Secondary peaks at -26 Da of the main peak can be contributed to the reduction of the azide moiety of unnatural amino acid *p*-azidophenylalanine to an amine in the reducing environment during bacterial expression.

#### Enzyme-DNA conjugation and purification

**Supplementary Table 2 | Anti-handles for enzyme-DNA conjugation.** These oligonucleotides were used for NHS coupling to BCN and subsequent conjugation to caspase-9. Average molecular weight (MW) and extinction coefficient (ε) are indicated. Anti-handle sequences were based on sequences reported in literature with minimal secondary structure and melting temperatures >40°C[19]. All sequences were tested with NUPACK to detect any possible undesired interactions[13]. Underlined thymine nucleotides were added as a spacer. IDT modifications /5AmMC6/ and /3AmMO/ were used for amino modifications at 5' and 3' end, respectively.

| ID | Use | Sequence (5' to 3') | MW (g mol$^{-1}$) | ε (L mol$^{-1}$ cm$^{-1}$) |
|---|---|---|---|---|
| a1 | Wildtype caspase-9, DNA origami and template **T** | $H_2N$-TTTTTTTTTTGAGTGAGTCGTATGA | 7893.2 | 238500 |
| a2 | Wildtype caspase-9, template **T** | CAGTCAGTCAGTCAGTTTTTTTTTT-$NH_2$ | 7830.1 | 230400 |
| a3 | Wildtype caspase-9, DNA origami (longer spacer) | $H_2N$-TTTTTTTTTTTTTTTTGAGTGAGTCGTATGA | 9414.2 | 279000 |
| a4 | Inactive mutants, DNA origami | $H_2N$-TTTTTTTTTTAGTCAGTCAGTCAGT | 7813.2 | 232600 |

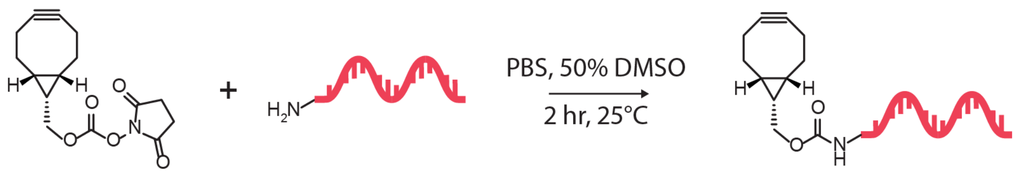


**a1**: $H_2N$-TTTTTTTTTTGAGTGAGTCGTATGA

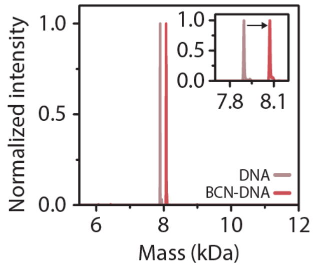


| | Calc. (Da) | Obs. (Da) |
|---|---|---|
| DNA | 7893.2 | 7895.9 |
| BCN-DNA | 8069.4 | 8072.4 |

**a2**: CAGTCAGTCAGTCAGTTTTTTTTTT-$NH_2$

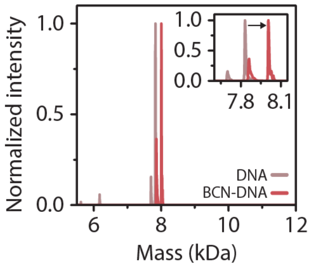


| | Calc. (Da) | Obs. (Da) |
|---|---|---|
| DNA | 7830.1 | 7832.7 |
| BCN-DNA | 8006.3 | 8008.3 |

**a3**: $H_2N$-TTTTTTTTTTTTTTTTGAGTGAGTCGTATGA

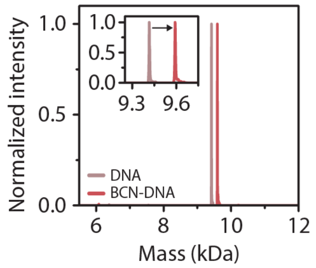


| | Calc. (Da) | Obs. (Da) |
|---|---|---|
| DNA | 9414.2 | 9416.6 |
| BCN-DNA | 9590.4 | 9592.8 |

**a4**: $H_2N$-TTTTTTTTTTAGTCAGTCAGTCAGT

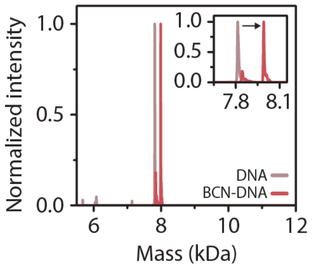


| | Calc. (Da) | Obs. (Da) |
|---|---|---|
| DNA | 7813.2 | 7815.1 |
| BCN-DNA | 7989.4 | 7991.2 |

**Supplementary Figure 3 | MS analysis of oligonucleotides before and after functionalization of the BCN moiety.** Deconvoluted mass spectra of amine-functionalized ODNs a1-a4 used in this study for enzyme-DNA conjugation before (gray) and after (red) NHS coupling to BCN (expected mass increase, 176.21 g mol$^{-1}$). Observed major peaks match the calculated molecular weights for all ODNs. Secondary peaks at -143-145 Da from the main peak could be attributed to depurination caused by heating of the sample during analysis.

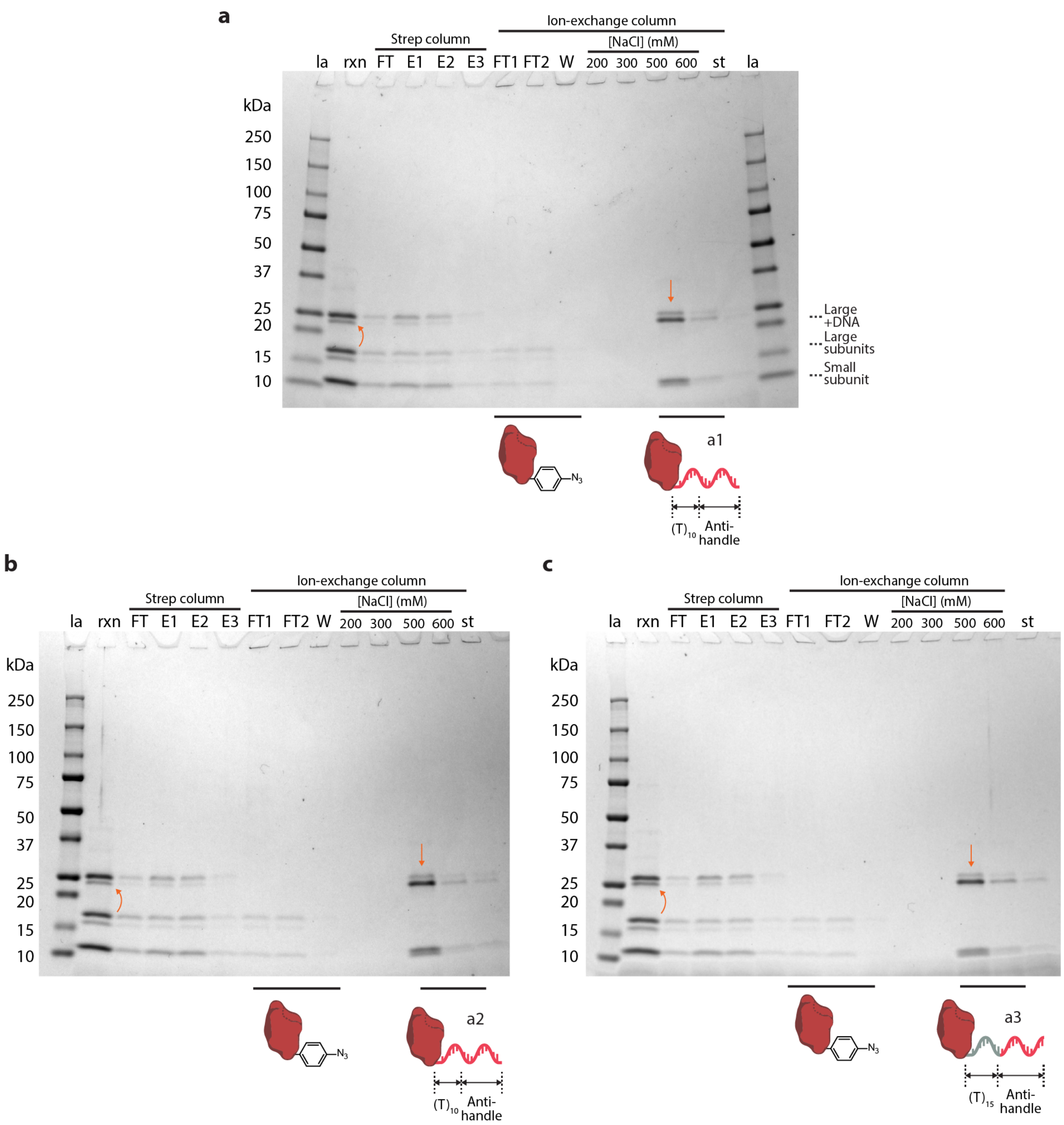


**Supplementary Figure 4 | SDS-PAGE analysis of enzyme-DNA conjugation reactions with wildtype caspase-9.** Images of SDS-PAGE gels (4-20% polyacrylamide, stained for proteins with Coomassie Brilliant Blue), of the conjugation and purification process for wildtype caspase-9 to BCN-functionalized a1 (**a**, ODN mass, 7.9 kDa), a2 (**b,** ODN mass, 7.8 kDa), and a3 (**c**, ODN mass, 9.4 kDa). DNA conjugation to the azide functionality located in the large subunit of caspase-9 results in a selective gel shift of bands corresponding to the large subunits (curved arrows). Subsequent *Strep*-tactin affinity chromatography (Strep column) to remove excess DNA and anion-exchange chromatography (Ion-exchange column) to remove unreacted protein was performed as described in the Methods section. While high concentrations on the ion-exchange column during purification resulted in further autoproteolytic processing of enzyme-DNA conjugates, literature findings suggest that the specific site of processing does not significantly alter the activation behavior of caspase-9[20]. Fractions indicated by straight orange arrows were aliquoted and stored as described in the Methods section. Labels: la, reference protein ladder (10-250 kDa); rxn, crude reaction mixture after overnight conjugation; FT, flow-through fraction; W, wash fraction; E, elution fraction; st, elution after stripping of the column with strip buffer (62.5 mM Tris, 10% glycerol, 50 mM DTT, 2.5% SDS (w/v), 0.01% bromophenol blue, pH 6.8).

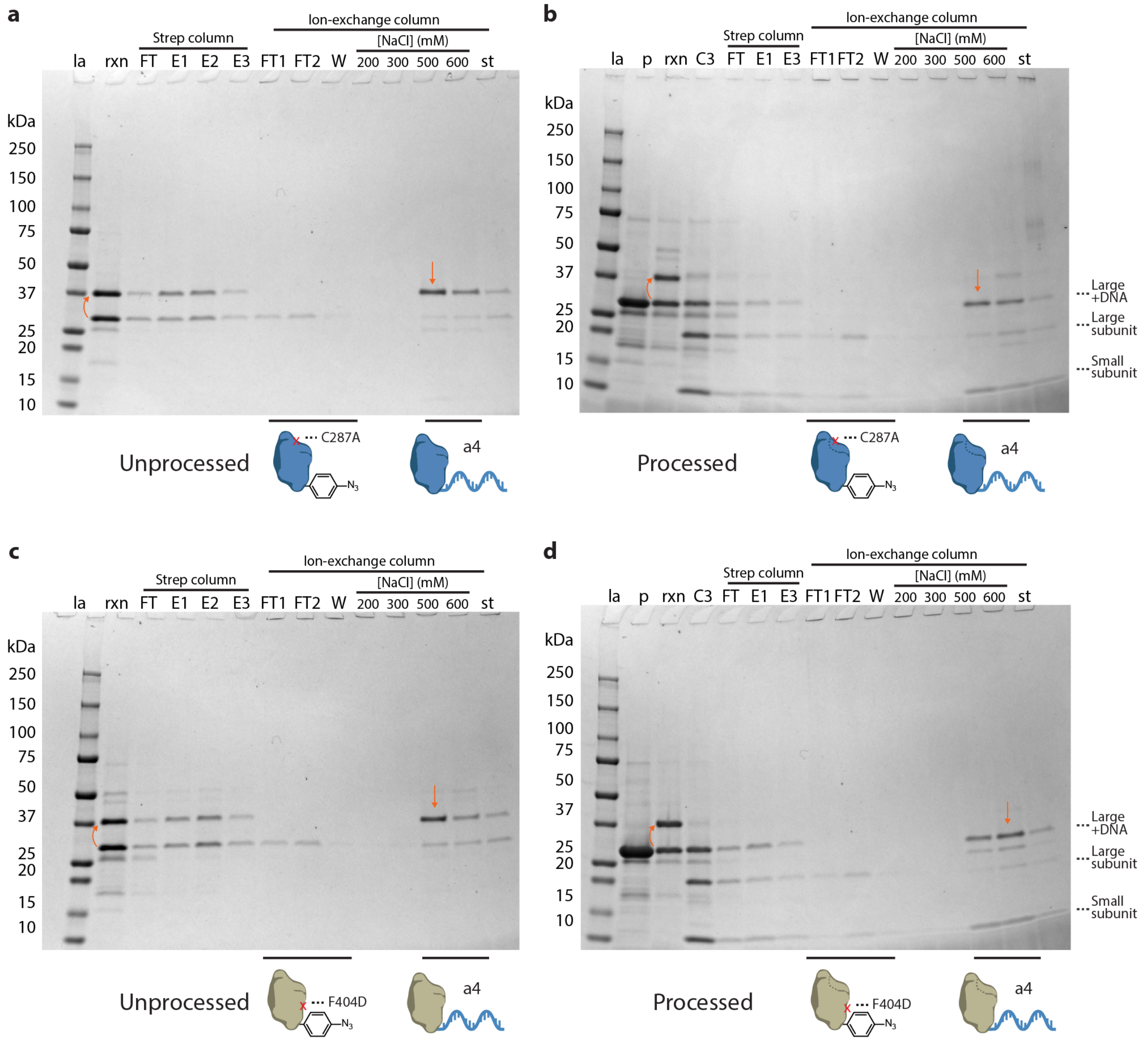


**Supplementary Figure 5 | SDS-PAGE analysis of enzyme-DNA conjugation reactions with caspase-9 point mutants.** Images of SDS-PAGE gels (4-20% polyacrylamide, stained for proteins with Coomassie Brilliant Blue) of the conjugation and purification process for caspase-9 point mutants C287A (**a,b**) and F404D (**c,d**) to BCN-functionalized a4 (ODN mass, 7.8 kDa). To mimic autocatalytic processing in both inactive mutants, processing at D330 was induced by incubating the reaction mixtures with caspase-3 (**b,d**). In all cases, *Strep*-tactin affinity chromatography (Strep column) to remove excess DNA and anion-exchange chromatography (Ion-exchange column) to remove unreacted protein was performed as described in the Methods section. Fractions indicated by straight orange arrows were aliquoted and stored as described in the Methods section. Labels: la, reference protein ladder (10-250 kDa); p, protein only; rxn, crude reaction mixture after overnight conjugation; C3, reaction mixture after processing by caspase-3; FT, flow-through fraction; W, wash fraction; E, elution fraction; st, elution after stripping of the column with strip buffer (62.5 mM Tris, 10% glycerol, 50 mM DTT, 2.5% SDS (w/v), 0.01% bromophenol blue, pH 6.8).

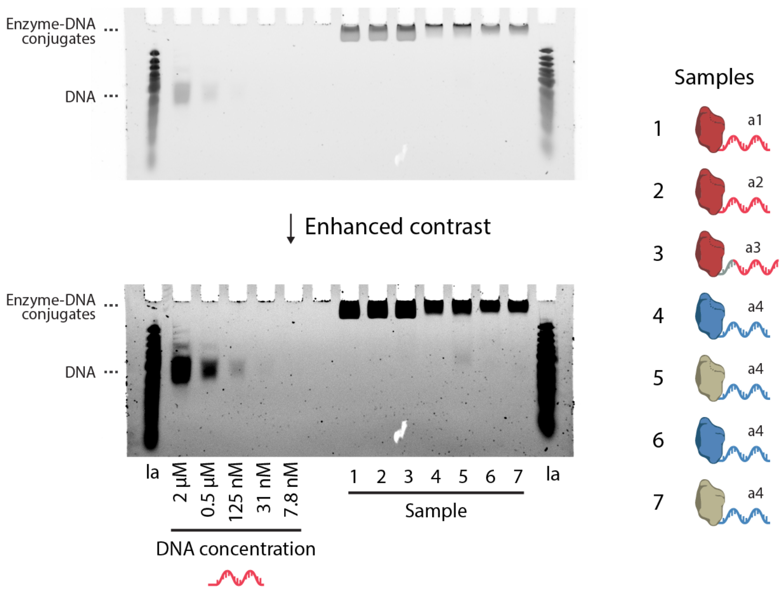


**Supplementary Figure 6 | Assessment of the purity of caspase-9 enzyme-DNA conjugates and removal of excess DNA**. Image and enhanced contrast image of urea-PAGE gel (15% acrylamide, stained for DNA with SYBR Gold) of all enzyme-DNA conjugates (labeled 1-7) used in this study. The purity of the conjugates was assessed using a four-fold dilution series of a 25-nt ssDNA reference, which allows detection of DNA down to a concentration of approximately 31 nM. Since the concentration of the enzyme-DNA conjugates on gel is ~2 µM, it follows that sample 5 contains ≤6% unreacted DNA, while in other samples <2% DNA remains. Label: la, reference dsDNA ladder.

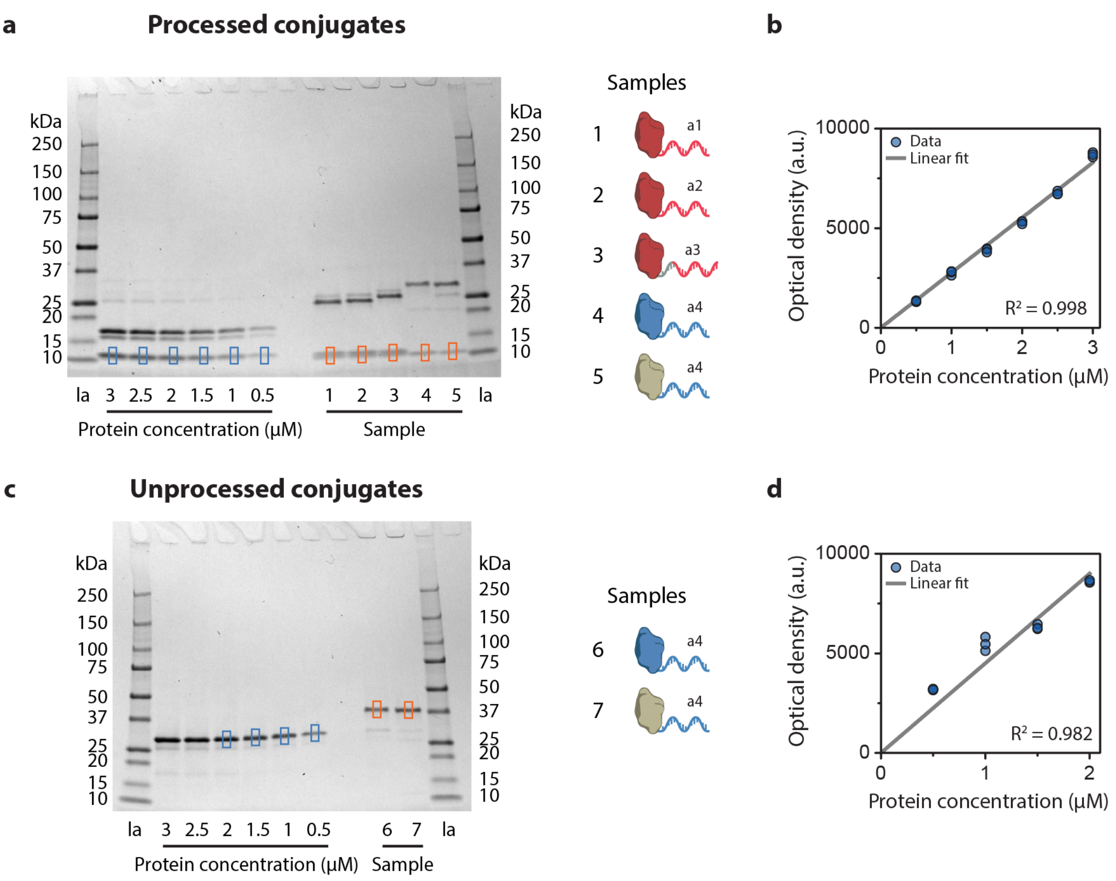


**Supplementary Figure 7 | Determination of the concentration of enzyme-DNA conjugate stocks using optical densitometry.** Typically, the concentration of protein and DNA samples are determined by measuring the absorption at 280 and 260 nm, respectively, which is difficult for enzyme-DNA conjugate samples that contain both species. Gel densitometry provides a straightforward alternative to accurately measure concentrations, and is based on constructing a calibration curve by determining the optical density of gel bands of samples with known concentrations. **a,c,** Images of SDS-PAGE gels (4-20% polyacrylamide, stained for proteins with Coomassie Brilliant Blue) used for gel densitometry of processed (**a**) and unprocessed (**c**) enzyme-DNA conjugates. Dilution series (0.5-3 µM, measured by absorption at 280 nm, as described in the Methods section) of unconjugated wildtype caspase-9 (**a**, small subunit) and point mutant C287A (**c**) were used for calibration. Label: la, reference protein ladder (10-250 kDa). **b,d,** The optical density of the bands indicated by blue squares was determined using the ImageJ gel analysis plugin (the optical density is defined as the area under the curve of linear profile plots perpendicular to the running direction) and fitted to a linear curve. Bands of unprocessed calibration samples at 3 and 2.5 µM (**c**) were overexposed and therefore not taken into consideration in the analysis. The resulting calibration curves were used to determine the concentration of processed (**b**, samples 1-5, orange squares) and unprocessed (**d**, samples 6 and 7, orange squares) enzyme-DNA conjugates. The full analysis procedure was repeated three times.

#### Calibration of substrate LEHD-AFC

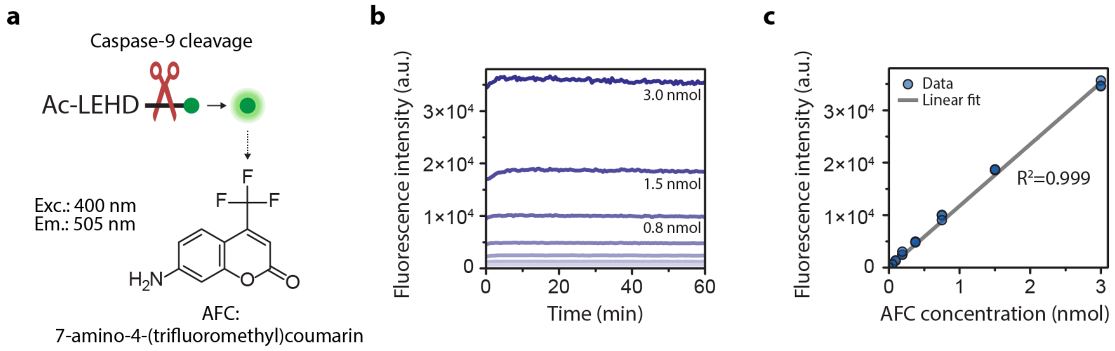


**Supplementary Figure 8 | Concentration calibration of the synthetic caspase-9 substrate LEHD-AFC. a**, Schematic depicting caspase-9 cleavage of the synthetic tetrapeptide LEHD-AFC. Proteolytic activity of caspase-9 leads to cleavage behind the aspartic acid residue, releasing and unquenching the coumarin derivative AFC. The fluorescence increase over time can be measured in a platereader by excitation (ex.) at 400 nm and measuring the emission (em.) at 505 nm. **b**, Graph depicting the fluorescence intensity of free AFC at various concentrations over time (two-fold dilution from a 60 µL 3 nmol stock solution in activity buffer, see Methods section). **c**, A calibration curve was constructed by calculating the mean fluorescence intensity (interval 20-60 min) for each concentration, and fitting the data to a linear curve. Calibrations were performed in triplicate with three different stock solutions of AFC. The concentrations were chosen such that the fluorescence intensities of all experiments fall within the range of the calibration curve.

#### Agarose gel analysis of two-enzyme DNA nanostructures

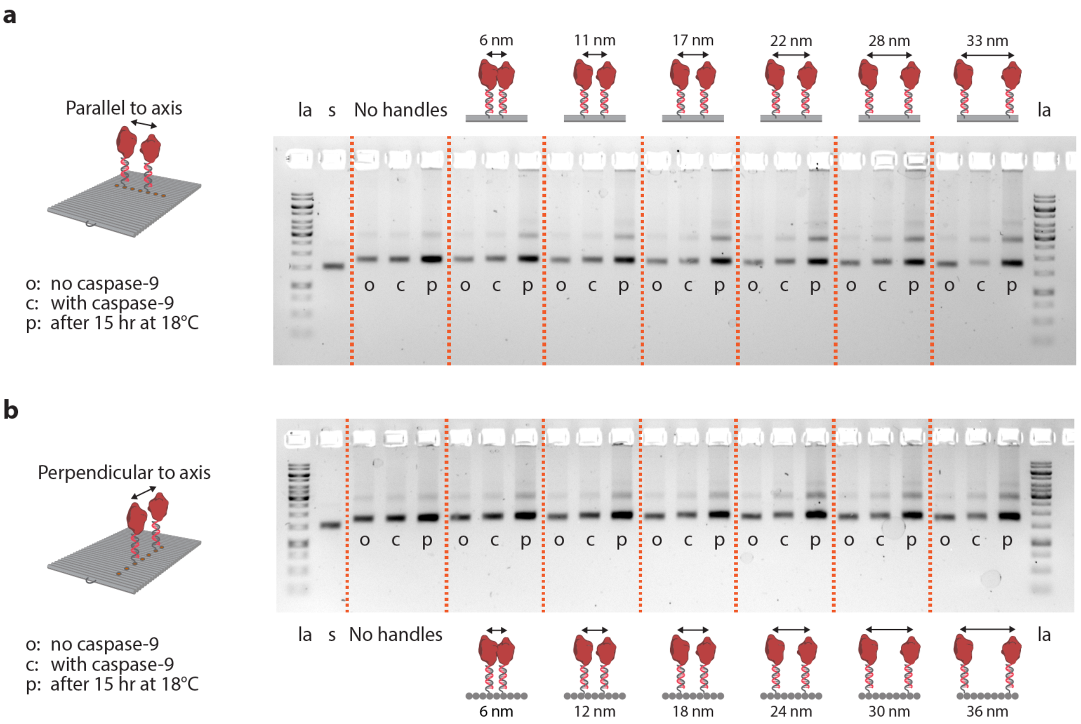


**Supplementary Figure 9 | Structural integrity and stability of two-enzyme DNA nanostructures.** Images of agarose gels (1.5% agarose, stained for DNA with SYBR Safe) of two-enzyme DNA nanostructures carrying two wildtype caspase-9 monomers positioned either parallel (**a**) or perpendicular (**b**) to the DNA helical axis at varying monomer separation distances, as reported in **Figure 3b**. Aliquots of DNA nanostructures (diluted to a DNA origami concentration of 2 nM) were taken before enzyme functionalization (o), after 2 hr incubation with caspase-9 at 4°C (c), and after 15 hr in the platereader at 18°C (p). No significant aggregation or disassembly of the DNA origami nanostructures was observed, suggesting that both the incorporation of caspase-9 and the measurement of enzymatic activity over the course of several hours do not affect the integrity and stability of the DNA nanostructures. Approximately 20% of the reaction volume was evaporated after 15 hr at 18°C, explaining the slightly higher band intensity observed in p samples. Labels: la, reference DNA ladder; s, ssDNA scaffold; no handles, DNA origami without handles for enzyme incorporation.

#### AFM analysis of DNA origami and one-enzyme DNA nanostructures

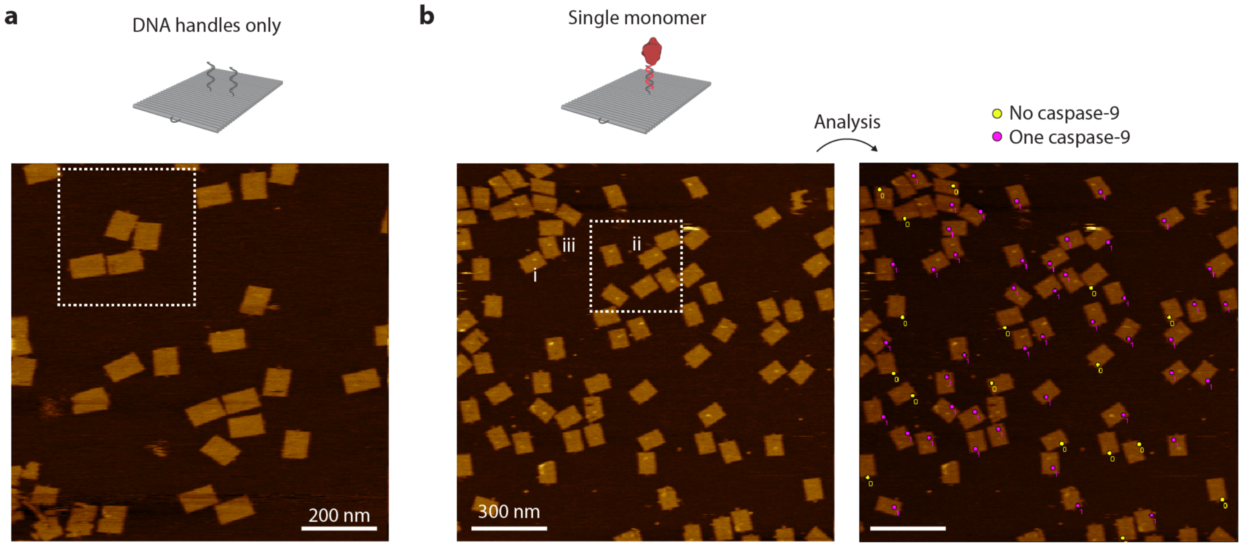


**Supplementary Figure 10 | AFM analysis of no-enzyme and one-enzyme DNA nanostructures. a,** Full-size topographic AFM image of purified DNA origami nanostructures (1.0×1.0 µm², 512×512 px). Dashed rectangle indicates the region used for the image in **Figure 2b**. Scale bar, 200 nm. **b**, Full-size topographic AFM image of one-enzyme DNA nanostructures (1.5×1.5 µm², 512×512 px). Dashed rectangle indicates the region used for the detail image in **Figure 2b**. Roman numerals indicate the individual DNA nanostructures used for the profile plots in **Supplementary Figure 13**. Enzyme incorporation efficiency was determined by manual counting, aided by the ImageJ Multi-point Tool, as described in the Methods section. DNA nanostructures with 0 and 1 enzymes are indicated in the right image with yellow and pink marks, respectively. Scale bars, 300 nm.

#### AFM analysis of two-enzyme DNA nanostructures

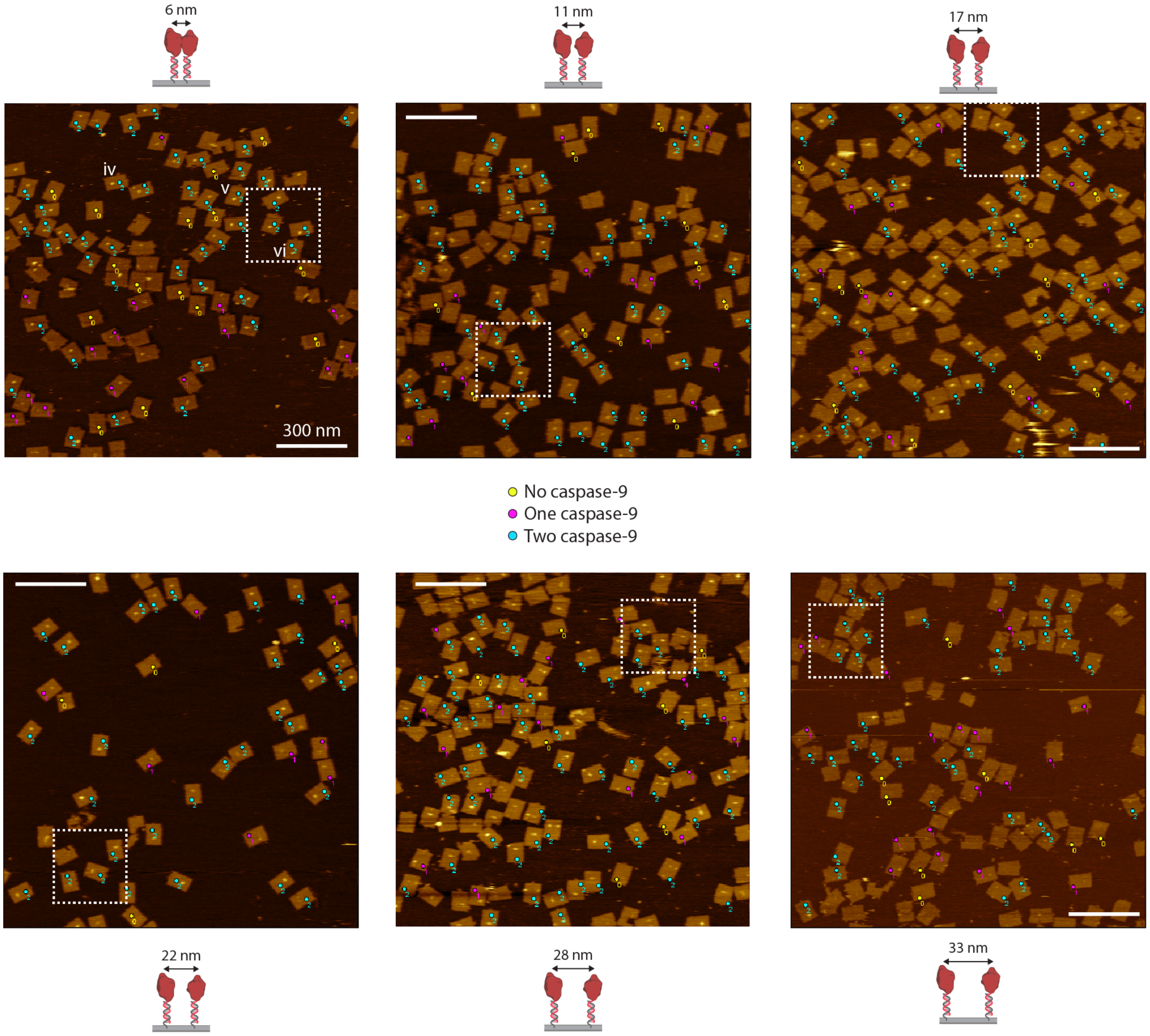


**Supplementary Figure 11 | AFM analysis of two-enzyme DNA nanostructures positioned parallel to the DNA helical axis.** Full-size topographic AFM images of two-enzyme DNA origami nanostructures (1.5×1.5 µm², 512×512 px). Dashed rectangles indicate the regions used for the images in **Figure 2c**. Roman numerals indicate the individual DNA nanostructures used for the profile plots in **Supplementary Figure 13**. Enzyme incorporation efficiency was determined by manual counting, aided by the ImageJ Multi-point Tool, as described in the Methods section. DNA nanostructures with 0, 1, and 2 enzymes are indicated with yellow, pink, and blue marks, respectively. Scale bars, 300 nm.

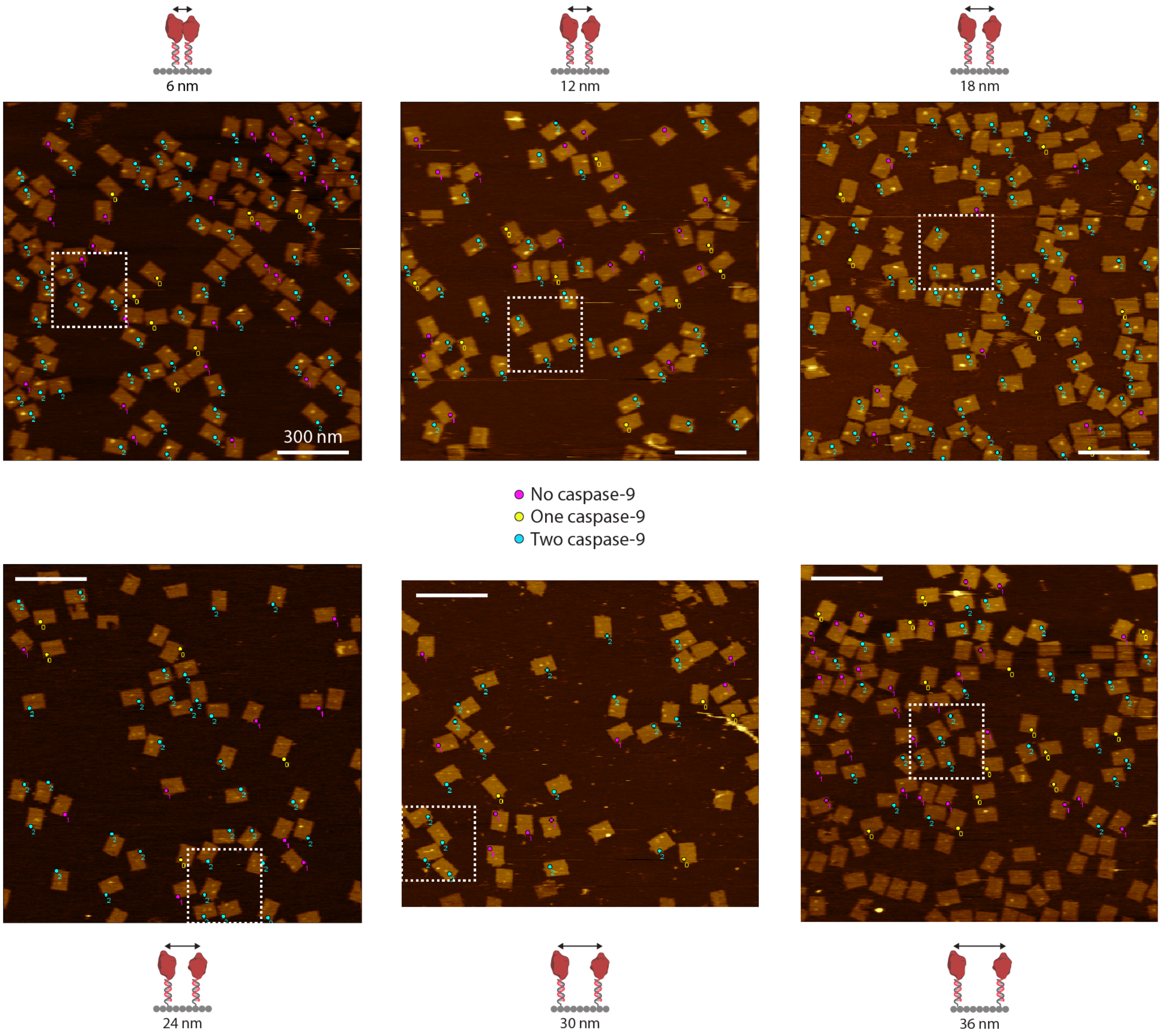


**Supplementary Figure 12 | AFM analysis of two-enzyme DNA nanostructures positioned perpendicular to the DNA helical axis.** Full-size topographic AFM images of two-enzyme DNA origami nanostructures (1.5×1.5 μm², 512×512 px). Dashed rectangles indicate the regions used for the images in **Figure 2c**. Enzyme incorporation efficiency was determined by manual counting, aided by the ImageJ Multi-point Tool, as described in the Methods section. DNA nanostructures with 0, 1, and 2 enzymes are indicated with yellow, pink, and blue marks, respectively. Scale bars, 300 nm.

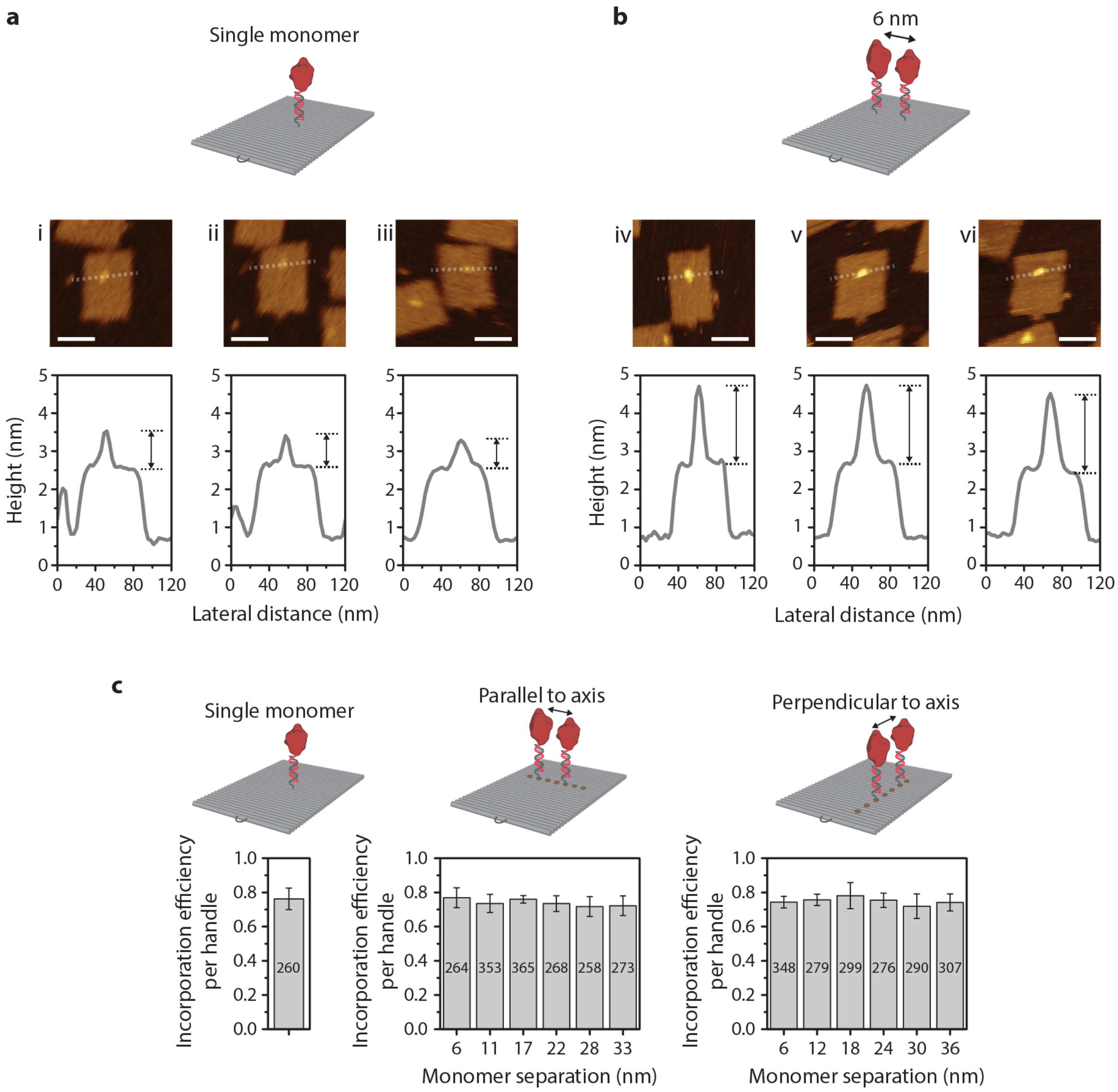


**Supplementary Figure 13 | Comparison of caspase-9 incorporation in one-enzyme and two-enzyme DNA nanostructures. a,b,** Profile plots of individual one-enzyme (**a**) and two-enzyme (**b**) DNA nanostructures based on AFM images of **Supplementary Figure 10** and **Supplementary Figure 11**, illustrating the dimensions of both the DNA origami nanostructure and the enzymes protruding from the surface. The height of the two-dimensional DNA origami corresponds to the diameter of a single DNA double helix, e.g. 2.0-2.5 nm. The clear difference in height of the objects on the surface of the DNA origami in **a** and **b** suggest the presence of two enzymes in the latter. The actual dimensions of the enzymes cannot easily be determined based on these AFM images, due to the translational and rotational freedom of the tethered enzymes and tip-sample convolution effects. Scale bars, 50 nm. **c**, Bar graphs indicating the enzyme incorporation efficiency per handle for one-enzyme and two-enzyme DNA nanostructures. For each sample, the average incorporation efficiency was determined by manual counting of the number of enzymes on well-formed DNA nanostructures (total sample size $n$ indicated in each bar) in at least four different AFM images, as described in the Methods section. Error bars indicate the standard deviation of the incorporation efficiency per image.

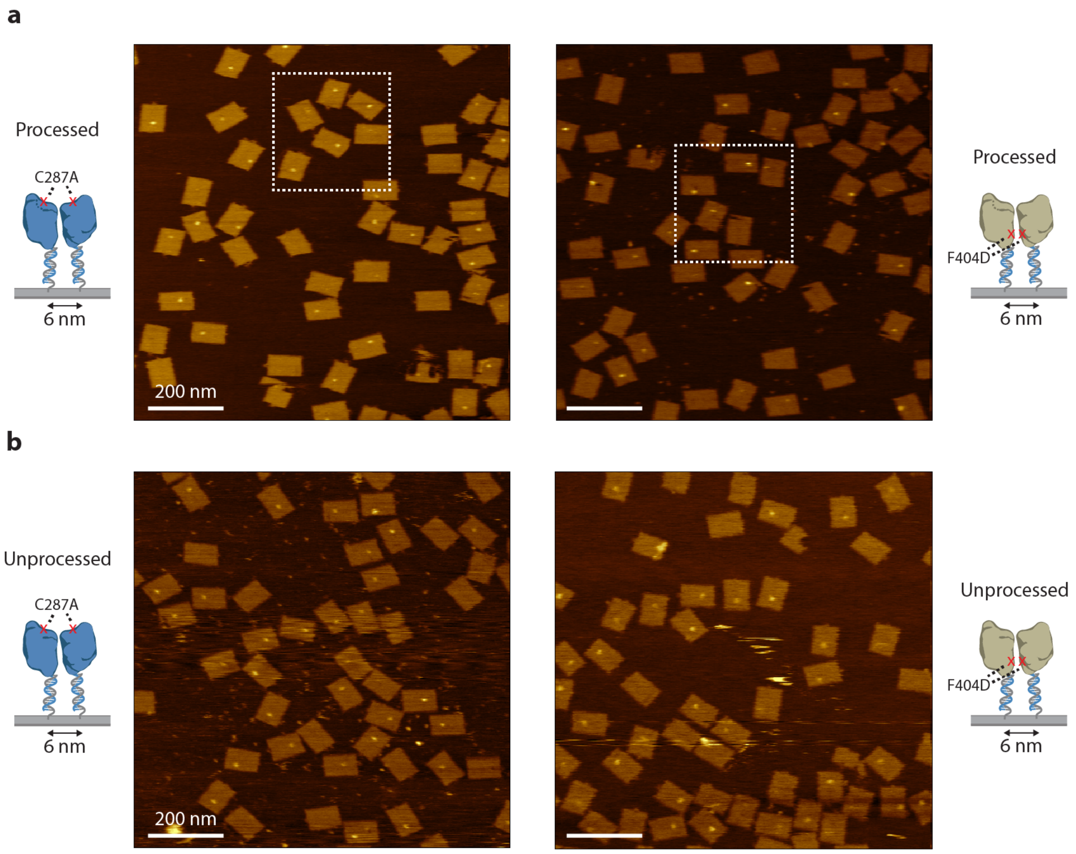


**Supplementary Figure 14 | AFM analysis of two-enzyme DNA nanostructures with caspase-9 point mutants C287A and F404D.** Full-size topographic AFM images of two-enzyme DNA nanostructures (1.0×1.0 µm², 512×512 px) with processed (**a**) and unprocessed (**b**) point mutants C287A and F404D. Dashed rectangles indicate the regions used for the images in **Figure 3e**. Scale bars, 200 nm.

#### Control experiments for two-enzyme DNA nanostructures

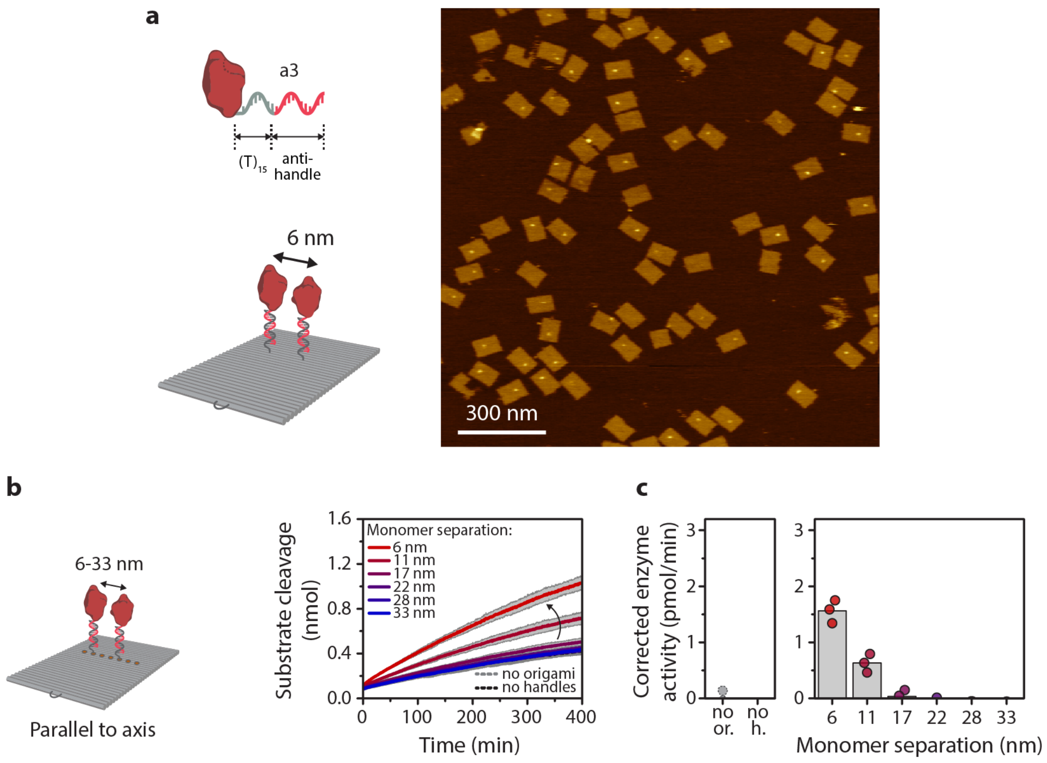


**Supplementary Figure 15 | Analysis of the effect of increased linker length on distance-dependent caspase-9 activity in two-enzyme DNA nanostructures. a**, Full-size topographic AFM image of two-enzyme DNA nanostructures (1.5×1.5 µm$^2$, 512×512 px) using wildtype caspase-9 enzyme-DNA conjugate with an extended ssDNA linker (15 instead of 10 nt) between anti-handle and enzyme. Scale bar, 300 nm. **b**, Distance-dependent enzyme activity of two-enzyme DNA nanostructures with monomers positioned parallel to the DNA helical axis at varying separation distances. Reactions were carried out with 4 nM DNA origami and 3 equivalents of enzyme-DNA conjugate per handle and incubated for 2 hr at 4°C. Activity was determined by monitoring cleavage of 167 µM LEHD-AFC caspase substrate at 18°C. Data is represented as mean activity ± s.d of three independent experiments. **c**, Enzyme activity was determined by taking the initial slope of the kinetic time traces in **b** and corrected by subtracting the mean background activity (no or.). The activity of the 6-nm sample is only 60% of the activity of the equivalent sample reported in **Figure 3b**, suggesting that a longer linker introduces additional rotational and translational flexibility, which adversely affects caspase-9 dimerization and activation. We conclude that the 10-nt ssDNA linker represents an optimal trade-off between necessary flexibility and tight positioning of adjacent monomers to enable an efficient protein-protein interaction. Bars represent mean activity. Labels: no or., no DNA origami present; no h., DNA origami without handles for enzyme incorporation.

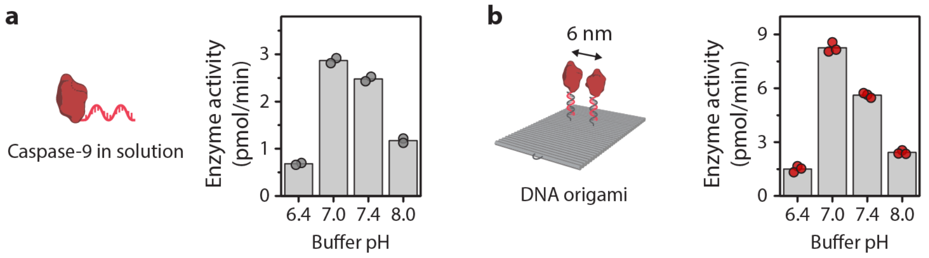


**Supplementary Figure 16 | Comparison of the effect of buffer pH on enzymatic activity of caspase-9 in solution (a) and on 6-nm two-enzyme DNA nanostructures (b).** Reactions were carried out with 24 nM enzyme-DNA conjugate and 0 nM (**a**) or 4 nM (**b**) DNA origami in activity buffer (10 mM Tris, 1 mM EDTA, 10 mM $MgCl_2$, 100 mM NaCl, 1 mM DTT, 0.1% (w/v) CHAPS) at four different pH levels. Activity was determined by monitoring cleavage of 167 µM LEHD-AFC caspase substrate at 18°C. Bars represent mean enzyme activity. Experiments were performed in duplo for **a**, and in triplo for **b**.

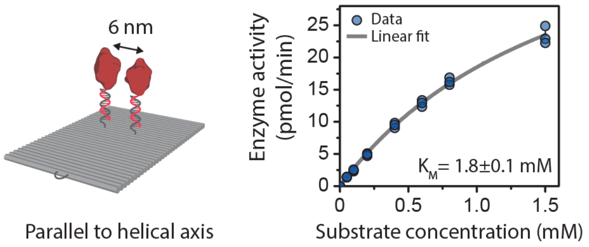


**Supplementary Figure 17 | Kinetic analysis of caspase-9 activity on 6-nm two-enzyme DNA nanostructures.** Reactions were carried out with 4 nM DNA origami and 3 equivalents of enzyme-DNA conjugate per handle and incubated for 2 hr at 4°C. Enzyme kinetics were determined by measuring protease activity at varying concentrations of synthetic tetrapeptide substrate LEHD-AFC, as described in the Methods section. The data was fitted with the standard Michaelis-Menten expression, resulting in a value for the Michaelis constant $K_{\mathrm{M}}$ of 1.8±0.1 mM. Bars represent mean enzyme activity. Experiments were performed in independent triplicates.

#### Geometric model and molecular dynamics simulations

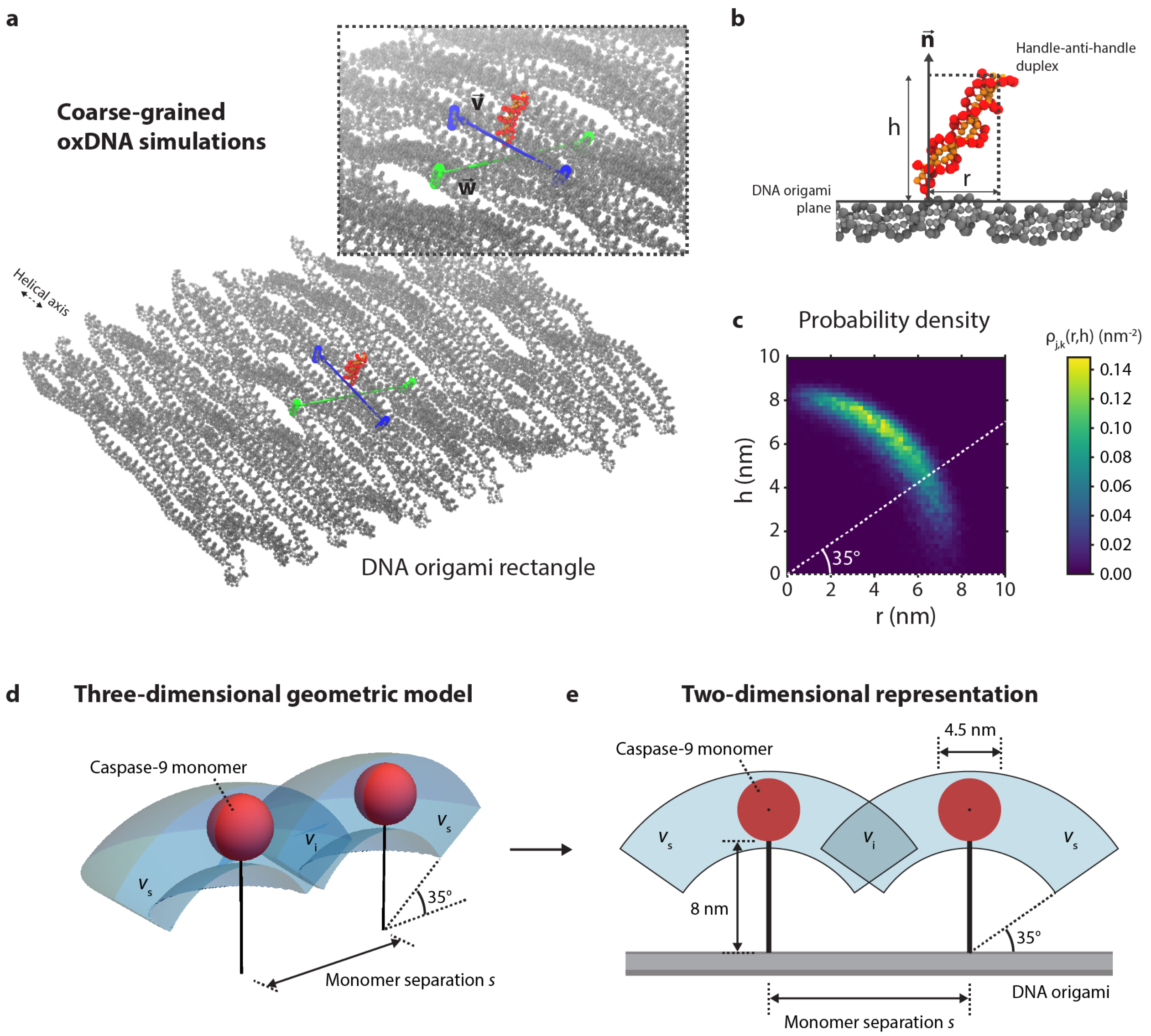


**Supplementary Figure 18 | Overview of the molecular dynamic simulations and geometric model describing tethered caspase-9 dimerization. a**, Snapshot of the coarse-grained molecular dynamics simulation of a DNA origami rectangle (gray) with a single handle-anti-handle duplex protruding from the surface (red and orange). The inset shows the orientation of vectors $\boldsymbol{v}$ (blue) and $\boldsymbol{w}$ (green), which were used to define the DNA origami plane relative to the handle-anti-handle duplex. **b**, Side view of the handle-anti-handle duplex attached to the DNA origami surface. Projection of the position of the end of the duplex onto the DNA origami plane and the normal vector $\boldsymbol{n}$ allows determination of $h$ and $r$, respectively. **c**, Plot of the binned probability density $\rho_{j,k}(r,h)$, calculated using Eq. 2. The simulations suggest that the probability of finding the rigid duplex oriented parallel to the DNA origami plane is low, and that the movement is roughly limited to a volume prescribed by a hemi-shell bounded by an angle of 35° with the DNA origami plane. **d**, Schematic overview of the three-dimensional geometric model, constructed based on the coarse-grained simulations. Each caspase-9 monomer (red) can freely move in a hemi-shell bounded by an angle of 35° with the DNA origami plane with volume $v_\mathrm{s}$ (blue, only half of the volume is displayed for clarity). Two caspase-9 monomers interact when their respective conformational volumes overlap, leading to an intersecting volume $v_\mathrm{i}$ that is a function of the monomer separation $s$. The graphic is drawn to scale and generated using Mathematica (v10, Wolfram). **e**,

Schematic two-dimensional representation of the geometric model. The diameter of a caspase-9 monomer was estimated based on the crystal structure (PDB: 1JXQ), while the length of the tether was determined by the length of the 15-nt handle-anti-handle duplex (15×0.34 nm ≈ 5.1 nm) and the approximate combined length (~3 nm) of the 10-nt ssDNA linker and the alkyl spacer introduced by the amino-functionalization of the ODN. The graphic is drawn to scale.

#### Agarose gel and AFM analysis of three- and four-enzyme DNA nanostructures

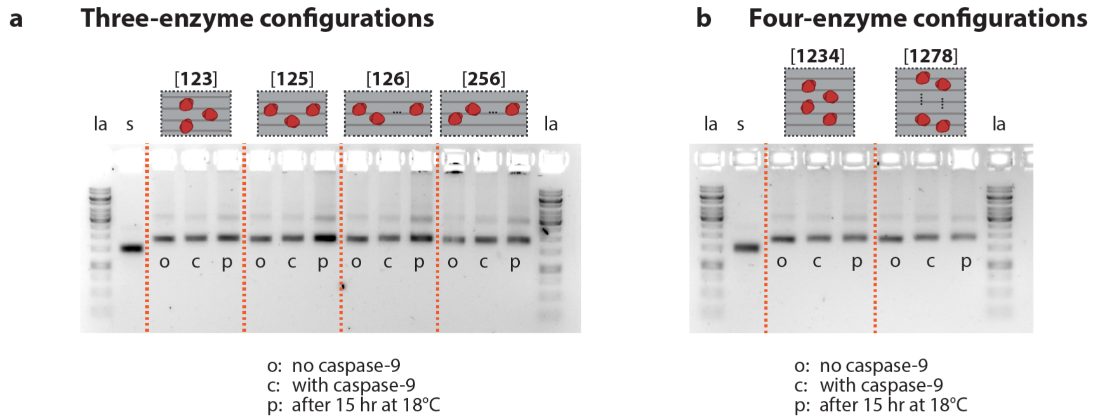


**Supplementary Figure 19 | Structural integrity and stability of three- and four-enzyme DNA nanostructures.** Images of agarose gels (1.5% agarose, stained for DNA with SYBR Safe) of three-enzyme (**a**) and four-enzyme (**b**) DNA nanostructures reported in **Figure 4b**. Aliquots of DNA nanostructures were taken before enzyme functionalization (o), after 2 hr incubation with caspase-9 at 4°C (c), and after 15 hr in the platereader at 18°C (p). All samples were run at a DNA origami concentration of 2 nM. No significant aggregation or disassembly of the DNA origami was observed, suggesting that both the incorporation of caspase-9 and the measurement of enzymatic activity over the course of several hours do not affect the integrity and stability of the DNA nanostructures. Labels: la, reference DNA ladder; s, ssDNA scaffold.

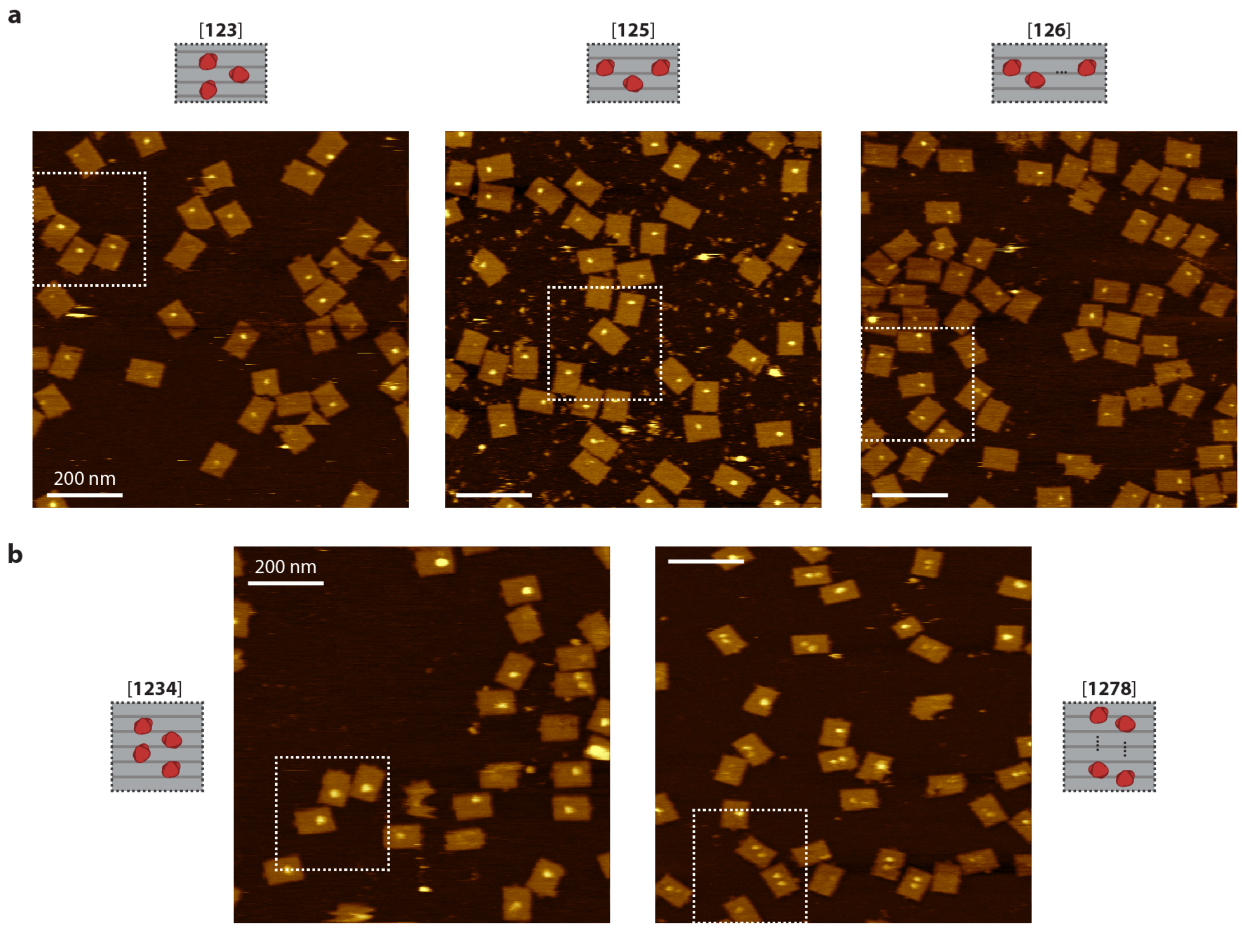


**Supplementary Figure 20 | AFM analysis of three- and four-enzyme DNA nanostructures.** Full-size topographic AFM images of three- (**a**) and four-enzyme (**b**) DNA nanostructures (1.0×1.0 μm², 512×512 px). Dashed rectangles indicate the regions used for the images in **Figure 4a**. Scale bars, 200 nm.

#### Agarose gel and AFM analysis of heterodimer DNA nanostructures

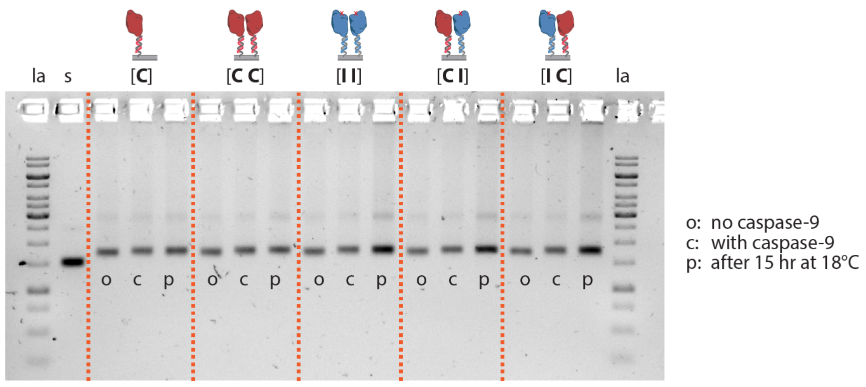


**Supplementary Figure 21 | Structural integrity and stability of heterodimer two-enzyme DNA nanostructures.** Images of agarose gels (1.5% agarose, stained for DNA with SYBR Safe) of DNA nanostructures reported in **Figure 5e**. Aliquots of DNA nanostructures were taken before enzyme functionalization (o), after

2 hr incubation with caspase-9 at 4°C (c), and after 15 hr in the platereader at 18°C (p). All samples were run at a DNA origami concentration of 2 nM. No significant aggregation or disassembly of the DNA origami was observed, suggesting that both the incorporation of caspase-9 and the measurement of enzymatic activity over the course of several hours do not affect the integrity and stability of the DNA nanostructures. Labels: la, reference DNA ladder; s, ssDNA scaffold.

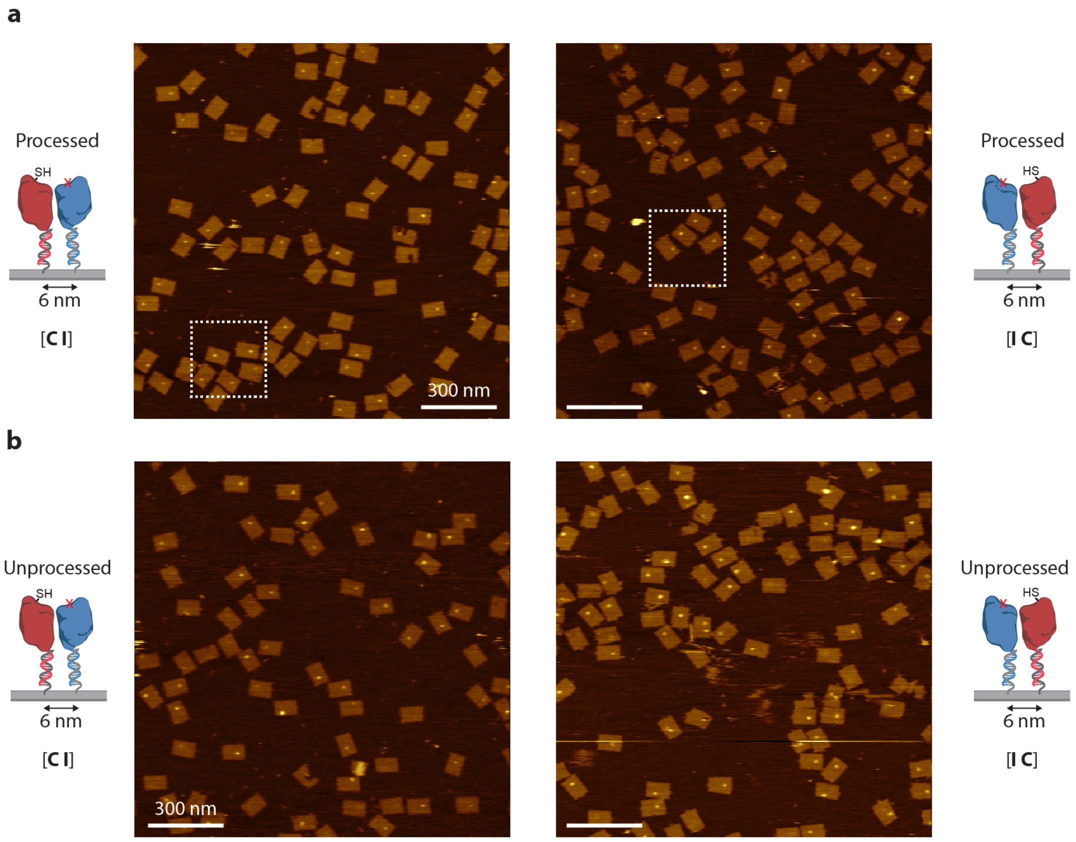


**Supplementary Figure 22 | AFM analysis of heterodimeric two-enzyme DNA nanostructures.** Full-size topographic AFM images of heterodimeric two-enzyme DNA nanostructures (1.5×1.5 µm², 512×512 px) with processed (**a**) and unprocessed (**b**) point mutant C287A. Dashed rectangles indicate the regions used for the images in **Figure 5d**. Scale bars, 300 nm.

#### Activity of unprocessed heterodimer DNA nanostructures

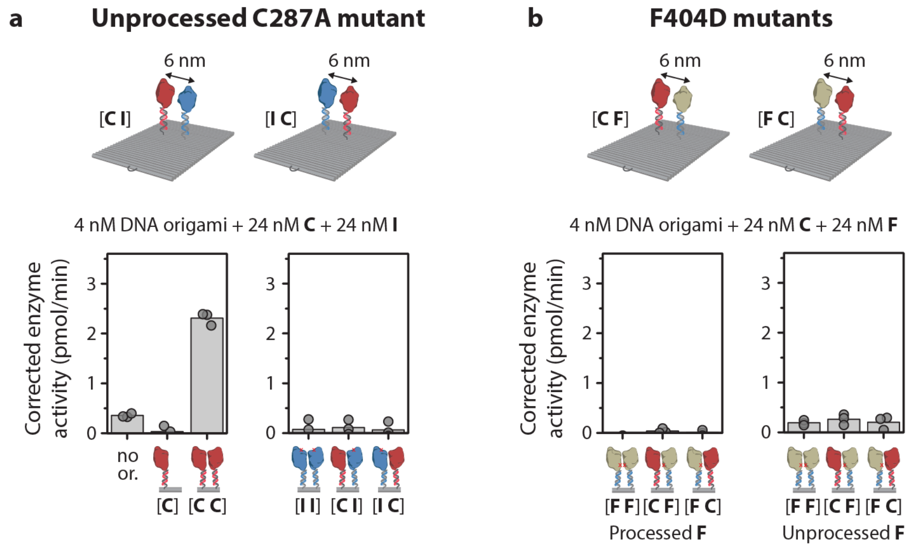


**Supplementary Figure 23 | Activity assays of heterodimer two-enzyme DNA nanostructures using unprocessed C287A caspase-9 mutant (a) and F404D mutants (b).** Reactions were carried out with 4 nM DNA origami and 24 nM of both wildtype (**C**) and inactive (i.e. unprocessed **I**, unprocessed **F**, or processed **F**) enzyme-DNA conjugate per handle and incubated for 2 hr at 4°C. Enzyme configurations are denoted using bracket notation, as reported in **Figure 5**. Activity was determined by monitoring cleavage of 167 µM LEHD-AFC caspase-9 substrate over time at 18°C. Bars represent mean corrected enzyme activity. Experiments were repeated in triplicate. Label: no or., no DNA origami present.

### Derivation of the thermodynamic models

#### Section 1: dimerization of tethered enzymes

We consider the dimerization of caspases that are tethered on a DNA origami scaffold via linkers at a certain separation distance. Each enzyme can be present in monomeric form or as part of a dimer. In this section we will derive the average number of dimers per DNA origami for 5 different enzyme configurations, as shown in **Supplementary Figure 24**, i.e. (Case 1) a DNA origami with 2 caspases with the linkers attached at a distance ℓ, (Case 2) a DNA origami with 2 caspases with the linkers attached at a distance 2ℓ, (Case 3) a DNA origami with 3 caspases with the linkers attached in a linear geometry with neighbors at a distance ℓ apart, (Case 4) a DNA origami with 3 caspases with the linkers attached in a triangular geometry all at distance ℓ apart, and (Case 5) a DNA origami with 4 caspases connected in a square geometry with sides of length ℓ.

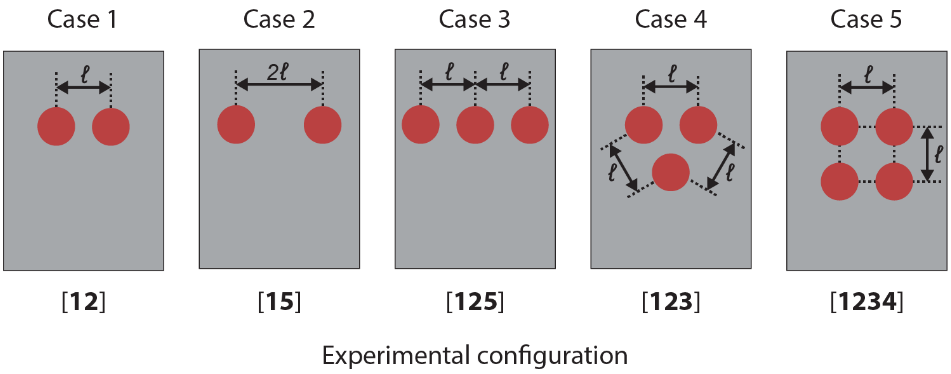


**Supplementary Figure 24 |** Schematics of the five distinct DNA origami scaffolds considered in the analysis in this section, where the number and geometry of linkers to which caspases can be bound differ.

**Case 1: two caspases connected with a linker to a scaffold at distance ℓ.** The first case we consider is that of dimerization of two caspases (C) that are fixed to a DNA-origami scaffold via two linkers at a certain separation. The two caspases can be present as two monomers or as a dimer. Because of the co-localization of the caspases by the DNA origami, the dimerization is different from dimerization of non-tethered caspases, where the fraction dimer is concentration dependent. Here, we should consider the dimerization reaction as two possible states with (concentration independent) rate constants (probabilities) that a dimer is formed or broken, i.e.,

$$\mathbf{C} + \mathbf{C} \underset{k_\ell^-\,(\mathrm{s}^{-1})}{\overset{k_\ell^+\,(\mathrm{s}^{-1})}{\rightleftarrows}} \mathbf{C\text{-}C}$$

where both $k_\ell^+$ and $k_\ell^-$ have unit $\mathrm{s}^{-1}$. The two states (monomers and dimer) are schematically depicted in **Supplementary Figure 25**.

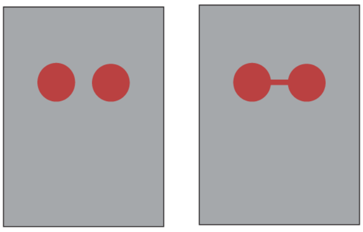


**Supplementary Figure 25 |** The two-enzyme configuration with linkers at a distance $\ell$ apart can be in 2 distinct states, i.e. monomeric (state 1, also denoted as 2M) and dimeric (state 2, also denoted as D).

We define the fraction of DNA origami with the caspases in dimeric state as $f_{\mathrm{D}}$ and the fraction with the caspases in monomeric state as $f_{2\mathrm{M}}$. Because the caspases on the DNA origami should either be in the monomeric state or form a dimer, it holds that

$$f_{\mathrm{D}} + f_{2\mathrm{M}} = 1. \tag{6}$$

Moreover, in equilibrium, the rates of dimerization and dimer dissociation should be equal, i.e.,

$$f_{2\mathrm{M}} k_\ell^+ = f_{\mathrm{D}} k_\ell^- . \tag{7}$$

With equilibrium constant

$$K_\ell = \frac{k_\ell^+}{k_\ell^-} \tag{8}$$

Eq. (7) can be rewritten as

$$f_{\mathrm{D}} = K_\ell f_{2\mathrm{M}} . \tag{9}$$

Substituting Eq. (6) in Eq. (9) yields

$$f_{\mathrm{D}} = K_\ell (1 - f_{\mathrm{D}}) \tag{10}$$

which can be rewritten as

$$f_{\mathrm{D}} = \frac{K_\ell}{1+K_\ell} \tag{11}$$

Analogously, the fraction of DNA origami with caspases in monomeric state as function of $K_\ell$ is given by

$$f_{2\mathrm{M}} = \frac{1}{1+K_\ell} . \tag{12}$$

The average number of dimers per DNA origami for this proximal two-enzyme configuration is simply $f_{\mathrm{D}}$. In the remainder of this text we will derive expressions for the average number of dimers per DNA origami for the other enzyme configurations as a function of this $f_{\mathrm{D}}$, which we denote as the *dimerization probability*.

**Case 2: two caspases connected with a linker to a scaffold at distance 2ℓ.** The second case we consider is similar to Case 1, but with caspases connected to the scaffold with a separation of $2\ell$ instead of $\ell$. Here, we again have two distinct states, as depicted in **Supplementary Figure 26**, so we again have

$$f_{\mathrm{D}_{2\ell}} + f_{2\mathrm{M}_{2\ell}} = 1 \tag{13}$$

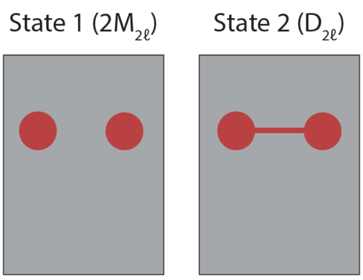


**Supplementary Figure 26 |** The two-enzyme configuration with monomers at a distance $2\ell$ can be in 2 distinct states, i.e. monomeric (state 1, also denoted as $2\mathrm{M}_{2\ell}$) and dimeric (state 2, also denoted as $\mathrm{D}_{2\ell}$).

When the distance between the linkers is 11 or 12 nm the activity is lower than at 6 nm. Therefore, we assume that the dimerization probability for 2 caspases at distance $2\ell$ apart will be a factor $a$ smaller than for 2 caspases at distance $\ell$ apart, i.e., $f_{\mathrm{D}_{2\ell}} = af_{\mathrm{D}}$, with $0 \leq a \leq 1$. The corresponding equilibrium constant $K_{2\ell}$ for this dimerization of these more distal caspases is related to $K_\ell$ as

$$K_{2\ell} = aK_\ell/(1 + (1 - a)K_\ell). \tag{14}$$

For the equilibrium condition now holds that

$$f_{\mathrm{D}_{2\ell}} = K_{2\ell} f_{2\mathrm{M}_{2\ell}} = \frac{aK_\ell}{1+(1-a)K_\ell} f_{2\mathrm{M}_{2\ell}}. \tag{15}$$

Substituting in Eq. (13), we can determine that the probability of finding the caspases in the monomeric state as function of $K_\ell$ is given by

$$f_{2\mathrm{M}_{2\ell}} = \frac{1+K_\ell(1-a)}{1+K_\ell} = 1 - \frac{aK_\ell}{1+K_\ell} = 1 - af_{\mathrm{D}}. \tag{16}$$

The average number of dimers per DNA origami for this more distal two-enzyme configuration is thus $af_{\mathrm{D}}$.

**Case 3: three caspases connected with a linker to a scaffold in linear geometry.** We now consider a DNA origami with three caspases connected to it that are placed in a linear fashion, where we assume that two caspases at a distance $\ell$ can dimerize with dimerization probability $f_{\mathrm{D}}$, and that the outer caspases (at a distance $2\ell$) can dimerize with the lower probability $af_{\mathrm{D}}$, as in Case 2. Such a DNA origami can then be in four distinct states, which are schematically depicted in **Supplementary Figure 27**. In the first state, which we denote as 3M, all caspases are present in monomeric form. In the second state, which we denote as $D_{12}$, the left and middle caspases form a dimer while the right caspase is present as a monomer. In the third state, which we denote as $D_{23}$, the left caspase is present as a monomer while the middle and right caspases form a dimer. In the fourth state, which we denote as $D_{13}$, the middle caspase is present as a monomer while the left and right caspases form a dimer.

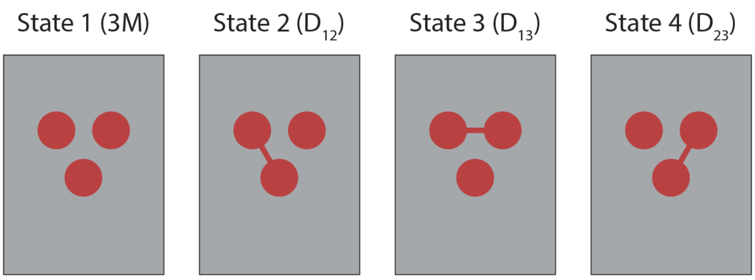


**Supplementary Figure 27 |** The linear three-enzyme configuration, assuming that also the two outer caspases can dimerize together, can be in 4 different states, i.e. monomeric state (3M) and three distinct dimeric states ($D_{12}$, $D_{23}$ and $D_{13}$).

If we denote the fraction of such DNA origami in a certain state $X$ by $f_X$, we can write

$$f_{3M} + f_{D_{12}} + f_{D_{23}} + f_{D_{13}} = 1 \tag{17}$$

as each origami should be in either of the 4 states. For states $D_{12}$ and $D_{23}$ it holds, in equilibrium, that

$$f_{D_{12}} = f_{D_{23}} = K_\ell f_{3M}\,, \tag{18}$$

whereas for the dimer between the 2 outer caspases, similarly as in Case 2, Eq. (15) holds that

$$f_{D_{13}} = K_{2\ell} f_{3M} = \frac{aK_\ell}{1+(1-a)K_\ell} f_{3M} \tag{19}$$

Substituting Eqs. (18) and (19) into Eq. (17) yields for the fraction of DNA origami with the caspases in monomeric form

$$f_{3M} = \frac{1}{1+\left(2+\frac{a}{1+(1-a)K_\ell}\right)K_\ell}\,. \tag{20}$$

For states $D_{12}$ and $D_{23}$ we thus have

$$f_{D_{12}} = f_{D_{23}} = \frac{K_\ell}{1+\left(2+\frac{a}{1+(1-a)K_\ell}\right)K_\ell}\,. \tag{21}$$

and for state $D_{13}$

$$f_{D_{13}} = \frac{aK_\ell}{1+\left(2+\frac{a}{1+(1-a)K_\ell}\right)K_\ell}\,. \tag{22}$$

Eqs. (20)-(22) can also be expressed in terms of $f_D$, i.e. the dimerization probability for the proximal two-enzyme configuration as described in Case 1, instead of $K_\ell$ as according to Eq. (10) holds that $K_\ell = f_D/(1 - f_D)$. This yields:

$$f_{3M} = \frac{1}{1+\left(2+\frac{a}{1+(1-a)\frac{f_D}{1-f_D}}\right)\frac{f_D}{1-f_D}} = \frac{(1-f_D)(1-af_D)}{1+f_D-2af_D{}^2}, \tag{23}$$

$$f_{D_{12}} = f_{D_{23}} = \frac{\frac{f_D}{1-f_D}}{1+\left(2+\frac{a}{1+(1-a)\frac{f_D}{1-f_D}}\right)\frac{f_D}{1-f_D}} = \frac{f_D(1-af_D)}{1+f_D-2af_D{}^2} \tag{24}$$

and

$$f_{D_{13}} = \frac{\frac{a}{1+(1-a)\frac{f_D}{1-f_D}}\left(\frac{f_D}{1-f_D}\right)}{1+\left(2+\frac{a}{1+(1-a)\frac{f_D}{1-f_D}}\right)\frac{f_D}{1-f_D}} = \frac{a(1-f_D)\,f_D}{1+f_D-2af_D{}^2}\,. \qquad (25)$$

**Case 4: three caspases connected with a linker to a scaffold in triangular geometry.** If the three caspases are placed in a triangular geometry, four different states exist, namely 1 state where all 3 caspases are present as monomers and three states for the three different possible dimers, as depicted in **Supplementary Figure 28**.

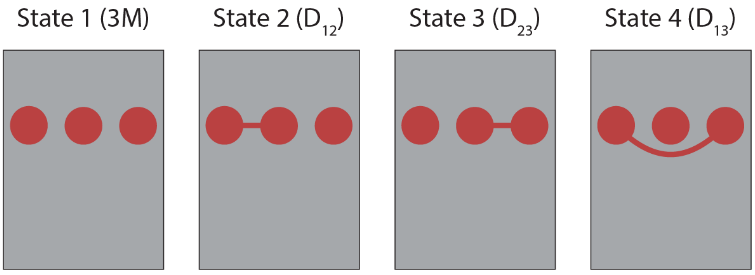


**Supplementary Figure 28 |** The triangular three-enzyme configuration can be in 4 different states, i.e. monomeric state (3M) and three distinct dimeric states ($D_{12}$, $D_{13}$ and $D_{23}$).

As each DNA origami should again be in either of the 4 states, we have

$$f_{3M} + f_{D_{12}} + f_{D_{13}} + f_{D_{23}} = 1 \qquad (26)$$

which together with $f_{D_{12}} = f_{D_{13}} = f_{D_{23}} = f_{3M}K_\ell$ yields

$$f_{3M} = \frac{1}{1+3K_\ell} \quad \text{and} \quad f_{D_{12}} = f_{D_{13}} = f_{D_{23}} = \frac{K_\ell}{1+3\ell}\,. \qquad (27)$$

Expressed in terms of $f_D$ (dimerization probability for the proximal two-enzyme configuration) that is

$$f_{3M} = \frac{1}{1+\frac{3f_D}{1-f_D}} = \frac{1-f_D}{1+2f_D} \qquad (28)$$

and

$$f_{D_{12}} = f_{D_{13}} = f_{D_{23}} = \frac{\frac{f_D}{1-f_D}}{1+\frac{3f_D}{1-f_D}} = \frac{f_D}{1+2f_D}\,. \qquad (29)$$

Note that for $a = 1$ Eqs. (23)-(25) reduce to Eqs. (28) and (29).

**Case 5: four caspases connected with a linker to a scaffold in square geometry.** In the case of 4 caspases in a proximal configuration, where dimers can be formed between any pair of caspases, the number of different states for the DNA origami consists of one state where all caspases are present as monomer, 4 distinct states with a single dimer between 2 not-diagonal neighboring caspases and two monomers, 2 distinct states with a single dimer between 2 diagonally neighboring caspases and two monomers, and 2 distinct states with 2 dimers (**Supplementary Figure 29**). The state where two pairs of diagonally neighboring caspases both dimerize is ignored as the first diagonally dimerized pair will prevent formation of the second dimer.

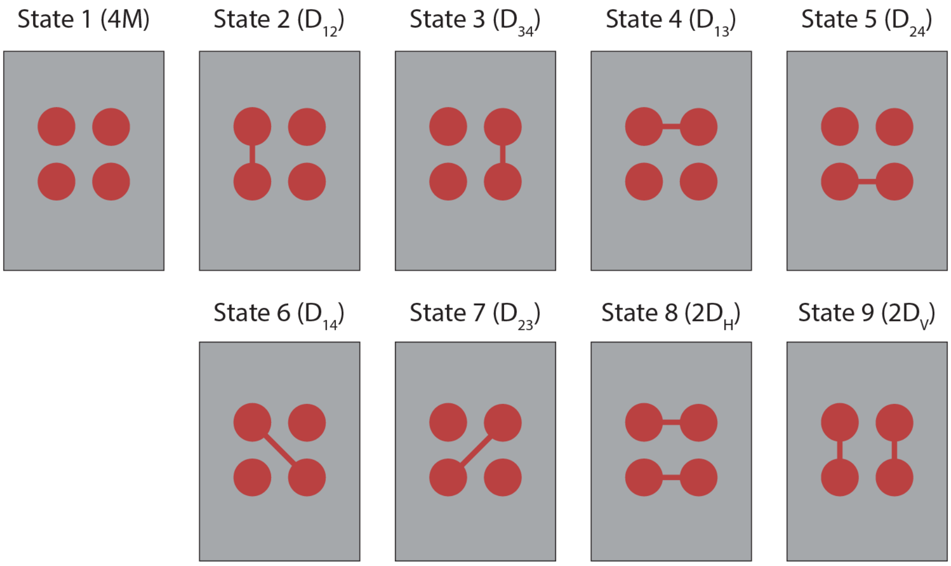


**Supplementary Figure 29 |** The proximal four-enzyme configuration can be in 9 different states, i.e. monomeric state (4M), six distinct single dimeric states ($D_{12}$, $D_{34}$, $D_{13}$, $D_{24}$, $D_{14}$ and $D_{23}$), and two distinct double dimeric states ($2D_H$ and $2D_V$).

As each DNA origami should again be in either of the different possible states, we have

$$f_{4\mathrm{M}} + f_{\mathrm{D}_{12}} + f_{\mathrm{D}_{34}} + f_{\mathrm{D}_{13}} + f_{\mathrm{D}_{24}} + f_{\mathrm{D}_{14}} + f_{\mathrm{D}_{23}} + f_{2\mathrm{D_H}} + f_{2\mathrm{D_V}} = 1 \tag{30}$$

For the equilibria between the state with all monomers and the states with a single dimer between horizontally or vertically neighboring caspases, we again obtain $f_{\mathrm{D}_{12}} = f_{\mathrm{D}_{34}} = f_{\mathrm{D}_{13}} = f_{\mathrm{D}_{24}} = K_\ell f_{4\mathrm{M}}$. Moreover, the equilibrium between states with two dimers and such a single dimer state yields

$$f_{2\mathrm{D_V}} = K_\ell f_{\mathrm{D}_{12}} = K_\ell{}^2 f_{4\mathrm{M}}\,. \tag{31}$$

As the diagonally neighboring caspases are at slightly larger distance apart, we assume again that their dimerization probability is a factor $b$ smaller than for the regular dimer, i.e., $f_{\mathrm{D}_{14}} = f_{\mathrm{D}_{23}} = bf_\mathrm{D}$ with $0 \le b \le 1$, which corresponds to equilibrium constants

$$K_{14} = K_{23} = bK_\ell/(1 + (1-b)K_\ell)\,. \tag{32}$$

Thus,

$$f_{\mathrm{D}_{14}} = f_{\mathrm{D}_{23}} = \frac{bK_\ell}{1+(1-b)K_\ell} f_{4\mathrm{M}}\,. \tag{33}$$

Together that yields for the fraction of DNA origami with all four caspases in monomeric form

$$f_{4\mathrm{M}} = \frac{1}{1+(4+2\frac{b}{1+(1-b)K_\ell})K_\ell+2K_\ell{}^2}\,, \tag{34}$$

for the fraction of DNA origami with one specific horizontal or vertical dimer

$$f_{\mathrm{D}_{12}} = f_{\mathrm{D}_{34}} = f_{\mathrm{D}_{13}} = f_{\mathrm{D}_{24}} = \frac{K_\ell}{1+(4+2\frac{b}{1+(1-b)K_\ell})K_\ell+2K_\ell{}^2}\,, \tag{35}$$

for the fraction of origamis with one specific diagonal dimer

$$f_{\mathrm{D}_{14}} = f_{\mathrm{D}_{23}} = \frac{\frac{b}{1+(1-b)K_\ell}K_\ell}{1+(4+2\frac{b}{1+(1-b)K_\ell})K_\ell+2K_\ell{}^2}\,, \tag{36}$$

and for the fraction of origamis in one specific state with two dimers

$$f_{2\mathrm{D_V}} = f_{2\mathrm{D_H}} = \frac{K_\ell{}^2}{1+\left(4+2\frac{b}{1+(1-b)K_\ell}\right)K_\ell+2K_\ell{}^2}\,. \tag{37}$$

In terms of the dimerization probability $f_\mathrm{D}$ this yields

$$f_{4\mathrm{M}} = \frac{1}{1+\left(4+2\frac{b}{1+(1-b)\frac{f_\mathrm{D}}{1-f_\mathrm{D}}}\right)\frac{f_\mathrm{D}}{1-f_\mathrm{D}}+2\left(\frac{f_\mathrm{D}}{1-f_\mathrm{D}}\right)^2} = \frac{(1-f_\mathrm{D})^2(1-bf_\mathrm{D})}{\left(1+2f_\mathrm{D}-f_\mathrm{D}{}^2\right)+bf_\mathrm{D}\left(1-6f_\mathrm{D}+3f_\mathrm{D}{}^2\right)} \tag{38}$$

$$f_{\mathrm{D}_{12}} = f_{\mathrm{D}_{34}} = f_{\mathrm{D}_{13}} = f_{\mathrm{D}_{24}} = \frac{\frac{f_\mathrm{D}}{1-f_\mathrm{D}}}{1+\left(4+2\frac{b}{1+(1-b)\frac{f_\mathrm{D}}{1-f_\mathrm{D}}}\right)\frac{f_\mathrm{D}}{1-f_\mathrm{D}}+2\left(\frac{f_\mathrm{D}}{1-f_\mathrm{D}}\right)^2}$$

$$= \frac{f_\mathrm{D}(1-f_\mathrm{D})(1-bf_\mathrm{D})}{\left(1+2f_\mathrm{D}-f_\mathrm{D}{}^2\right)+bf_\mathrm{D}\left(1-6f_\mathrm{D}+3f_\mathrm{D}{}^2\right)} \tag{39}$$

$$f_{\mathrm{D}_{14}} = f_{\mathrm{D}_{23}} = \frac{\frac{b}{1+(1-b)\frac{f_\mathrm{D}}{1-f_\mathrm{D}}}\left(\frac{f_\mathrm{D}}{1-f_\mathrm{D}}\right)}{1+\left(4+2\frac{b}{1+(1-b)\frac{f_\mathrm{D}}{1-f_D}}\right)\frac{f_\mathrm{D}}{1-f_\mathrm{D}}+2\left(\frac{f_\mathrm{D}}{1-f_\mathrm{D}}\right)^2} = \frac{b\,f_\mathrm{D}(1-f_\mathrm{D})^2}{\left(1+2f_\mathrm{D}-f_\mathrm{D}{}^2\right)+bf_\mathrm{D}\left(1-6f_\mathrm{D}+3f_\mathrm{D}{}^2\right)} \tag{40}$$

$$f_{2\mathrm{D_V}} = f_{2\mathrm{D_H}} = \frac{\left(\frac{f_\mathrm{D}}{1-f_\mathrm{D}}\right)^2}{1+\left(4+2\frac{b}{1+(1-b)\frac{f_\mathrm{D}}{1-f_\mathrm{D}}}\right)\frac{f_\mathrm{D}}{1-f_\mathrm{D}}+2\left(\frac{f_\mathrm{D}}{1-f_\mathrm{D}}\right)^2} = \frac{f_\mathrm{D}{}^2(1-bf_\mathrm{D})}{\left(1+2f_\mathrm{D}-f_\mathrm{D}{}^2\right)+bf_\mathrm{D}\left(1-6f_\mathrm{D}+3f_\mathrm{D}{}^2\right)} \tag{41}$$

**Comparison of the average number of dimers per DNA origami for Cases 1-5.** For the two-enzyme configurations the average number of dimers per DNA origami simply equals the fraction of DNA origamis with the caspases in dimeric form. For the proximal two-enzyme configuration (Case 1) the average number of dimers per DNA origami thus simply equals the dimerization probability $f_\mathrm{D}$, while for the more distal two-enzyme configuration (Case 2) the average number of dimers per DNA origami equals $af_\mathrm{D}$.

In Case 3, i.e. the DNA origami with 3 caspases in a linear geometry, the average number of dimers per DNA origami equals the sum of the probabilities either of the dimers is present, i.e. the sum of $f_{\mathrm{D}_{12}}$, $f_{\mathrm{D}_{23}}$, and $f_{\mathrm{D}_{13}}$. Using Eqs. (24) and (25), this yields for the average number of dimers per DNA origami

$$2\frac{f_\mathrm{D}(1-af_\mathrm{D})}{1+f_\mathrm{D}-2af_\mathrm{D}{}^2} + \frac{a(1-f_\mathrm{D})\,f_\mathrm{D}}{1+f_\mathrm{D}-2af_\mathrm{D}{}^2} = \frac{(2+a)f_\mathrm{D}-3af_\mathrm{D}{}^2}{1+f_\mathrm{D}-2af_\mathrm{D}{}^2}. \tag{42}$$

In Case 4, i.e. the DNA origami with 3 caspases in a triangular geometry, the average number of dimers per DNA origami equals the sum of the probabilities of the 3 possible dimer states according to Eq. (29), resulting in $\frac{3f_D}{1+2f_D}$ (i.e. the same as Eq. (42) assuming $a = 1$).

In Case 5, i.e. the origami with 4 caspases in a square geometry, the average number of dimers per DNA origami equals the sum of the probabilities of the single dimer states plus twice the sum of the probabilities of the double dimer states, i.e., $f_{\mathrm{D}_{12}} + f_{\mathrm{D}_{34}} + f_{\mathrm{D}_{13}} + f_{\mathrm{D}_{24}} + f_{\mathrm{D}_{14}} + f_{\mathrm{D}_{23}} + 2\left(f_{2\mathrm{D_V}} + f_{2\mathrm{D_H}}\right)$. Using Eqs. (39)-(41), that yields for the average number of dimers per DNA origami

$$\frac{4f_{\mathrm{D}}(1-f_{\mathrm{D}})(1-bf_{\mathrm{D}})+2b\,f_{\mathrm{D}}(1-f_{\mathrm{D}})^2+4f_{\mathrm{D}}{}^2(1-bf_{\mathrm{D}})}{(1+2f_{\mathrm{D}}-f_{\mathrm{D}}{}^2)+bf_{\mathrm{D}}(1-6f_{\mathrm{D}}+3f_{\mathrm{D}}{}^2)} = f_{\mathrm{D}}\frac{4+2b(1-4f_{\mathrm{D}}+f_{\mathrm{D}}{}^2)}{(1+2f_{\mathrm{D}}-f_{\mathrm{D}}{}^2)+bf_{\mathrm{D}}(1-6f_{\mathrm{D}}+3f_{\mathrm{D}}{}^2)}\,. \tag{43}$$

For $b=0$ this reduces to

$$\frac{4f_{\mathrm{D}}}{1+2f_{\mathrm{D}}-f_{\mathrm{D}}^2} \tag{44}$$

while for $b=1$ it reduces to

$$f_{\mathrm{D}}\frac{6-8f_{\mathrm{D}}+2f_{\mathrm{D}}{}^2}{1+3f_{\mathrm{D}}-7f_{\mathrm{D}}{}^2+3f_{\mathrm{D}}{}^3}\,. \tag{45}$$

In **Supplementary Figure 30**, these average numbers of dimers per DNA origami are summarized and plotted as a function of the dimerization probability $f_{\mathrm{D}}$ for all 5 cases.

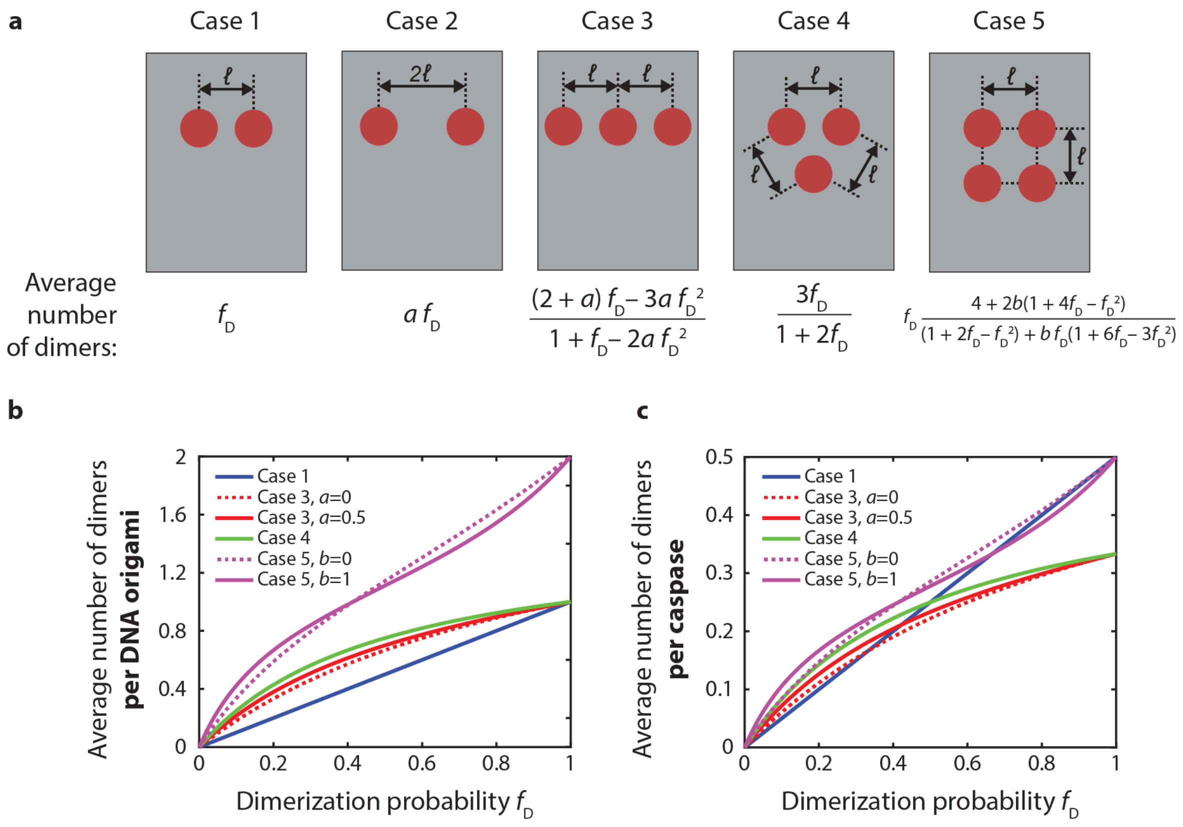


**Supplementary Figure 30 | The average number of dimers for all cases. a,** Overview of the formulae per case. **b,c,** Graphical representation of the average number of dimers per DNA origami (**b**) and per caspase (**c**), plotted as function of the dimerization probability $f_{\mathrm{D}}$.

#### Section 2: dimerization of tethered enzymes with incomplete enzyme incorporation

In the previous section we derived the average number of dimers per DNA origami for five different enzyme configurations as a function of the dimerization probability $f_{\mathrm{D}}$ in the proximal two-enzyme configuration. In this derivation we assumed that at each of the designated positions on the DNA origami a caspase is present. In practice, however, incorporation of caspases is not perfect, such that caspases will only be present at a certain fraction of the designated positions. In this section we will derive how this *incorporation efficiency* ($p$) influences the average number of dimers per DNA origami, for each of the 5 cases considered in the previous section. In this we will assume that this incorporation efficiency is equal for all designated positions and independent on the presence of neighboring caspases.

**Case 1 and 2: two-enzyme configurations.** For a two-enzyme configuration there are 4 possibilities. The first possibility, with probability $(1-p)^2$, is that both sites remain empty. The second possibility, with probability $p(1-p)$, is that a caspase is present only at the first site. The third possibility, i.e. that the second site contains a caspase, has the same probability. And the fourth and last possibility, with probability $p^2$, is that both sites are occupied. These 4 possibilities are also shown in **Supplementary Figure 31**, for the two-enzyme configuration at distance $\ell$, along with their probabilities to occur. In the first 3 possibilities of course no dimers can be formed, while the fourth possibility is exactly the situation for which we derived the average number of dimers to be equal to $f_\mathrm{D}$ in the previous section. Combined, with incorporation efficiency $p$ and dimerization efficiency $f_\mathrm{D}$, the average number of dimers per DNA origami equals

$$p^2\, f_\mathrm{D}. \tag{46}$$

For the two-enzyme configuration at $2\ell$, the probabilities of the 4 incorporation states are the same and the resulting average number of dimers per origami equals

$$p^2 a f_\mathrm{D}. \tag{47}$$

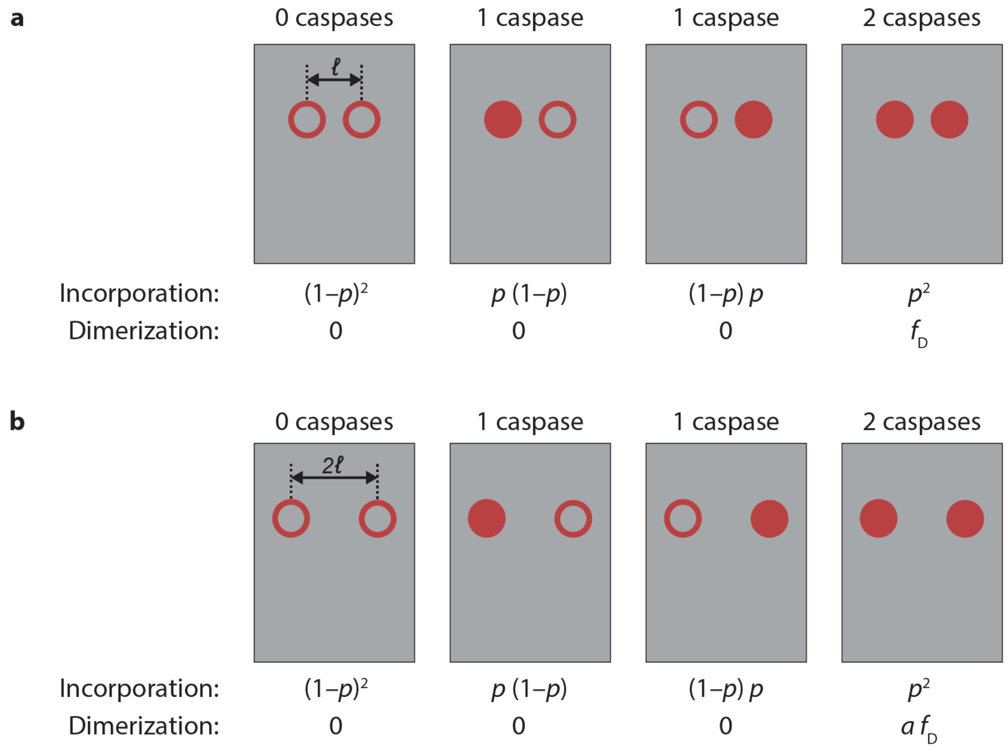


**Supplementary Figure 31 |** All possible occupancy states for two-enzyme configurations at distance $\ell$ (**a**) and $2\ell$ (**b**) with their corresponding probability at incorporation efficiency $p$ as well as the corresponding average number of dimers per DNA origami (as calculated for that geometry in Section 1).

**Case 3: linear three-enzyme configuration.** With three sites for enzyme incorporation, each with efficiency $p$, the number of possibilities increases to 8. The first possibility, with probability $(1-p)^3$, is that all three sites remain empty. Then there are three possibilities, each with probability $p(1-p)^2$, that one of the sites is occupied and the other two remain empty. In all of these first four possibilities no dimers can be formed, as they contain at most 1 caspase. Then there are also three possibilities, each with probability $p^2(1-p)$, that one site remains empty while the other two are occupied. The two cases where one of these two sites is in the middle are identical to the situation for which we derived (Case 1 in Section 1) the average number of dimers per DNA origami to be equal to $f_{\mathrm{D}}$. The possibility in which only both outer sites are occupied is identical to the situation of Case 2 in Section 1 where the average number of dimers per DNA origami equals $af_{\mathrm{D}}$. The final possibility, with probability $p^3$, is that all three sites are occupied. This is again exactly the situation for which we derived (Case 3 in Section 1) the average number of dimers per DNA origami to be equal to $\frac{(2+a)f_{\mathrm{D}}-3af_{\mathrm{D}}{}^2}{1+f_{\mathrm{D}}-2af_{\mathrm{D}}{}^2}$. These 8 possibilities are shown in **Supplementary Figure 32** along with their probabilities to occur and the average number of dimers per DNA origami. Combined that thus means that, with incorporation efficiency $p$ and dimerization probability $f_{\mathrm{D}}$, the average number of dimers per DNA origami equals

$$p^2(1-p)f_{\mathrm{D}}(2+a)+p^3\frac{(2+a)f_{\mathrm{D}}-3af_{\mathrm{D}}{}^2}{1+f_{\mathrm{D}}-2af_{\mathrm{D}}{}^2}. \quad (48)$$

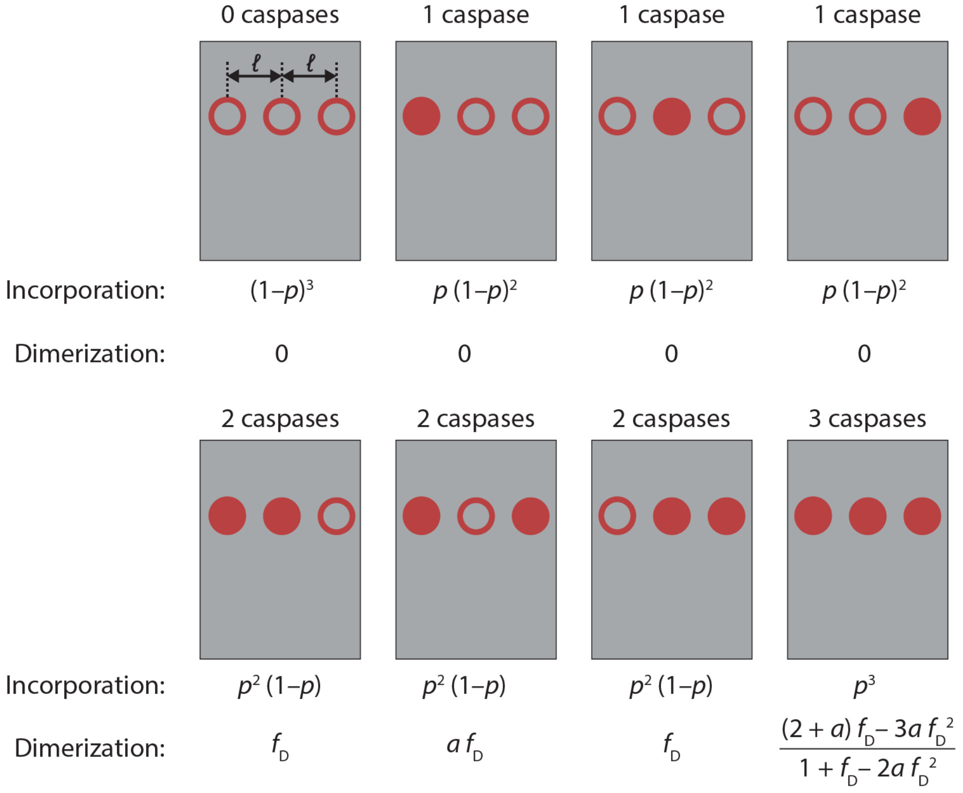


**Supplementary Figure 32 |** All possible occupancy states for the linear three-enzyme configuration with their corresponding probability at incorporation efficiency $p$ as well as the corresponding average number of dimers per DNA origami (as calculated for that geometry in Section 1).

**Case 4: triangular three-enzyme configuration.** With three linkers in a triangular geometry, the number of possibilities for enzyme occupancy remains the same as in Case 3, i.e. 8, and also the probabilities for all instances remain unchanged. These 8 possibilities are shown in **Supplementary Figure 33**. In the 4 possibilities with 0 or 1 enzymes still no dimers can be formed. In the 3 possibilities with 2 caspases, however, it no longer matters which of the three sites remains empty. In each of these possibilities there are simply 2 neighboring enzymes, which follows exactly the situation for which we derived (Case 1 in Section 1) the average number of dimers per DNA origami to be equal to $f_D$. The final possibility, i.e. where all three linkers bind a caspase, is exactly the situation for which we derived (Case 4 in Section 1) the average number of dimers per DNA origami as $3f_{\mathrm{D}}/(1+2f_{\mathrm{D}})$. Combined, with incorporation efficiency $p$ and dimerization probability $f_{\mathrm{D}}$, the average number of dimers per DNA origami thus equals

$$3p^2(1-p)f_\mathrm{D} + 3p^3\frac{f_\mathrm{D}}{(1+2f_\mathrm{D})} = \frac{3p^2 f_\mathrm{D}(1+2f_\mathrm{D}(1-p))}{(1+2f_\mathrm{D})} \quad (49)$$

An important control for this configuration is the '2+1' configuration (reflecting experimental configurations [**126**] and [**256**]), in which one of the incorporation sites is positioned such that dimerization is not possible. This case is illustrated in **Supplementary Figure 34**, with again for each occupancy state its probability to occur and the corresponding average number of dimers per DNA origami indicated. Only in the 2 states where both sites that are close to each other are occupied, a dimer can be formed. Combined, with incorporation efficiency $p$ and dimerization probability $f_\mathrm{D}$, the average number of dimers per origami thus equals $p^2(1-p)f_\mathrm{D} + p^3 f_\mathrm{D}$, which simplifies to $p^2 f_\mathrm{D}$. This is, not surprisingly, exactly the same number as obtained in the earlier Case 1.

The average number of dimers per DNA origami for both the triangular and '2+1' three-enzyme configurations are shown as a function of the incorporation efficiency $p$ and the dimerization probability $f_\mathrm{D}$ in **Supplementary Figure 35**. The relative difference in the average number of dimers per DNA origami between these two configurations, i.e., the difference between the value for the triangular configuration minus that for the '2+1' configuration divided by the value for the '2+1' configuration, is also shown. For $f_\mathrm{D} = 0.9, p = 0.75$ this yields 0.786 as average number of dimers per DNA origami for the triangular configuration and 0.506 as average number of dimers per DNA origami on the '2+1' configuration. This corresponds to an increase of 0.280 dimers per DNA origami, i.e., a 55% increase.

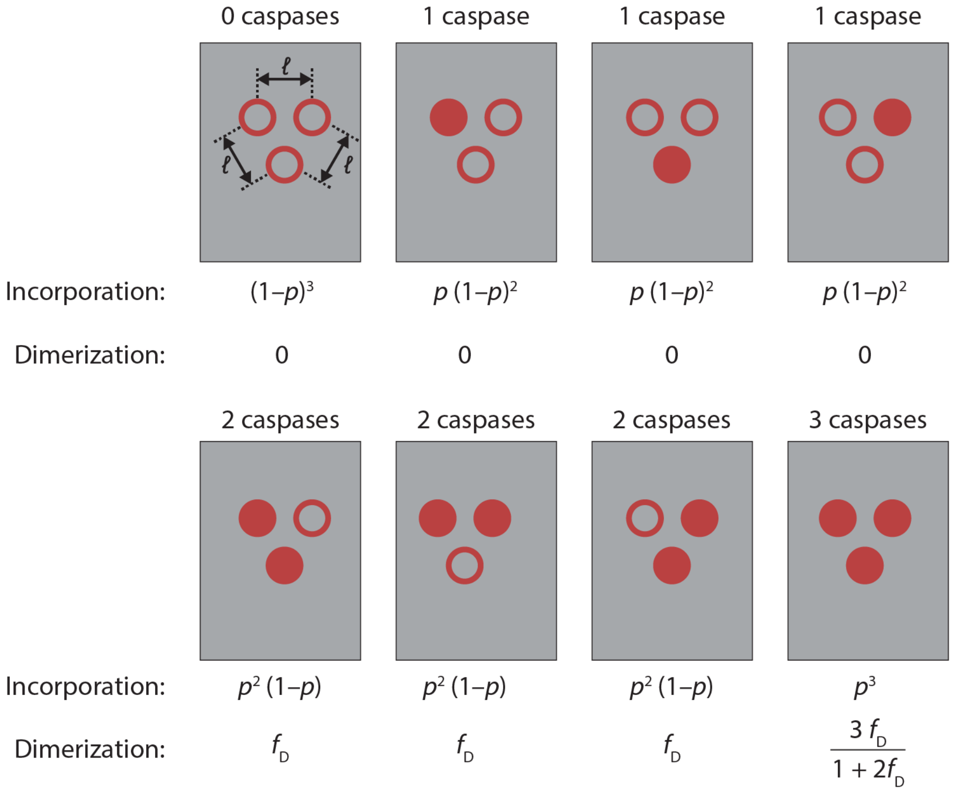


**Supplementary Figure 33 |** All possible occupancy states for the triangular three-enzyme configuration with their corresponding probability at incorporation efficiency $p$as well as the corresponding average number of dimers per DNA origami (as calculated for that geometry in Section 1).

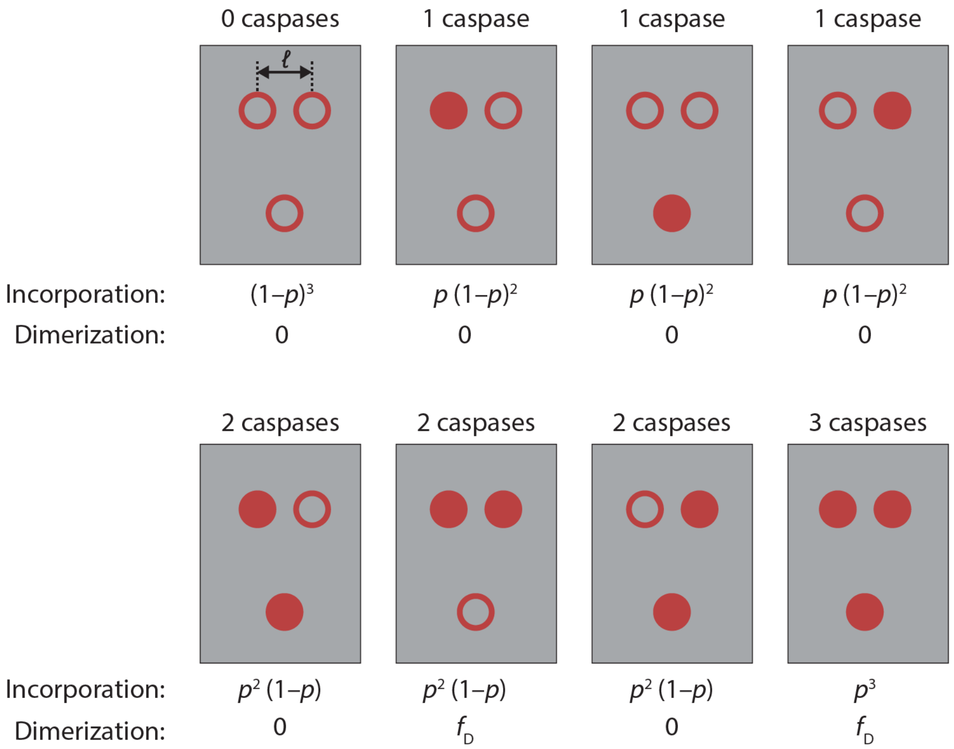


**Supplementary Figure 34 |** All possible occupancy states for the '2+1' three-enzyme configuration with their corresponding probability at incorporation efficiency $p$ as well as the corresponding average number of dimers per DNA origami (as calculated for that geometry in Section 1).

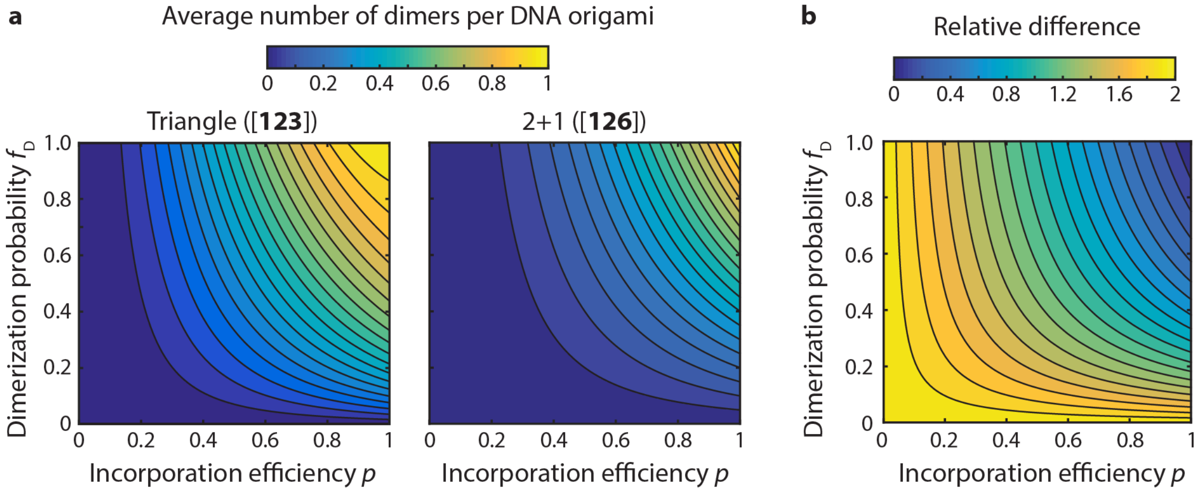


**Supplementary Figure 35 |** The average number of dimers per DNA origami as a function of both incorporation efficiency $p$ and the dimerization probability $f_D$ for the triangular three-enzyme configuration (**a**; left graph, reflecting experimental configuration [**123**]) and control '2+1' configuration (**a**; right graph, reflecting experimental configurations [**126**] and [**256**]). **b,** Relative increase in the average number of dimers per DNA origami in triangular configuration as compared to the control '2+1' configuration.

**Case 5: four-enzyme configurations.** With four enzyme incorporation sites in a proximal configuration (reflecting the experimental [**1234**] configuration), each with probability $p$, the number of possibilities increases to 16. The first possibility, with probability $(1-p)^4$, is that all sites remain empty. Then there are four possibilities, each with probability $p(1-p)^3$, that one site is occupied while the other three sites remain empty. In all of these first 5 possibilities no dimers can be formed. Next, there are six possibilities, each with probability $p^2(1-p)^2$, that two sites remain empty while the other two are occupied. In two out of these six instances two linkers on opposite corners are bound, in which case the average number of dimers per DNA origami is $bf_{\mathrm{D}}$ (Case 2 in Section 1 with a factor $b$ instead of $a$), while the other 4 instances are equivalent to the situation for which we derived (Case 1 in Section 1) the average number of dimers per DNA origami as $f_{\mathrm{D}}$. Then there are four possibilities, each with probability $p^3(1-p)$, that one site remains empty while the other three are occupied. Though the geometry is not exactly the same as in Case 3 in Section 1, the average number of dimers is given by the same formula but with factor $b$ instead of $a$, i.e., $\frac{(2+b)f_{\mathrm{D}}-3bf_{\mathrm{D}}{}^2}{1+f_{\mathrm{D}}-2bf_{\mathrm{D}}{}^2}$. Finally, the last possibility, with probability $p^4$, is that all four sites are occupied. This is again exactly the situation for which we derived (Case 5 in Section 1) the average number of dimers per DNA origami to be equal to $f_D\frac{4+2b(1-4f_{\mathrm{D}}+f_{\mathrm{D}}{}^2)}{(1+2f_{\mathrm{D}}-f_{\mathrm{D}}{}^2)+bf_{\mathrm{D}}(1-6f_{\mathrm{D}}+3f_{\mathrm{D}}{}^2)}$. All these 16 possible incorporation states are shown in **Supplementary Figure 36** along with their probability to occur and the average number of dimers per DNA origami. Combined that thus means that, with incorporation efficiency $p$ and dimerization probability $f_D$, the average number of dimers per origami equals

$$p^2(1-p)^2(4+2b)f_{\mathrm{D}}+p^3(1-p)\,4\frac{(2+b)f_{\mathrm{D}}-3bf_{\mathrm{D}}{}^2}{1+f_{\mathrm{D}}-2bf_{\mathrm{D}}{}^2}+p^4f_D\frac{4+2b(1-4f_{\mathrm{D}}+f_{\mathrm{D}}{}^2)}{(1+2f_{\mathrm{D}}-f_{\mathrm{D}}{}^2)+bf_{\mathrm{D}}(1-6f_{\mathrm{D}}+3f_{\mathrm{D}}{}^2)}. \qquad (50)$$

An important control for the proximal four-enzyme configuration is the distal setup, in which two pairs of incorporation sites are positioned such that the distance between the pairs is too large to allow for dimerization in that direction. This case is illustrated in **Supplementary Figure 37**, with again for each occupancy state its probability to occur and the corresponding average number of dimers per DNA origami indicated. Only in the states where two neighboring sites are occupied, a dimer can be formed. Combined that thus means that, with incorporation efficiency $p$ and dimerization probability $f_{\mathrm{D}}$, the average number of dimers per DNA origami equals $2p^2(1-p)^2f_{\mathrm{D}}+4p^3(1-p)f_{\mathrm{D}}+2p^4f_{\mathrm{D}}$, which simplifies to $2p^2f_{\mathrm{D}}$. Since this configuration can be viewed as two non-interacting two-enzyme configurations, but on the same DNA origami, we obtained not surprisingly the same result (multiplied by two) as obtained in the earlier Case 1 for the single two-enzyme configuration.

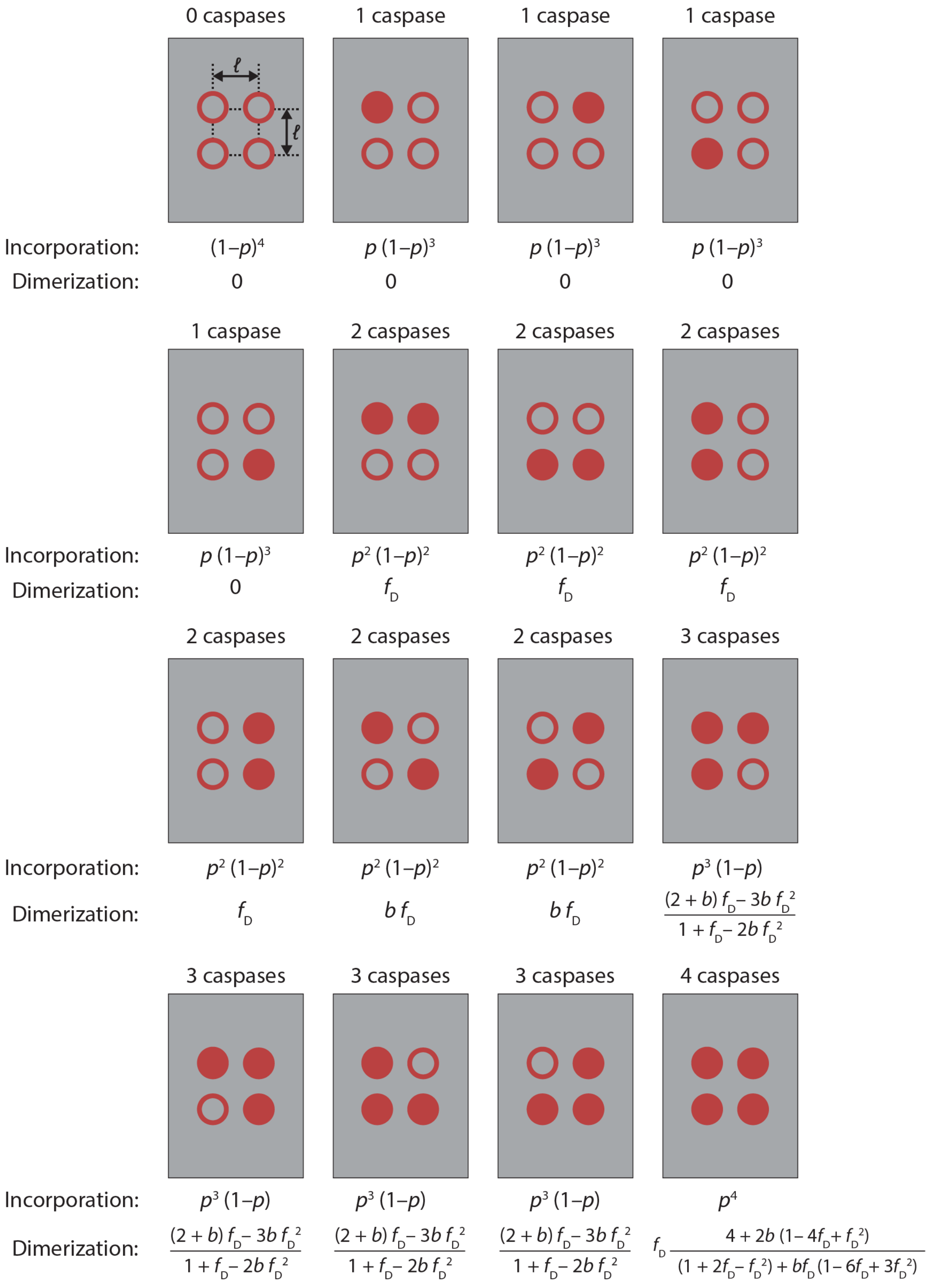


**Supplementary Figure 36 |** All possible occupancy states for the proximal four-enzyme configuration with their corresponding probability at incorporation efficiency $p$ as well as the corresponding average number of dimers per DNA origami (as calculated for that geometry in Section 1).

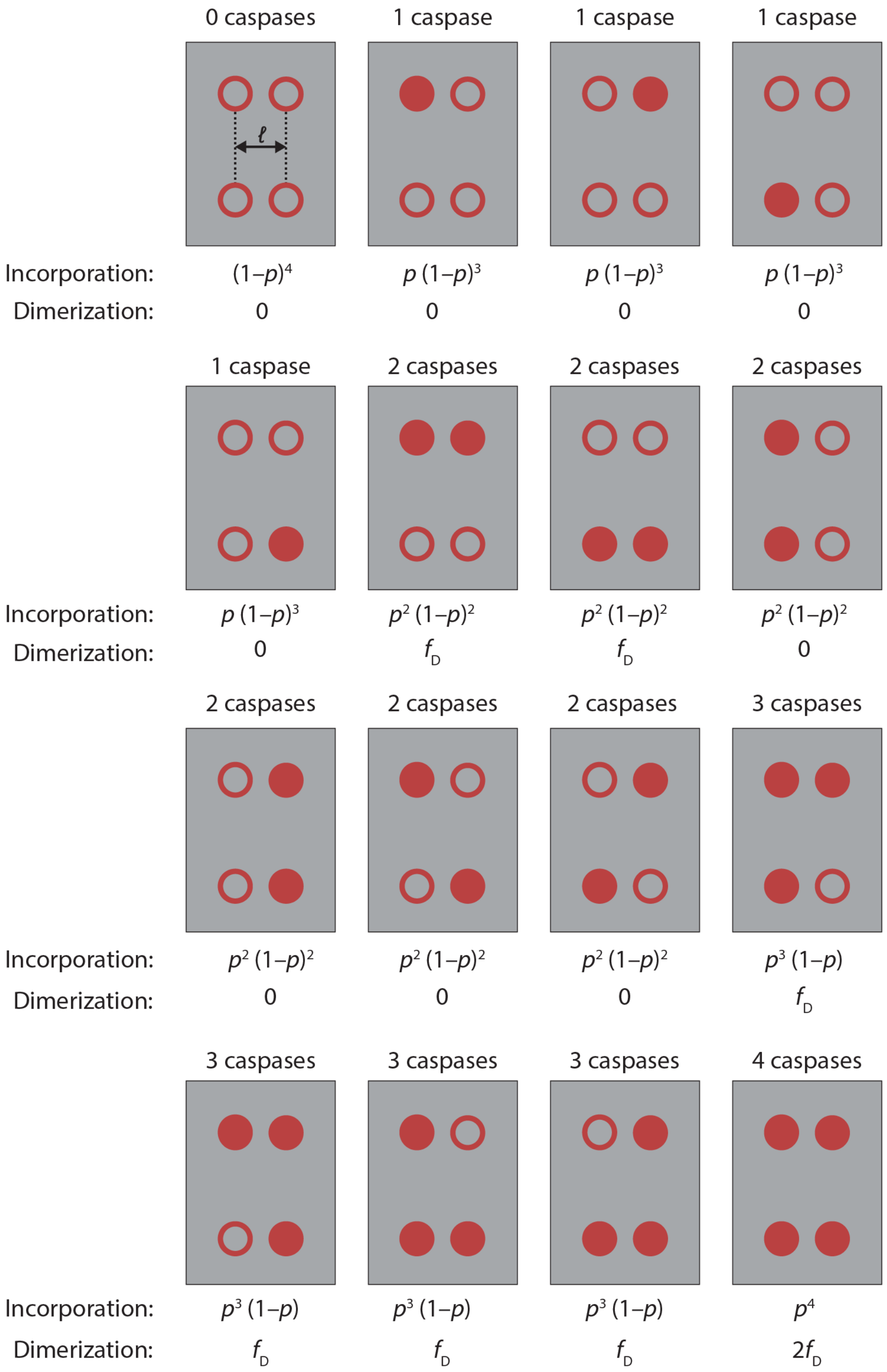


**Supplementary Figure 37 |** All possible occupancy states for the distal four-enzyme configuration with their corresponding probability at incorporation efficiency $p$ as well as the corresponding average number of dimers per DNA origami (as calculated for that geometry in Section 1).

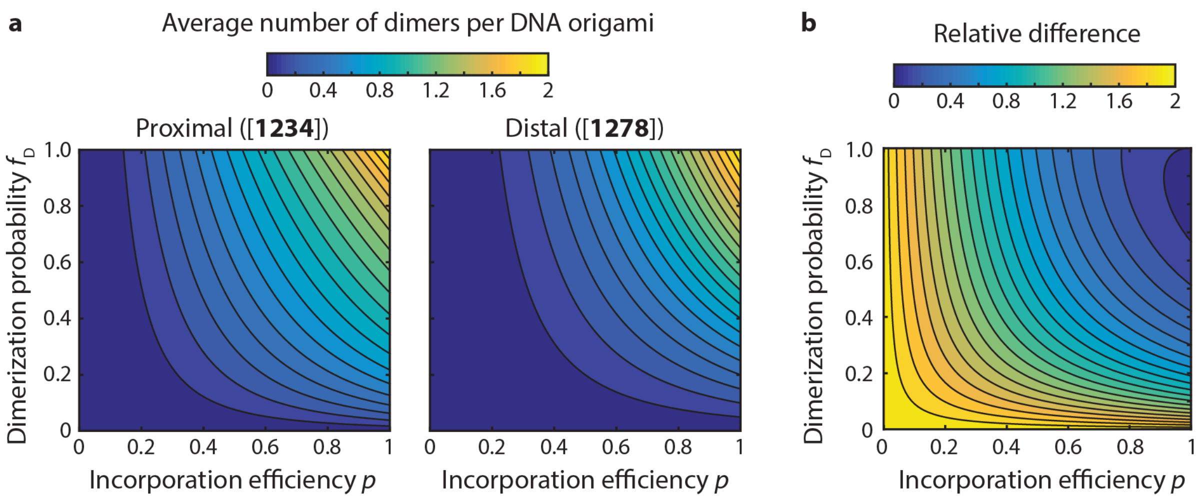


**Supplementary Figure 38 |** The average number of dimers per DNA origami as a function of the incorporation efficiency $p$ and dimerization probability $f_{\mathrm{D}}$ for the proximal (**a**; left graph, reflecting experimental configuration [**1234**]) and distal four-enzyme configuration (**a**; right graph, reflecting experimental configuration [**1278**]). **b,** Relative increase in the average number of dimers per DNA origami in the proximal configuration as compared to the control distal configuration.

The average number of dimers per DNA origami for both the proximal and distal four-enzyme configurations are shown as a function of the incorporation efficiency $p$ and the dimerization probability $f_{\mathrm{D}}$ in **Supplementary Figure 38**. The relative difference in the average number of dimers per DNA origami between these two configurations, i.e., the difference between the value for the proximal minus the distal configuration divided by the value for the distal configuration, is also shown. With full diagonal interactions, i.e. $b = 1$, and with $f_{\mathrm{D}} = 0.9, p = 0.75$, this yields 1.148 as the average number of dimers per DNA origami in the proximal configuration and 1.013 as the average number of dimers per DNA origami in the distal configuration. That comprises an increase of 0.135 dimers per DNA origami, which corresponds to a 13.4% increase.

The results shown in the histograms of **Figure 4f** were directly obtained by evaluating Eqs. (46)-(50) derived in this section, assuming $f_{\mathrm{D}} = 0.9, p = 0.75, a = 0.5, b = 1$.

### DNA origami design and DNA sequences

The DNA origami rectangle used in this study was designed using caDNAno v0.2 based on the *tall rectangle* design by Rothemund (**Supplementary Figure 39** and **Supplementary Figure 40**)[8]. The 7249-nt single-stranded M13mp18 scaffold strand folds into a single-layer structure of 32 helices using 192 staple strands (**Supplementary Table 6**). To correct for global twist of the DNA origami rectangle, 3 base pair deletions per helix were introduced[21]. To prevent DNA origami aggregation through blunt-end stacking all 32 edge staples were omitted during folding and not listed here. For incorporation of caspase-9 enzyme-DNA conjugates, various unmodified staples were replaced by staples functionalized at the 3' end with 15-nt handle strands (**Supplementary Table 3**-**Supplementary Table 5**). A short single-stranded TT linker was included between staple and handle to prevent biased orientation of the handle with respect to the DNA origami structure[22,23].

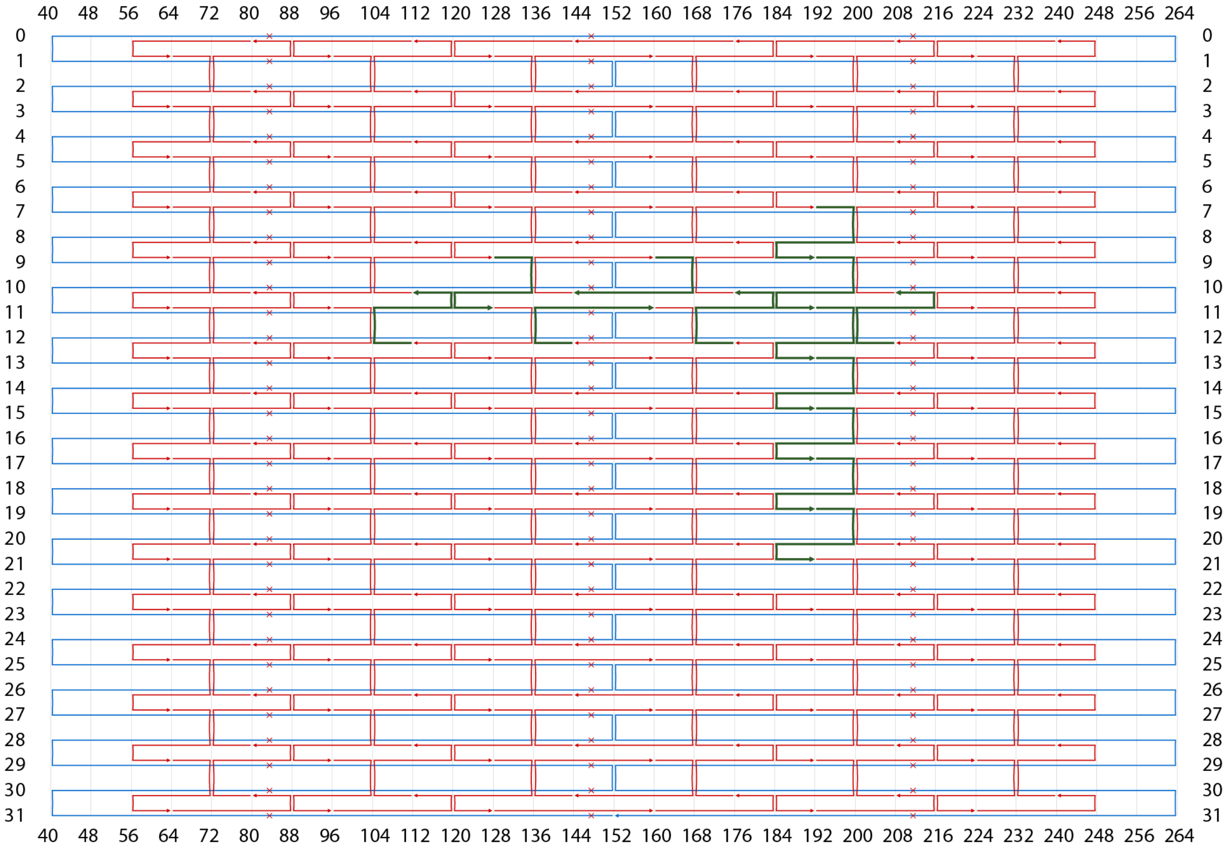


**Supplementary Figure 39 | Schematic overview of the DNA origami structure used for all two-enzyme DNA nanostructures.** The scaffold strand is shown in light blue and unmodified staple strands in red. Staples that are used for enzyme incorporation are shown in green and correspond to handle-extended staple strands in **Supplementary Table 3** and **Supplementary Table 5**. Base-pair deletions to correct for global twist of the structure are indicated by crosses and 3' ends of DNA strands are indicated by arrows. Numbers on left and right indicate the reference helix number, while numbers on top and bottom indicate reference nucleotide position.

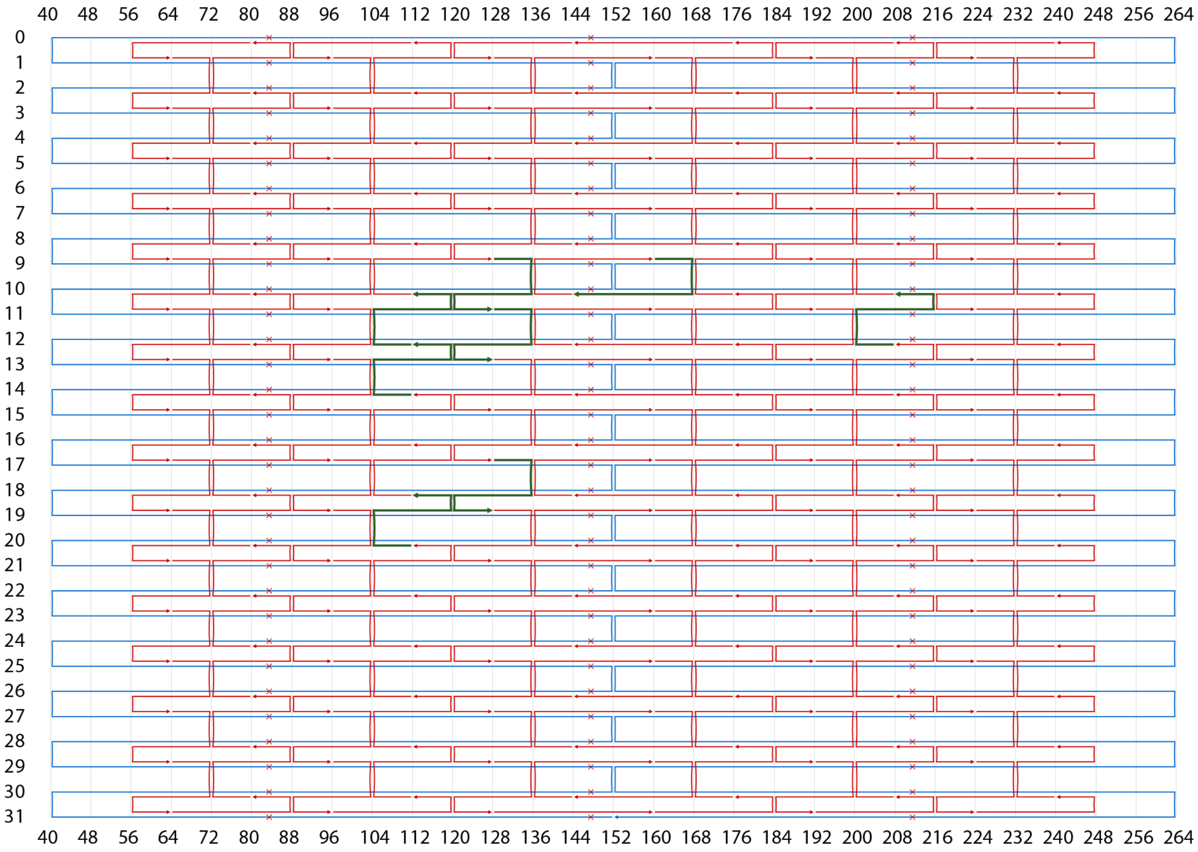


**Supplementary Figure 40 | Schematic overview of the DNA origami structure used for three- and four-enzyme DNA nanostructures.** The scaffold strand is shown in light blue and unmodified staple strands in red. Staples that are used for enzyme incorporation are shown in green and correspond to handle-extended staple strands in **Supplementary Table 4**. Base-pair deletions to correct for global twist of the structure are indicated by crosses and 3’ ends of DNA strands are indicated by arrows. Numbers on left and right indicate the reference helix number, while numbers on top and bottom indicate reference nucleotide position.

**Supplementary Table 3 | Handle-extended staple strands for two-enzyme DNA nanostructures.** To assemble DNA origami structures for two-enzyme incorporation, two unmodified staple strands were replaced with handle-extended staple strands. For example, h1 and h3 were used for the 11-nm setup parallel to the helical axis, and v1 and i were used for the 6-nm setup perpendicular to the axis. Underlined thymine nucleotides were added as a spacer.

| ID | Unmodified staple ID | Distance to h1 (nm) | Distance to v1 (nm) | Sequence (5' to 3') |
|---|---|---|---|---|
| h1 | 74 | 0 | - | TCATTGAATTTTGCAAAAGAAGTTGATTCATCTTTCATACGACTCACTC |
| h2 | 57 | 6 | - | AAAAATCTTTATTACAGGTAGAAATTGCCAGATTTCATACGACTCACTC |
| h3 | 58 | 11 | - | TCTTACCGGAAACAATGAAATAGCACTAACGTTTCATACGACTCACTC |
| h4 | 75 | 17 | - | AGCGTCCATAGTAAAATGTTTAGTTTATCCCTTTCATACGACTCACTC |
| h5 | 76 | 22 | - | GTTACAAATAAGAAACGATTTTTTCCAATAATTTTCATACGACTCACTC |
| h7 | 77 | 33 | - | TAACGAGCGAAAATAGCAGCCTTAGAGAGATTTCATACGACTCACTC |
| v1 | 47 | - | 0 | AGACAAAAGCAACATATAAAAGAAAAGTAAGCTTTCATACGACTCACTC |
| v3 | 71 | - | 12 | TCAAAAATGTCTTTCCAGAGCCTAAGGCTTATTTTCATACGACTCACTC |
| v4 | 83 | - | 18 | CCGGTATTTCATCGAGAACAAGCAACGCGCCTTTTCATACGACTCACTC |
| v5 | 95 | - | 24 | GTTTATCAAGAGAATATAAAGTACAAAAGCCTTTTCATACGACTCACTC |
| v6 | 107 | - | 30 | GTTTAGTAAAATTTAATGGTTTGAGGTCTGAGTTTCATACGACTCACTC |
| v7 | 119 | - | 36 | AGACTACCCCCTTAGAATCCTTGAGATGAAACTTTCATACGACTCACTC |
| i | 59 | 28 | 6 | AGATAGCCAAGAATTGAGTTAAGCGTTTAACGTTTCATACGACTCACTC |

**Supplementary Table 4 | Handle-extended staple strands for three- and four-enzyme DNA nanostructures.** To assemble DNA origami structure for three- and four-enzyme incorporation, appropriate unmodified staple strands were replaced with handle-extended staple strands. Handle-staple IDs correspond to the notation used in **Figure 4**. Underlined thymine nucleotides were added as a spacer.

| ID | Unmodified staple ID | Sequence (5' to 3') |
|---|---|---|
| o1 | 74 | TCATTGAATTTTGCAAAAGAAGTTGATTCATCTTTCATACGACTCACTC |
| o2 | 57 | AAAAATCTTTATTACAGGTAGAAATTGCCAGATTTCATACGACTCACTC |
| o3 | 86 | CTTTAATTAAAGACTTCAAATATCATAAATATTTTCATACGACTCACTC |
| o4 | 69 | GGGGGTAAATACTGCGGAATCGTCGCGTTTTATTTCATACGACTCACTC |
| o5 | 58 | TCTTACCGGAAACAATGAAATAGCACTAACGTTTCATACGACTCACTC |
| o6 | 77 | TAACGAGCGAAAATAGCAGCCTTAGAGAGATTTCATACGACTCACTC |
| o7 | 122 | TAAAACTAGTAGCTATTTTTGAGATTTAGAACTTTCATACGACTCACTC |
| o8 | 105 | ACCCTGTACGCAAGGATAAAAATTGATCTACATTTCATACGACTCACTC |

**Supplementary Table 5 | Handle-extended staple strands for DNA nanostructures with inactive mutants.** To assemble DNA origami structure for homo- and heterodimers of inactive mutants, appropriate unmodified staple strands were replaced with handle-extended staple strands. For the inactive homodimers of **Figure 3e** m1 and m2 were used, while for the heterodimers of **Figure 5** either m1 and h2, or h1 and m2, were used. Underlined thymine nucleotides were added as a spacer.

| ID | Unmodified staple ID | Sequence (5' to 3') |
|---|---|---|
| m1 | 74 | TCATTGAATTTTGCAAAAGAAGTTGATTCATCTTACTGACTGACTGACT |
| m2 | 57 | AAAAATCTTTATTACAGGTAGAAATTGCCAGATTACTGACTGACTGACT |

**Supplementary Table 6 | Sequences of unmodified staple strands.** Locations of the 5’ and 3’ end are indicated using the reference helix number used in **Supplementary Figure 39** and **Supplementary Figure 40**, with the reference nucleotide position denoted in brackets. Staple strands indicated in green can be extended with handle strands to allow for enzyme incorporation (see **Supplementary Table 3**-**Supplementary Table 5**).

| Staple ID | Location of 5' end | Location of 3' end | Sequence (5’ to 3’) |
|---|---|---|---|
| 1 | 0[79] | 1[63] | ACTGAGTTTCGTCACCAGTACAAATCATAGTT |
| 2 | 0[111] | 1[95] | GCAAGCCCAATAGGAACCCATGTACGTCTTTC |
| 3 | 0[143] | 1[127] | CCTCAGAGCCACCACCCTCATTTTGTATGGGA |
| 4 | 0[175] | 0[144] | CCCTCAGAACCGCCACCCTCAGAACCGCCAC |
| 5 | 0[207] | 1[191] | TATCACCGTACTCAGGAGGTTTAGATTATTCT |
| 6 | 0[239] | 1[223] | GGGTTGATATAAGTATAGCCCGGATGAGACTC |
| 7 | 1[64] | 3[63] | AGCGTAACAAAAGGCTCCAAAAGGTTCGAGGT |
| 8 | 1[96] | 3[95] | CAGACGTTAATAATTTTTTCACGTCGATAGTT |
| 9 | 1[128] | 3[127] | TTTTGCTAAATAGAAAGGAACAACGCCCACGC |
| 10 | 1[160] | 2[144] | TGCCCCCTAACAGTGCCCGTATAATTTCAGC |
| 11 | 1[192] | 3[191] | GAAACATGTAATAAGTTTTAACGGAGGTTGAG |
| 12 | 1[224] | 3[223] | CTCAAGAGCATGGCTTTTGATGATTATTCACA |
| 13 | 2[79] | 0[80] | CTCCAAAAGATCTAAAGTTTTGTCCGTAAC |
| 14 | 2[111] | 0[112] | TTGCGAATAGTAAATGAATTTTCTCAGGGATA |
| 15 | 2[143] | 1[159] | GGAGTGAGAACAACTTTCAACAGACAGTTAA |
| 16 | 2[175] | 0[176] | CCTTGAGTGCCTATTTCGGAACCTTACCGCCA |
| 17 | 2[207] | 0[208] | TGTACTGGAAAGTATTAAGAGGCATAGGTG |
| 18 | 2[239] | 0[240] | GCGTCATAAAGGATTAGGATTAGCCGTCGAGA |
| 19 | 3[64] | 5[63] | GAATTTCTCAACGGCTACAGAGGCTTCCATTA |
| 20 | 3[96] | 5[95] | GCGCCGACGCAGCGAAAGACAGCACTACGAAG |
| 21 | 3[128] | 5[127] | ATAACCGATAAAGGCCGCTTTTGCAAAAGAAT |
| 22 | 3[160] | 4[144] | AGAGCCGCAGAGCCGCCACCAGAAAGGCTTG |
| 23 | 3[192] | 5[191] | GCAGGTCATCAGAACCGCCACCCTTTTGCCTT |
| 24 | 3[224] | 5[223] | AACAAATAACCGGAACCGCCTCCCTCATCGGC |
| 25 | 4[79] | 2[80] | GAGGGTAGTAAACAGCTTGATACTGAAAAT |
| 26 | 4[111] | 2[112] | CACCCTCAAATGACAACAACCATCTAAAGGAA |
| 27 | 4[143] | 3[159] | CAGGGAGTTATATTCGGTCGCTGCCACCACC |
| 28 | 4[175] | 2[176] | CCACCCTCCGCCAGCATTGACAGGGGTCAGTG |
| 29 | 4[207] | 2[208] | CGCCACCCGACGATTGGCCTTGAACAGGAG |
| 30 | 4[239] | 2[240] | GAGCCACCAATCCTCATTAAAGCCTCCAGTAA |
| 31 | 5[64] | 7[63] | AACGGGTACGACCTGCTCCATGTTATAAGGGA |
| 32 | 5[96] | 7[95] | GCACCAACATCATCGCCTGATAAAAGGACAGA |
| 33 | 5[128] | 7[127] | ACACTAAACAAGCGCGAAACAAAGTAGGCTGG |
| 34 | 5[160] | 6[144] | GTAATCAGAATGAAACCATCGATACCCCAGC |
| 35 | 5[192] | 7[191] | TAGCGTCACCATTACCATTAGCAAAGCGCCAA |
| 36 | 5[224] | 7[223] | ATTTTCGGGGAATTAGAGCCAGCAGATTGAGG |
| 37 | 6[79] | 4[80] | GAAATCCGAAATACGTAATGCCATCGGAAC |
| 38 | 6[111] | 4[112] | AGATTTGTCTAAAACGAAAGAGGCGGGATCGT |
| 39 | 6[143] | 5[159] | GATTATACACACTCATCTTTGACGCAGCACC |
| 40 | 6[175] | 4[176] | ACGTCACCTAGCGACAGAATCAAGCAGAGCCA |
| 41 | 6[207] | 4[208] | CAGTAGCAGACTGTAGCGCGTTTTCAGAGC |
| 42 | 6[239] | 4[240] | GCCATTTGTCATAGCCCCCTTATTCGGAACCA |
| 43 | 7[64] | 9[63] | ACCGAACTAAACACCAGAACGAGTCTTTAATC |
| 44 | 7[96] | 9[95] | TGAACGGTTCATTCAGTGAATAAGTTAAGAAC |
| 45 | 7[128] | 9[127] | CTGACCTTATTCATTACCCAAATCTTGGGAAG |
| 46 | 7[160] | 8[144] | AATAGAAAGAATAAGTTTATTTTGTGACAAG |
| **47** | **7[192]** | **9[191]** | **AGACAAAAGCAACATATAAAAGAAAAGTAAGC** |
| 48 | 7[224] | 9[223] | GAGGGAAGAGCAAACGTAGAAAATAAGGAAAC |
| 49 | 8[79] | 6[80] | CTGACGAGGACCAACTTTGAAAGTTGTGTC |
| 50 | 8[111] | 6[112] | AAAGCTGCGTACAGACCAGGCGCATACAACGG |

| | | | |
|---|---|---|---|
| 51 | 8[143] | 7[159] | AACCGGATCATCAAGAGTAATCTTCACAATC |
| 52 | 8[175] | 6[176] | ACACCACGATTCATATGGTTTACCGGCCGGAA |
| 53 | 8[207] | 6[208] | TAAAGGTGGGGCGACATTCAACCAAATCAC |
| 54 | 8[239] | 6[240] | AGTATGTTGTAAATATTGACGGAACGACTTGA |
| 55 | 9[64] | 11[63] | ATTGTGAAACATAACGCCAAAAGGTAACCCTC |
| 56 | 9[96] | 11[95] | TGGCTCATTTTAGGAATACCACATCAAAATAG |
| **57** | **9[128]** | **11[127]** | **AAAAATCTTTATTACAGGTAGAAATTGCCAGA** |
| **58** | **9[160]** | **10[144]** | **TCTTACCGGAAACAATGAAATAGCACTAACG** |
| **59** | **9[192]** | **11[191]** | **AGATAGCCAAGAATTGAGTTAAGCGTTTAACG** |
| 60 | 9[224] | 11[223] | CGAGGAAAATTGAGCGCTAATATCTACAGAGA |
| 61 | 10[79] | 8[80] | ATGCAGATTTACCTTATGCGATTGCTTGCC |
| 62 | 10[111] | 8[112] | AGTTGAGATATACCAGTCAGGACGAACGTAAC |
| 63 | 10[143] | 9[159] | GAACAACAACGTTAATAAAACGAAATAGCTA |
| 64 | 10[175] | 8[176] | AAGAGCAAAAGCCCTTTTTAAGAAACGCAAAG |
| 65 | 10[207] | 8[208] | TAACCCACGAACAAAGTTACCAGACATACA |
| 66 | 10[239] | 8[240] | AGAGGGTACGCAATAATAACGGAATATTACGC |
| 67 | 11[64] | 13[63] | GTTTACCAACGAGAATGACCATAAAGTCAGAA |
| 68 | 11[96] | 13[95] | CGAGAGGCTCCCCCTCAAATGCTTATTAAGAG |
| **69** | **11[128]** | **13[127]** | **GGGGGTAAATACTGCGGAATCGTCGCGTTTTA** |
| 70 | 11[160] | 12[144] | AATCCAAAATAAACAGCCATATTAACTGGAT |
| **71** | **11[192]** | **13[191]** | **TCAAAAATGTCTTTCCAGAGCCTAAGGCTTAT** |
| 72 | 11[224] | 13[223] | GAATAACATTTATCCTGAATCTTAGTTTTAGC |
| 73 | 12[79] | 10[80] | TTCAGAAAGACGACGATAAAAACTCAACTA |
| **74** | **12[111]** | **10[112]** | **TCATTGAATTTTGCAAAAGAAGTTGATTCATC** |
| **75** | **12[143]** | **11[159]** | **AGCGTCCATAGTAAAATGTTTAGTTTATCCC** |
| **76** | **12[175]** | **10[176]** | **GTTACAAATAAGAAACGATTTTTCCAATAAT** |
| **77** | **12[207]** | **10[208]** | **TAACGAGCGAAAATAGCAGCCTTAGAGAGA** |
| 78 | 12[239] | 10[240] | GCTACAATTAAAAACAGGGAAGCGACAAAGTC |
| 79 | 13[64] | 15[63] | GCAAAGCGTGGCTTAGAGCTTAATTAAATATG |
| 80 | 13[96] | 15[95] | GAAGCCCGGCTCCTTTTGATAAGAGTTTCATT |
| 81 | 13[128] | 15[127] | ATTCGAGCCAACAGGTCAGGATTACTGCGAAC |
| 82 | 13[160] | 14[144] | AATAGCAAATCGTAGGAATCATTAACCGGAA |
| **83** | **13[192]** | **15[191]** | **CCGGTATTTCATCGAGAACAAGCAACGCGCCT** |
| 84 | 13[224] | 15[223] | GAACCTCCCCAAGAACGGGTATTATGAACAAG |
| 85 | 14[79] | 12[80] | TTTGCGGAGATTGCATCAAAAAGTAAACAG |
| **86** | **14[111]** | **12[112]** | **CTTTAATTAAAGACTTCAAATATCATAAATAT** |
| 87 | 14[143] | 13[159] | GCAAACTCTTCAAAGCGAACCAGCCGCGCCC |
| 88 | 14[175] | 12[176] | TTATTTTCGCAAATCAGATATAGAATTTGCCA |
| 89 | 14[207] | 12[208] | TACCGCACCTAAGAACGCGAGGCCCAACGC |
| 90 | 14[239] | 12[240] | TTATCATTCGACTTGCGGGAGGTTTGCACCCA |
| 91 | 15[64] | 17[63] | CAACTAAACTACTAATAGTAGTAGAAAGAATT |
| 92 | 15[96] | 17[95] | CCATATAATGGGGCGCGAGCTGAACAGAGCAT |
| 93 | 15[128] | 17[127] | GAGTAGATTGGTCAATAACCTGTTACATTATG |
| 94 | 15[160] | 16[144] | AACAACATAATTCTGTCCAGACGAGATACAT |
| **95** | **15[192]** | **17[191]** | **GTTTATCAAGAGAATATAAAGTACAAAAGCCT** |
| 96 | 15[224] | 17[223] | AAAAATAATTTAGGCAGAGGCATTAAATTCTT |
| 97 | 16[79] | 14[80] | CATCAATTGTACGGTGTCTGGAAGGTCATT |
| 98 | 16[111] | 14[112] | TTTTCATTCAGTTGATTCCCAATTGAGAGTAC |
| 99 | 16[143] | 15[159] | TTCGCAAATTAGTTTGACCATTACGACAATA |
| 100 | 16[175] | 14[176] | GGTAAAGTGTTCAGCTAATGCAGAAGCCGTTT |
| 101 | 16[207] | 14[208] | CAGTAATAACAATAGATAAGTCCAACCAAG |
| 102 | 16[239] | 14[240] | ACATGTAATATCCCATCCTAATTTGTCTTTCC |
| 103 | 17[64] | 19[63] | AGCAAAATGTAAAGATTCAAAAGGCAATATGA |
| 104 | 17[96] | 19[95] | AAAGCTAAATTTTAAATGCAATGCATTAATGC |
| **105** | **17[128]** | **19[127]** | **ACCCTGTACGCAAGGATAAAAATTGATCTACA** |
| 106 | 17[160] | 18[144] | AACACCGGTAAATAAGGCGTTAAAAAGCCTT |
| **107** | **17[192]** | **19[191]** | **GTTTAGTAAAATTTAATGGTTTGAGGTCTGAG** |
| 108 | 17[224] | 19[223] | ACCAGTATATATATTTTAGTTAATTTAGGTTG |

| | | | |
|---|---|---|---|
| 109 | 18[79] | 16[80] | ATGTGTAGTAAGCAATAAAGCCTAAGGTGG |
| 110 | 18[111] | 16[112] | CCTCATATATCGGTTGTACCAAAATAGCTATA |
| 111 | 18[143] | 17[159] | TATTTCAAATACTTTTGCGGGAGTAAGAATA |
| 112 | 18[175] | 16[176] | CCGTGTGAAATCATAATTACTAGACGACAAAA |
| 113 | 18[207] | 16[208] | TCTGACCTTCATATGCGTTATACTTCGAGC |
| 114 | 18[239] | 16[240] | TTTTTCAAAAAGCCAACGCTCAACCAACGCCA |
| 115 | 19[64] | 21[63] | TATTCAACTCAGAAAAGCCCCAAATGTAAACG |
| 116 | 19[96] | 21[95] | CGGAGAGGGCATGTCAATCATATGTTAAATTT |
| 117 | 19[128] | 21[127] | AAGGCTATAAGAGAATCGATGAACCCAATAGG |
| 118 | 19[160] | 20[144] | AATAGTGATAGATTAAGACGCTGAGAGTCTG |
| **119** | **19[192]** | **21[191]** | **AGACTACCCCCTTAGAATCCTTGAGATGAAAC** |
| 120 | 19[224] | 21[223] | GGTTATATCTTCTGTAAATCGTCGTTACATTT |
| 121 | 20[79] | 18[80] | GTTGATAACGTTCTAGCTGATAACTGAGTA |
| **122** | **20[111]** | **18[112]** | **TAAAACTAGTAGCTATTTTTGAGATTTAGAAC** |
| 123 | 20[143] | 19[159] | GAGCAAACCAGGTCATTGCCTGAGAAGAGTC |
| 124 | 20[175] | 18[176] | CGATAGCTATTTATCAAAATCATAAATACCGA |
| 125 | 20[207] | 18[208] | TTAATTTTTTTTTAACCTCCGGCTTCATCT |
| 126 | 20[239] | 18[240] | TAACCTTGAACTATATGTAAATGCGAGAAAAC |
| 127 | 21[64] | 23[63] | TTAATATTACGGCGGATTGACCGTCATCGTAA |
| 128 | 21[96] | 23[95] | TTGTTAAATAACAACCCGTCGGATGACGACGA |
| 129 | 21[128] | 23[127] | AACGCCATGCCAGCTTTCATCAACACTCCAGC |
| 130 | 21[160] | 22[144] | TCAATTACTCGCGCAGAGGCGAATTCTGGCC |
| 131 | 21[192] | 23[191] | AAACATCACGCCTGATTGCTTTGAATAATGGA |
| 132 | 21[224] | 23[223] | AACAATTTCCTTTTACATCGGGAGAATTATTT |
| 133 | 22[79] | 20[80] | GGGAACAATTGTTAAAATTCGCATACCCCG |
| 134 | 22[111] | 20[112] | TGAGCGAGTCAGCTCATTTTTTAAGGTAATCG |
| 135 | 22[143] | 21[159] | TTCCTGTACAAAAATAATTCGCGTATTCATT |
| 136 | 22[175] | 20[176] | TTACAAAACTGAGCAAAAGAAGATAAACATAG |
| 137 | 22[207] | 20[208] | AACGGATTAGAAAACAAAATTAACTATTAA |
| 138 | 22[239] | 20[240] | TAACAGTACATTTGAATTACCTTTTGAGTGAA |
| 139 | 23[64] | 25[63] | CCGTGCATGCTATTACGCCAGCTGTTGGGTAA |
| 140 | 23[96] | 25[95] | CAGTATCGGTTGGGAAGGGCGATCCGTTGTAA |
| 141 | 23[128] | 25[127] | CAGCTTTCAAAGCGCCATTCGCCAATGCCTGC |
| 142 | 23[160] | 24[144] | TGATTGTTGATGGCAATTCATCAATGCCGGA |
| 143 | 23[192] | 25[191] | AGGGTTAGCGGAATTATCATCATAGATAATAC |
| 144 | 23[224] | 25[223] | GCACGTAATTTTGCGGAACAAAGAACTTTACA |
| 145 | 24[79] | 22[80] | GCCTCTTCCTGCCAGTTTGAGGGTCTCCGT |
| 146 | 24[111] | 22[112] | GCGCAACTGCCTCAGGAAGATCGCATTAAATG |
| 147 | 24[143] | 23[159] | AACCAGGCCGGCACCGCTTCTGGTATAATCC |
| 148 | 24[175] | 22[176] | TATCAGATTGGATTATACTTCTGAATACCAAG |
| 149 | 24[207] | 22[208] | AGAAGGAGAACCTACCATATCAAAAACAAT |
| 150 | 24[239] | 22[240] | CATTATCAAACAGAAATAAAGAAAATATACAG |
| 151 | 25[64] | 27[63] | CGCCAGGGCATACGAGCCGGAAGCGAGCTAAC |
| 152 | 25[96] | 27[95] | AACGACGGGAAATTGTTATCCGCTCTGCCCGC |
| 153 | 25[128] | 27[127] | AGGTCGACTTCGTAATCATGGTCACCAGCTGC |
| 154 | 25[160] | 26[144] | ACTAATAGAAAATATCTTTAGGAGGGTACCG |
| 155 | 25[192] | 27[191] | ATTTGAGGCAACAGTTGAAAGGAAAAAACAGA |
| 156 | 25[224] | 27[223] | AACAATTCCCCTCAATCAATATCTCGCCTGCA |
| 157 | 26[79] | 24[80] | CCACACAATTTTCCCAGTCACGAGGTGCGG |
| 158 | 26[111] | 24[112] | TCCTGTGTCCAGTGCCAAGCTTGCTTCAGGCT |
| 159 | 26[143] | 25[159] | AGCTCGAATCTAGAGGATCCCCGCACTAACA |
| 160 | 26[175] | 24[176] | GGTTATCTATTAGAGCCGTCAATATTCCTGAT |
| 161 | 26[207] | 24[208] | TGGCAAATATTTAGAAGTATTAGAACCACC |
| 162 | 26[239] | 24[240] | ATATCAAAGACAACTCGTATTAAATTGAGTAA |
| 163 | 27[64] | 29[63] | TCACATTACACCGCCTGGCCCTGACCCCAGCA |
| 164 | 27[96] | 29[95] | TTTCCAGTCCAGTGAGACGGGCAAGTTCCGAA |
| 165 | 27[128] | 29[127] | ATTAATGACGTATTGGGCGCCAGGAAAGAATA |
| 166 | 27[160] | 28[144] | CCGAACGACCTAAAACATCGCCATGGGAGAG |

| | | | |
|---|---|---|---|
| 167 | 27[192] | 29[191] | GGTGAGGCGGCTATTAGTCTTTAATCAATCGT |
| 168 | 27[224] | 29[223] | ACAGTGCCAGAATACGTGGCACAGCAGATTCA |
| 169 | 28[79] | 26[80] | TTGCCCTTATTGCGTTGCGCTCACACAATT |
| 170 | 28[111] | 26[112] | TCTTTTCACGGGAAACCTGTCGTGTAGCTGTT |
| 171 | 28[143] | 27[159] | GCGGTTTGATCGGCCAACGCGCGTAAAAATA |
| 172 | 28[175] | 26[176] | CTGATAGCACCACCAGCAGAAGATTTGAGGAA |
| 173 | 28[207] | 26[208] | TTTTGAATGGTCAGTATTAACACGGTCAGT |
| 174 | 28[239] | 26[240] | AAAGCGTAACGCTGAGAGCCAGCAAACCTCAA |
| 175 | 29[64] | 31[63] | GGCGAAAAATGGCCCACTACGTGAGGTGCCGT |
| 176 | 29[96] | 31[95] | ATCGGCAAGTCAAAGGGCGAAAAAAGGGAGCC |
| 177 | 29[128] | 31[127] | GCCCGAGAAGAGTCCACTATTAAAAGCCGGCG |
| 178 | 29[160] | 30[144] | ATGGAAATGCCATTGCAACAGGAATTCCAGT |
| 179 | 29[192] | 31[191] | CTGAAATGTGCTGGTAATATCCAGGAATCCTG |
| 180 | 29[224] | 31[223] | CCAGTCACGCCTGAGTAGAAGAACGGCCACCG |
| 181 | 30[79] | 28[80] | TCAGGGCGTCCTGTTTGATGGTGCAGCTGA |
| 182 | 30[111] | 28[112] | ACTCCAACAATCCCTTATAAATCAGTGGTTTT |
| 183 | 30[143] | 29[159] | TTGGAACATAGGGTTGAGTGTTGAAACGCTC |
| 184 | 30[175] | 28[176] | TACCGCCAACCTACATTTTGACGCTGCGCGAA |
| 185 | 30[207] | 28[208] | ATCGGCCTGATTATTTACATTGGACAATAT |
| 186 | 30[239] | 28[240] | CATCACTTACGACCAGTAATAAAACTGACCTG |
| 187 | 31[64] | 30[80] | AAAGCACTAAATCGGAACCCTAACCGTCTA |
| 188 | 31[96] | 30[112] | CCCGATTTAGAGCTTGACGGGGAAGAACGTGG |
| 189 | 31[128] | 31[159] | AACGTGGCGAGAAAGGAAGGGAATTAAAGGG |
| 190 | 31[160] | 30[176] | ATTTTAGACAGGAACGGTACGCCAAACAATAT |
| 191 | 31[192] | 30[208] | AGAAGTGTTTTTATAATCAGTGATCAAACT |
| 192 | 31[224] | 30[240] | AGTAAAAGAGTCTGTCCATCACGCAGTAATAA |
